# Environmental and genetic drivers differentially shaped resident and transient gut microbiota during the Macaronesian chaffinch radiation

**DOI:** 10.1101/2025.08.20.671302

**Authors:** B. Karina Montero, Mark A.F. Gillingham, Mark Ravinet, Juan Carlos Illera

**Affiliations:** Biodiversity Research Institute, Consejo Superior de Investigaciones Científicas (CSIC) and Oviedo University–Principality of Asturias, University of Oviedo, Campus of Mieres, Mieres E-33600, Spain; Centre for the Synthesis and Analysis of Biodiversity, Foundation for Research on Biodiversity, Montpellier 34000, France; Center for Evolutionary and Functional Ecology (CEFE), Centre National de la Recherche Scientifique (CNRS), Montpellier 34000, France; Department of Ornithology, Max Planck Institute for Biological Intelligence, Eberhard Gwinner Straße, 82319 Seewiesen, Germany; Centre for Ecological and Evolutionary Synthesis, Department of Biosciences, University of Oslo, Oslo 0316, Norway

**Keywords:** avian gut microbiome, co-diversification, phylosymbiosis, *Fringilla*, environmental filtering, island biogeography theory, diet metabarcoding, host genetic control, inbreeding

## Abstract

Disentangling whether gut microbiome diversification reflects host evolutionary history or spatial processes remains challenging because these factors are often confounded. The Macaronesian chaffinch (*Fringilla*) radiation provides a replicated natural experiment to separate these effects. Using long-read 16S sequencing, diet metabarcoding, and whole-genome data, we evaluated competing models of community assembly, testing host-mediated dispersal versus neutral assembly, and environmental filtering versus host control. We show that microbiome alpha diversity across host species was largely shaped by island size and isolation, aligning with Island Biogeography Theory, rather than host colonisation history. This indicates that microbial dispersal occurs independently of their hosts. Moreover, spatial and host factors influenced different aspects of the microbiome: geography and diet primarily predicted which taxa were present, whereas host genetic relatedness and heterozygosity were associated with the abundance of established residents. Notably, we show that inbreeding compromises a host’s ability to regulate microbial abundance, leading to highly variable microbiomes. While we detected phylosymbiosis (closely related hosts harboring similar microbiomes), we found no evidence of co-diversification between birds and microbes. Instead, associations likely arise from convergent ecological filtering, where shared host traits filter similar bacteria from the local environment. Together, our results show that avian microbiomes are shaped by distinct spatial and host-mediated processes, and that separating microbial presence from abundance is important for understanding the mechanisms driving microbiome assembly.

## Introduction

The host microbiome underpins a wide range of biological functions, from improving nutrient uptake [1–3] to modulating behaviour [4, 5], profoundly impacting host health [6–8] and fitness [9]. These roles suggest microbiomes may facilitate adaptation to novel environments [10–12], with composition reflected in proxies of habitat use, such as diet [13]. Over host evolutionary history, reciprocal host-microbe interactions can generate patterns whereby microbiome composition mirrors host phylogeny [14–18], a phenomenon known as phylosymbiosis [19]. While phylosymbiosis may reflect co-diversification through mutual selective forces [20], it can also arise via environmental filtering, by which local abiotic and biotic conditions favour certain microbial taxa, through shared ecological niches or through parallel selective pressures [21–23]. Alternatively, neutral processes, such as stochastic dispersal and colonisation-extinction dynamics, can also structure microbiomes independently of host control, as predicted by island biogeography theory [24] and unified neutral models [25, 26]. Ultimately, stochasticity and host-mediated selection interact to shape microbiome structure [27, 28]; a central challenge is to disentangle the relative contributions of these mechanisms.

Here, we develop a framework to disentangle spatial and evolutionary drivers of microbiome diversification, testing four mechanisms with distinct predictions (Figure 1): (1) host-mediated dispersal (colonisation history structures microbiomes through founder effects); (2) host-independent dispersal (biogeographic factors predict diversity); (3) host control and environmental filtering shape distinct components of microbiome composition (genetics structures abundant taxa, environment shapes rare taxa); and (4) co-diversification (phylogenetic congruence among host-selected microbial lineages). We test these mechanisms using natural bird populations of the genus *Fringilla* across the Macaronesian archipelagos.

**Figure 1:**
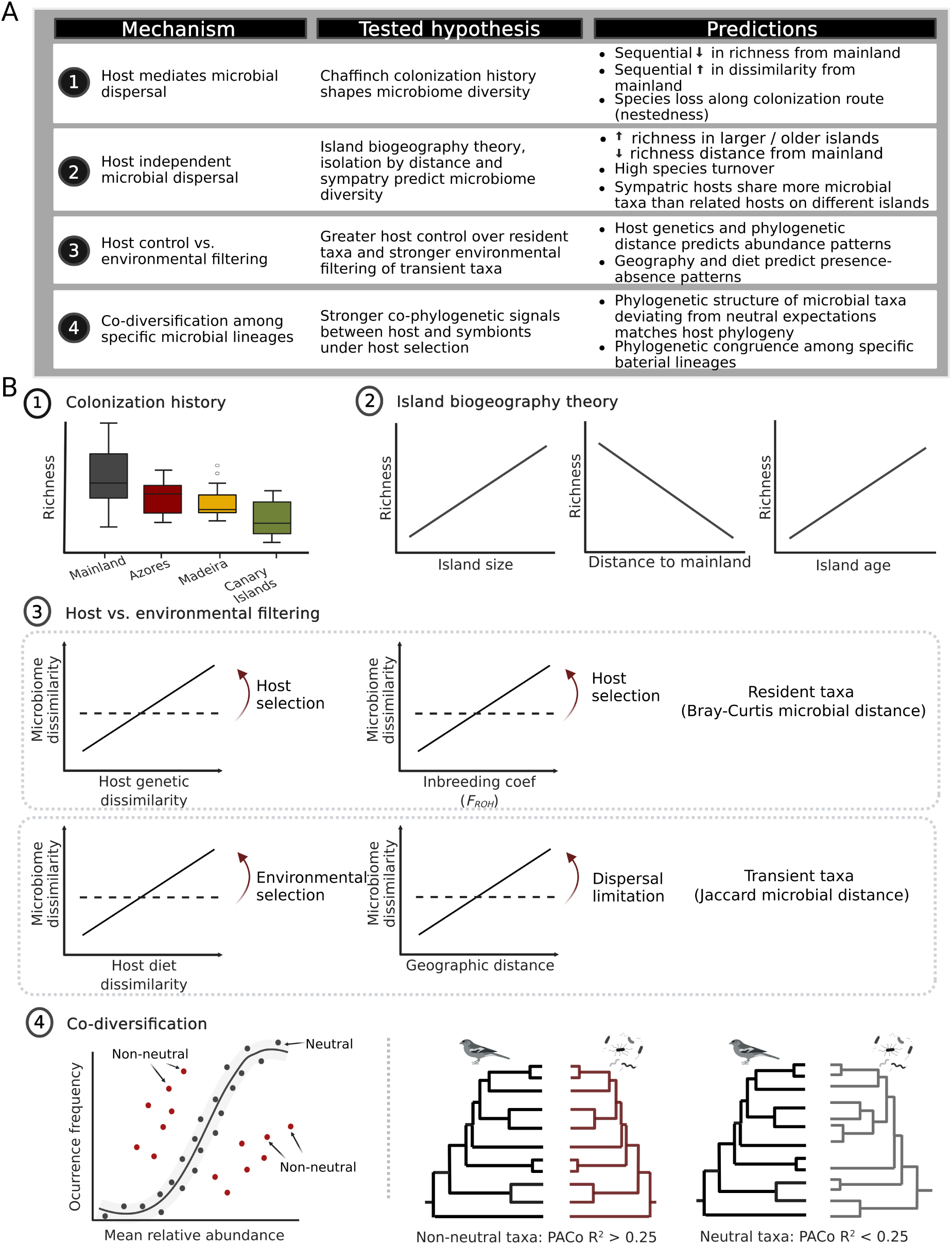
Framework for testing mechanisms of avian microbiome assembly and diversification. **A**. Summary of four mechanisms, hypotheses, and key predictions tested in this study. Mechanisms 1–2 evaluate spatial processes distinguishing host-mediated from host-independent microbial dispersal. Mechanism 3 assesses the relative contributions of host genetic control and environmental filtering to microbiome composition. Mechanism 4 tests for host-microbiome co-diversification through co-phylogenetic signal between hosts and microbial **B**. Main predictions for each of the four mechanisms. Mechanism 1: Host-mediated dispersal predicts a sequential decline in microbial richness along the host’s colonisation route. Mechanism 2: Host-independent dispersal predicts island biogeography patterns (island area, distance to mainland, and island age). Mechanism 3: Host control predicts correlations with host genetics (top), while environmental filtering predicts correlations with diet and geography (bottom). Arrows indicate the ecological mechanisms at play. Mechanism 4: Co-diversification predicts phylogenetic congruence between host and microbe trees, particularly for taxa under selection (i.e., taxa deviating from neutral expectations). Figure created with Biorender.

*Fringilla* colonised the North Atlantic Macaronesian archipelagos in two independent events, giving rise to two blue chaffinch and three common chaffinches species [29–32] (Figure 2A). Blue chaffinches arrived in the Canary Islands about 3 mya [33] and are now restricted to the pine forests of Tenerife and Gran Canaria. In contrast, Macaronesian common chaffinches diverged from their mainland counterparts *∼* 0.8 mya [31] and occupy 15 islands across the Azores, Madeira, and Canary archipelagos, where they primarily inhabit ”Monteverde” forests and, to a lesser extent, mixed woodland habitats [34]. Notably, their colonisation followed a counter-intuitive route from the mainland to the remote Azores, then southward via stepping stone dispersal to Madeira and finally the Canary Islands [31, 32] (Figure 2B).

**Figure 2:**
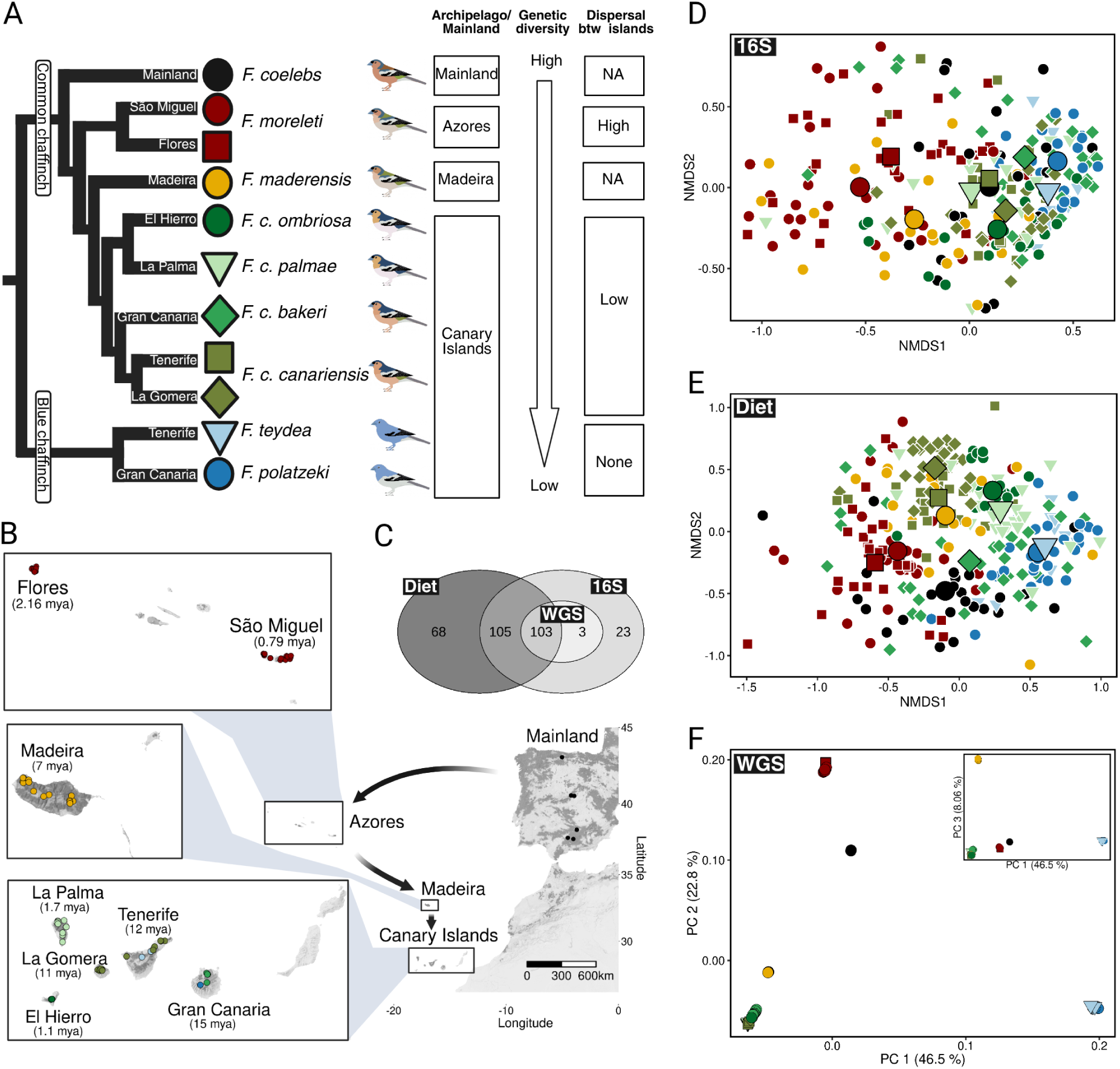
Overview of the study system and dataset. **A**. Phylogenetic relationships, geographic distributions, and genetic diversity patterns across Macaronesian finches (genus *Fringilla*).We studied six species: four common chaffinches and two blue chaffinches. Three subspecies are currently recognised for *F. canariensis*, the common chaffinch in the Canary Islands [59]. Genetic diversity follows a gradient consistent with stepwise colonisation: Azorean and mainland chaffinches maintain high genetic diversity and dispersal rates, *F. maderensis* and *F. canariensis* show intermediate reductions due to founder effects [31], while blue chaffinches exhibit the lowest diversity with no inter-island dispersal [60] (see Supplementary Table S1). **B**. colonisation route of common chaffinches across Macaronesia (black arrows) [31] and sampling locations. Numbers in brackets indicate island age (mya). Dot colors correspond to populations shown in **A**. The genus *Fringilla* is naturally absent from Fuerteventura and Lanzarote. **C**. Sample sizes for microbiome profiling (16S rRNA), diet metabarcoding (rbcL and COI markers), and whole-genome sequencing (WGS). **D–E**. Non-metric multidimensional scaling (NMDS) ordination based on Bray-Curtis distances for (**D**) gut microbiomes (species-level or lowest taxonomic assignment; *n* = 236) and Raup-Crick distances for (**E**) diet composition (family-level or lowest taxonomic assignment) from merged rbcL and COI marker data (*n* = 276). **F**. Host population genetic structure from principal component analysis of WGS data. Inset shows PC1 vs. PC3.

The recent radiation of chaffinches, combined with their ecological similarity across Macaronesia, offers a natural experiment for studying microbiome diversification mechanisms. Colonisation of new environments, such as oceanic islands, often triggers adaptive radiations driven by ecological opportunity [35–39], leading to pronounced phenotypic divergence [40–45]. However, many island radiations exhibit minimal ecological niche disparity [46], a pattern known as non-adaptive radiation, which typically occurs under limited geographic overlap [47, 48] and is mainly driven by neutral forces such as genetic drift [45, 49, 50]. Importantly, the limited niche disparity observed in insular non-adaptive radiations, as seen in Macaronesian finches [45], makes them particularly well suited for disentangling the relative contributions of spatial and evolutionary processes to diversification.

First, we assess whether hosts mediate microbial dispersal by examining whether chaffinch colonisation history shapes microbiome diversity. Under this scenario, we predict sequential loss of microbial richness along the colonisation route, increasing dissimilarity from mainland populations, and beta diversity dominated by nestedness, reflecting stepwise species loss analogous to host genetic founder effects (Figure 1B, Prediction 1). Second, we evaluate whether microbial dispersal occurs independently of host movement by testing predictions from Island Biogeography Theory [24, 51, 52] and isolation-by-distance [53]. Here, microbial richness should correlate with island area, age, and isolation, with beta diversity dominated by species turnover rather than nestedness (Figure 1B, Prediction 2). Third, we assess the relative contribution of host control and environmental filtering by testing whether host genetics preferentially structures established (abundant) taxa while environmental factors (geography, diet) primarily shape transient (rare) taxa. Specifically, we predict host genetics (phylogenetic distance, species and subspecies genetic differentiation, and estimates of inbreeding) correlates with abundance-weighted community patterns, while geography and diet predict presence-absence patterns (Figure 1B, Prediction 3). Fourth, we test whether co-diversification occurs among specific microbial lineages by examining whether symbionts under host control show stronger co-phylogenetic signals than neutral taxa. We predict that microbial lineages deviating from neutral expectations will exhibit phylogenetic structure that matches host phylogeny, indicating co-evolutionary partnerships rather than incidental associations (Figure 1B, Prediction 4). The relatively recent divergence of Macaronesian chaffinches (*∼* 0.8-3 mya) may limit our ability to detect co-diversification signals using the slow-evolving 16S rRNA marker. Nevertheless, this system offers an opportunity to address the ongoing debate over host genetic control of microbiome structure. This question has been widely investigated in mammals [14–17, 54–56], but remains contentious in birds, where transient microbial taxa are thought to dominate and host control is generally considered minimal [57, 58].

By integrating host genomics, long-read 16S rRNA microbiome profiling, and fine-grained dietary data within a well-characterised island radiation, we disentangle spatial and evolutionary drivers of gut microbiome structure across replicated island systems. Our findings clarify mechanisms underlying phylosymbiosis patterns and advance understanding of how host control and neutral assembly processes shape avian microbiomes in natural populations.

## Results

We sampled a total of 302 birds spanning 11 populations across Macaronesia and the Iberian mainland (12–36 individuals per population; Figure 2B; Supplementary Table S1 and S2). The six sampled *Fringilla* species comprised *F. coelebs* (mainland), *F. moreletti* (Azores), *F. maderensis*, and *F. canariensis* (with five named subspecies across Madeira and the Canary Islands), and the Canary Island endemics *F. polatzeki* and *F. teydea* (Figure 2A). From each bird, we collected faeces for microbiome taxonomic profiling using synthetic long-read (SLR) full-length sequencing of the 16S rRNA gene and diet characterisation through rbcL and COI amplicon sequencing. After quality filtering, 234 samples were retained for the microbiome dataset (9–28 per population) and 276 for the diet dataset (12–34 per population), with both data types available for 208 individuals (9–26 per population; Figure 2C; Supplementary Table S1). We analysed bacterial features from phylum to species level, and to investigate diversity at finer taxonomic scales, we used operational taxonomic units (OTUs) clustered at 99% similarity. Unless otherwise specified, we report results based on species (but see Supplementary Materials for results at higher taxonomic ranks and at the OTU level).

Microbiome composition differed strongly among finch species and their populations across all four distance metrics tested (PERMANOVA, Bray-Curtis: *R*^2^ = 0.16, *p <*0.001; Jaccard: *R*^2^ = 0.10, p *<*0.001; weighted UniFrac: *R*^2^ = 0.19, *p <*0.001; unweighted UniFrac: *R*^2^ = 0.12, *p <*0.001;Figure 2D; Supplementary Table S3). Diet composition was likewise structured by species and population (PERMANOVA, Raup-Crick: *R*^2^ = 0.51, *p <*0.001; Figure 2E).

In contrast, neither age nor sex was associated with microbiome composition or alpha diversity (PERMANOVA on composition: all *R*^2^ *≤* 0.005, all *p >*0.17; Supplementary Table S3; GLMMs on richness and Faith’s PD: all *p <*0.14). Diet composition was similarly unaffected by age or sex (PERMANOVA, both *R*^2^*<*0.004, *p >*0.18). We therefore pooled samples across these demographic categories for subsequent analyses.

We conducted whole-genome-sequencing (WGS) on blood samples from a subset of individuals (106 individuals, Figure 2C; Supplementary Table S1), generating a dataset of *∼* 51.6 million genotyped SNPs. All three data types (microbiome, diet, and WGS) were available for 103 individuals (2–11 per population; Figure 2C; Supplementary Table S1). Genetic diversity of Macaronesian chaffinches exhibits a gradient consistent with founder effects during sequential island colonisation events (e.g. [61]). Blue chaffinches, in particular, show high levels of inbreeding and low genetic diversity ([30], Supplementary Table S1). Principal component analysis (PCA) of the WGS data and a maximum-likelihood phylogeny built from mitochondrial consensus genomes recapitulated previously described patterns of population structure and evolutionary relationships among Macaronesian finches [31] (Figure 2F).

### Island biogeography but not colonisation history explains microbiome diversity

Microbiome diversity patterns contradicted predictions from host-mediated dispersal (Mechanism 1, Figure 1). Most island populations exhibited microbial richness similar to or greater than mainland finches (Figure 3A), a pattern that held when accounting for bacterial phylogenetic diversity (Faith’s PD; Supplementary Table S4 and Figure S1A). In fact, two of five populations of Canary Island chaffinches and both blue chaffinch species showed higher microbial richness than mainland populations (Figure 3A; Supplementary Table S4 and Figure S1A). Variation in microbial richness between sites was not explained by diet richness (GLMM with negative binomial distribution; richness: *β* [95% *CI*] = 0.040 [-0.047, 0.128]; *p* = 0.367; Faith’s PD: *β* [95% *CI*] = 0.035 [-0.037, 0.107]; *p* = 0.340). Dissimilarity from mainland was high across all populations with no sequential increase along the colonisation route (Figure 3B; UniFrac: Supplementary Figure S1 B-C). Beta diversity was dominated by turnover (90.01% for Bray-Curtis, 84.71% for Jaccard; Figure 3C, Supplementary Figures S2 and S3), rather than the nestedness expected under sequential founder effects, indicating that hosts acquire microbes primarily from local environments rather than through vertical transmission.

**Figure 3:**
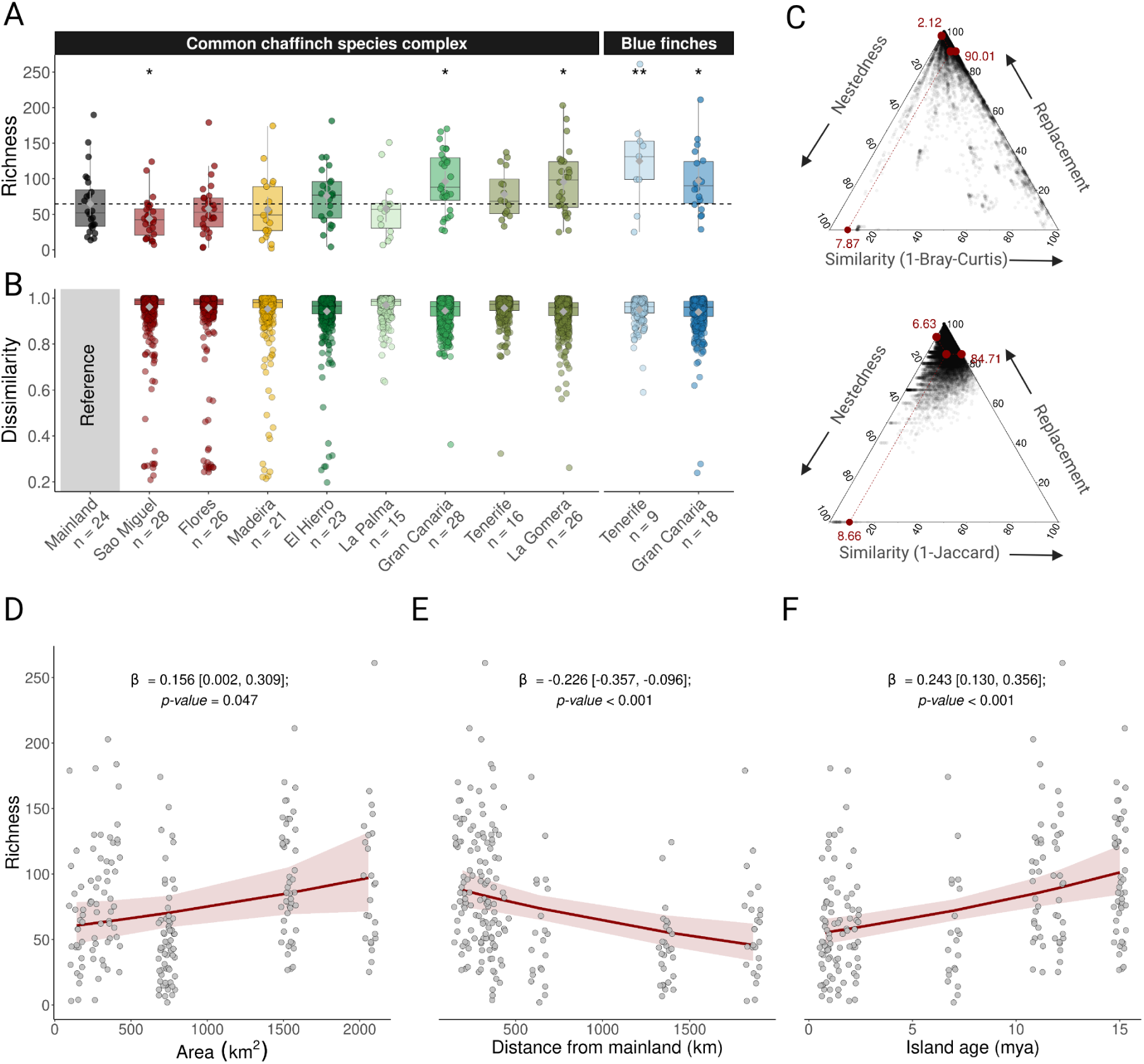
Microbiome diversity patterns according to colonisation history and biogeography. **A**. Species-level microbial richness across populations of the common chaffinch complex and the two blue chaffinch species. For the common chaffinch, populations are ordered along the colonisation route: mainland, Azores, Madeira, and Canary Islands. Asterisks indicate populations whose microbiome richness differs significantly from the mainland reference (negative binomial GLM; *P *<*0.05, **P *<*0.01). **B**. Microbiome dissimilarity (Bray-Curtis) between mainland and each Macaronesian population, computed from pairwise distances between every individual on each island and every mainland individual. Box plots show the distribution of pairwise dissimilarities; grey diamonds with whiskers indicate bootstrapped means and 95% confidence intervals. Mainland serves as the reference and is therefore left blank. **C**. Ternary plots showing the additive components of beta diversity, computed from abundance data (Bray-Curtis, top) and presence-absence data (Jaccard, bottom). Each black dot corresponds to a pair of common chaffinch individuals from different populations; coordinates reflect the relative contributions of richness difference, replacement (turnover), and similarity. Red dots at the plot edges indicate the mean values of these components, and the central red dot marks the centroid. **D-F**. Relationships between microbiome richness and **(D)** island area, **(E)** distance from mainland, and **(F)** island age (negative binomial GLMMs with population as a random effect). Sample sizes: n = 234 individuals for panels **A** and **B**; n = 207 (common chaffinch complex only) for panel **C**; n = 210 (Macaronesian populations only) for panels **D–F**.

We found that sympatric species on Gran Canaria and Tenerife, which occur in the same islands but have different evolutionary histories (Figure 2), harboured more similar microbiomes (13.1% shared species) than allopatric populations of the same species complex on different islands (11.2% shared; *p <*0.05, all comparisons consistent using UniFrac distances; Supplementary Figure S4), confirming the dominant role of geography over host evolutionary history in shaping microbial communities.

In support of island biogeography predictions (Mechanism 2, Figure 1), microbial richness increased with island area and age, and decreased with distance to the mainland (Figure 3 D-F, Supplementary Table S5), and these patterns held when accounting for bacterial phylogeny (Supplementary Figure S1 D-F). Microbiome dissimilarity increased significantly with geographic distance (Mantel tests: Bray–Curtis *r* = 0.16, Jaccard *r* = 0.37, weighted UniFrac *r* = 0.13, unweighted UniFrac *r* = 0.29, p *≤* 0.001 for all metrics; Supplementary Table S6), and this relationship persisted after controlling for host phylogenetic distance (partial Mantel tests: Bray-Curtis *r* = 0.12, Jaccard *r* = 0.36, weighted UniFrac *r* = 0.12, unweighted UniFrac *r* = 0.27, p *<*0.005 for all metrics). Together, these results indicate geographic distance shape microbiome structure independently of host evolutionary history, consistent with isolation-by-distance and host-independent dispersal limitation.

### Host genetic effects are stronger for abundant than rare microbial taxa

Our data support the hypothesis that host genetics plays a prominent role in shaping abundant taxa, while the composition of rare taxa responds primarily to diet and geography (Mechanism 3, Figure 1). To test this, we assessed three complementary aspects of host genetic variation: mitochondrial phylogenetic distance (shared host ancestry), nuclear *F_ST_* (population-level genomic differentiation), and nuclear *F_ROH_* (individual-level inbreeding and heterozygosity loss). Multiple regression on distance matrices (MRM) examining the relative contribution of host phylogenetic distance, diet, and geography revealed significant signals of phylosymbiosis (i.e., similar microbiome composition among closely related hosts), but showed that the taxonomic scale at which host phylogeny predicted microbiome dissimilarity differed between metrics (Figure 4A; Supplementary Table S7). For abundance-based patterns, phylogeny was a significant predictor at finer taxonomic scales (family-OTU: *R*^2^ = 0.035 - 0.045, *p*-value *<*0.001; Supplementary Table S7). In contrast, for presence-absence patterns, phylogeny predicted dissimilarity at broad taxonomic levels (order-phylum: *R*^2^ = 0.015 - 0.042, *p*-value *<*0.05; Supplementary Table S7) and at the OTU level (*R*^2^ = 0.038, *p*-value = 0.038). This contrast suggests that host genetics influences which higher-rank lineages are present in the gut, while at finer resolution it shapes the relative abundances of established resident microbiota. Geography and diet were the dominant predictors of occupancy patterns (Figure 4A; Supplementary Table S7). However, all models left a large fraction of unexplained variation at all taxonomic levels (*≥*79% for Jaccard; *≥*93% for Bray-Curtis), consistent with high turnover of rare, transient taxa typical of avian guts.

**Figure 4:**
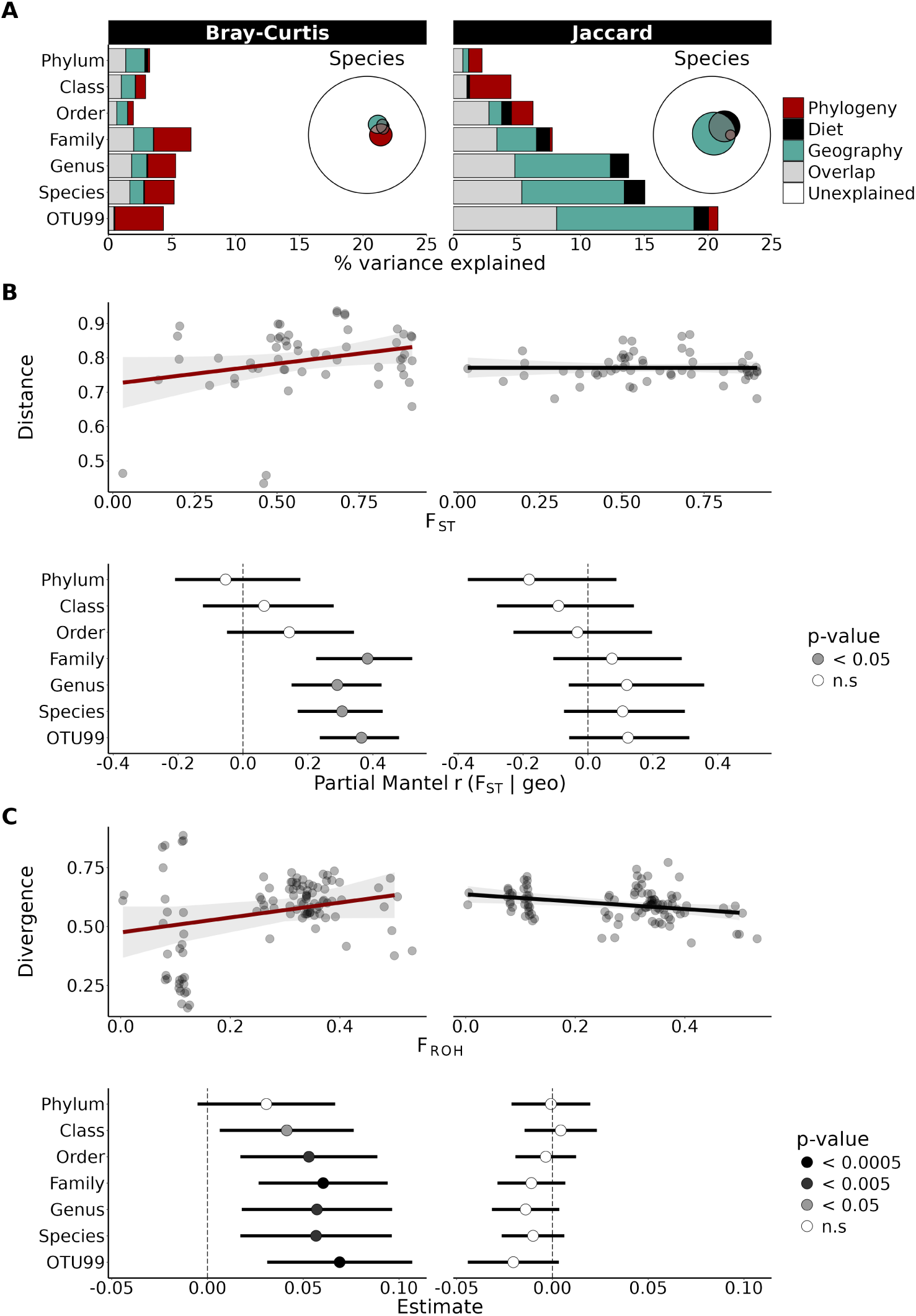
Host genetics impacts microbiome abundance but not occupancy patterns. **A**. Variance partitioning of microbiome dissimilarity decomposed into contributions of host mitogenome phylogeny (cophenetic distance), diet (Raup-Crick dissimilarity), and geography (log-transformed pairwise geographic distance), using multiple regression on distance matrices (MRM) with two microbiome metrics: Bray-Curtis (left; abundance-based) and Jaccard (right; presence–absence). Bars show the unique and shared contributions of each predictor across taxonomic ranks; Euler diagrams summarise the species-level partitioning. **B** Top: relationship between population genetic differentiation (*F_ST_* )and population-level microbiome dissimilarity at the species level, with each point the median pairwise Bray-Curtis or Jaccard distance across 1000 random subsamples of 9 individuals per population. Bottom: median partial Mantel *r* (controlling for log-transformed geographic distance) across the same 1000 subsamples, with 2.5–97.5% percentile intervals, across taxonomic ranks. Fill colour indicates the median permutation *p*-value. **C**. Effect of host inbreeding on microbiome stability. Top: relationship between individual inbreeding coefficient (*F_ROH_*) and microbiome among-individual divergence (distance-to-median in *β*-diversity) at the species level. Bottom: GLMM effect sizes (*β ±* 95% CI) for *F_ROH_* on among-individual divergence across taxonomic ranks, with fill colour indicating *p*-value. Sample sizes: n = 103 individuals for all panels.

Population genetic differentiation (*F_ST_* ) confirmed the stronger effects of host genetics on resident taxa, correlating with abundance-based dissimilarity (family-OTU: median partial Mantel *r* = 0.29 - 0.38, median *p*-value *<*0.05) but not presence-absence data (Jaccard: median partial Mantel *r* between –0.18 and 0.12, median *p*-value *>*0.2 at all taxonomic levels; Figure 4B). Similarly, reduced heterozygosity (*F_ROH_*) increased among-individual microbiome variation only for abundance metrics (Figure 4C), indicating that inbreeding weakens host control over microbial abundances rather than colonisation potential. Together, these findings demonstrate that host control acts primarily on the resident component of the microbiome, while environmental filtering shape the transient fraction.

### Evidence of co-phylogeny but not co-diversification

We next aimed to distinguish between co-phylogenetic signal (closely related hosts harbouring related microbes) [62–64] and true co-diversification (synchronised host-microbe speciation) [62, 63]. To this end, we tested the nature of host-microbe associations with three complementary approaches. First, we evaluated community-wide (global) patterns by including the entire microbial community, comparing neutral versus non-neutral (putatively host-selected) subsets to determine whether taxa under host control show stronger phylogenetic associations. Second, we identified specific bacterial families with significant co-phylogenetic signals and applied event-based reconciliation methods to test for phylogenetic congruence (i.e. matching branching patterns and divergence timing), required to infer co-diversification [62, 63]. Third, we directly compared host and microbial phylogenies to evaluate whether microbial diversification corresponds to host evolutionary events or instead reflects geographic structuring and environmental acquisition. Together, these analyses revealed weak associations between host and microbe phylogenies but no evidence of co-diversification (Mechanism 4, Figure 1).

### Weak co-phylogenetic signals across global and neutrality-partitioned patterns

To identify taxa that deviates from neutrality, we applied the prokaryote neutral model (PNM) [65–67]. Model fit was poor at fine scales (OTU:R2 = -0.18) but improved at higher taxonomic levels (R2 = 0.31–0.79, species to phylum; Figure 5A; Supplementary Figure S5), indicating that most fine-scale microbial variants are unique to individual hosts with limited community equilibrium and suggesting that short-read studies analysing coarser taxonomic scales may overestimate neutral structuring.

**Figure 5:**
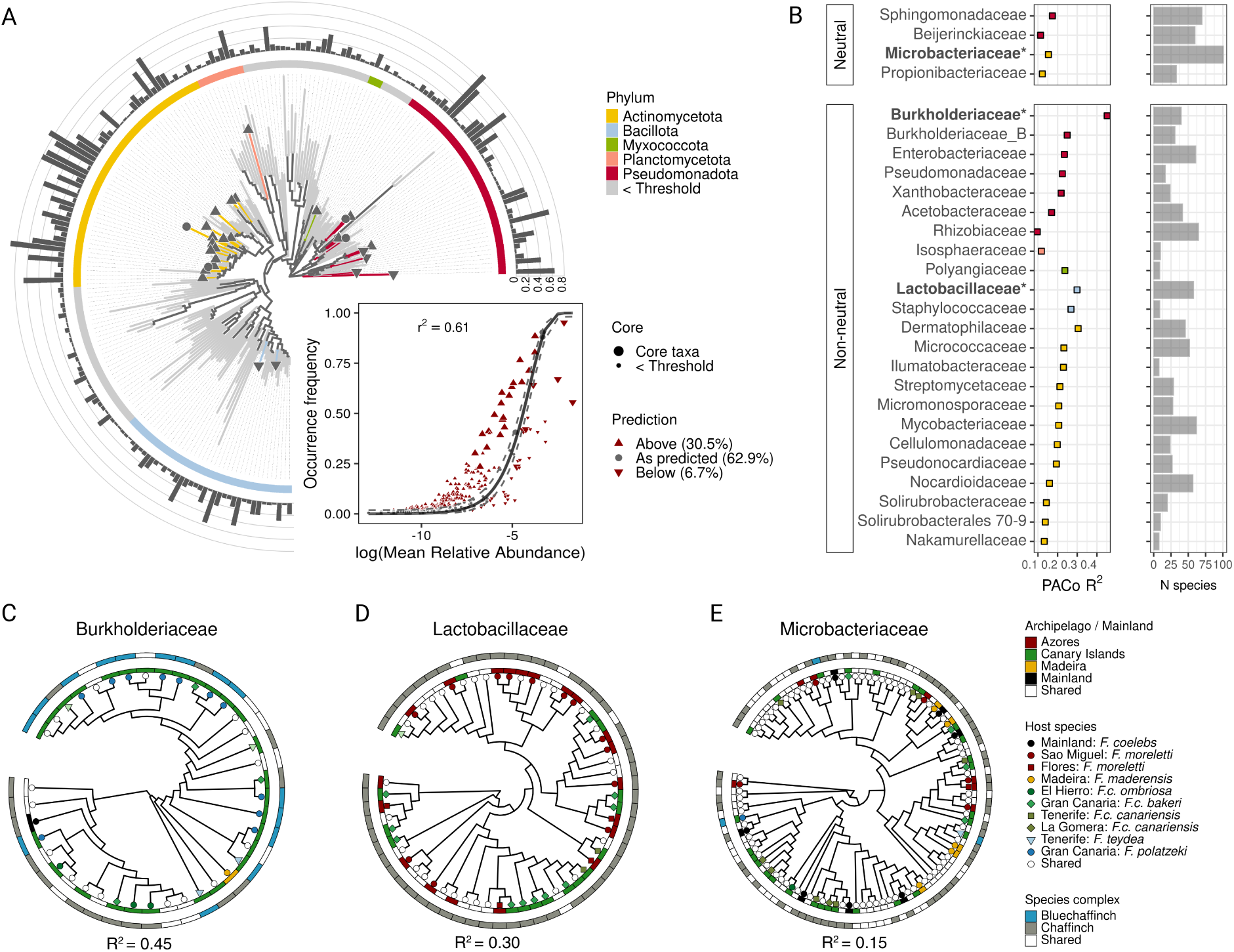
Evidence of host-microbiome co-phylogeny but not of co-diversification. **A**. Phylogenetic relationships of prevalent microbial families found in at least two individuals per population. Tip symbols and branch colors denote core families present on all islands. Symbols indicate whether each family’s abundance deviates above (upward triangle), below (downward triangle), or fits neutral expectations according to the procaryote neutral model (PNM) at the family level. The inner ring displays phylum assignments, and the outer ring shows prevalence of families across all samples. The inset shows the PNM model fit at the family level. **B**. PACo goodness of fit estimates (*R*^2^) for 27 core families with at least five microbial species (details in Methods). Bold family labels mark significant co-phylogeny signals after Bonferroni correction (*p <*0.05). **C-E**. Phylogenies of microbial lineages with significant PACo signals. Tip symbols indicate host populations. The inner and outer strips correspond to archipelago and species complex origins, respectively. Sample sizes: n = 208 individuals for panels **A** and **C - E**; n = 103 for panel **B**.

Contrary to predictions (Figure 1B, Prediction 4), we did not detect differences in the strength of co-phylogeny between non-neutral taxa (deviating either above or below predictions) and the neutral subset. PACo tests (Procrustean Approach to Cophylogeny) [68, 69], across all microbial taxa detected weak associations between host and microbe phylogenies at coarser taxonomic levels (genus-family: R2 = 0.07–0.12). However, at finer scales (OTU-species), yielded significant associations for both empirical and randomised host trees (R2 = 0.05-0.08), indicating potential false positives and suggesting that this approach is not reliable to assess co-phylogeny signals when the ratio of host–symbiont links is high, as observed in other systems [70, 71]. These weak global signals (R2 *<*0.25) suggest host control rather than co-diversification, which typically requires stronger cophylogenetic associations (R2 *>*0.25) and congruent branching patterns [62].

### Core families show co-phylogenetic signal but no phylogenetic congruence

Three of 27 core bacterial families (i.e. families found across all populations) showed significant co-phylogenetic signals (p *<*0.05): Burkholderiaceae (R2 = 0.45), Lactobacillaceae (R2 = 0.30), and Microbacteriaceae (R2 = 0.15) (Figure 5B). Within these family-specific datasets, randomised trees were not significant, confirming non-random associations (Supplementary Table S8). However, phylogenetic reconciliation using eMPRess [72], revealed high uncertainty (*>*1000 equally likely solutions) with no evidence of matching speciation events between hosts and microbes (co-speciation/transfer ratio *>*1, p *>*0.05; Supplementary Table S9), indicating that host-switching and microbial duplication dominate over co-speciation events.

### Microbial diversification reflects island geography, not host evolution

Direct comparison of host and microbe phylogenies revealed no matching splits corresponding to host speciation or colonisation events (Figures 5C–E). Instead, microbial diversification patterns followed island geography: Burkholderiaceae diversified predominantly within the Canary Islands, across both chaffinch and blue chaffinch lineages; Lactobacillaceae diversified within common chaffinches without tracking host phylogeny; and Microbacteriaceae, a species rich lineage, exhibited a lack of geographic or host structure, with numerous taxa shared across different archipelagos and host species complexes. These patterns support environmental acquisition followed by host control rather than co-diversification.

## Discussion

The non-adaptive radiation of chaffinch species in Macaronesia provided a natural experiment to test competing hypotheses of microbiome diversification that have proven challenging to disentangle [22, 23]. We provide pioneering evidence that host-microbiomes display diversity patterns consistent with island biogeography theory [24, 52, 73]. Microbiome community assembly was dominated by high microbial turnover, with geographic distance predicting dissimilarity even after controlling for host phylogeny. These results demonstrate that in isolated host populations, microbial dispersal limitation significantly shapes gut microbiomes independently of host evolutionary history.

Our results reveal a fundamental distinction in how environmental versus host factors influence different components of the microbiome. Strong associations between geography and gut microbiome structure, and to a lesser extent diet, were most pronounced in presence-absence (Jaccard) analyses, which are particularly sensitive to the detection of rare taxa. In contrast, abundance-based metrics (Bray-Curtis) highlighted more subtle, yet consistent, effects of host genetics. This suggests that presence-absence patterns capture variation of the environmentally acquired, transient fraction of the microbiome, whereas abundance-weighted analyses emphasize the established, potentially resident “core” gut microbiota [23].

Population genetic analyses reinforced this pattern: both host genetic differentiation (*F_ST_* ) and homozygosity (*F_ROH_*) correlated positively with abundance-based microbiome dissimilarity but showed no relationship with presence-absence patterns. Inbred finches, in particular vulnerable blue chaffinch populations with characteristically low genetic diversity, exhibited greater among-individual microbiome variation, but only for abundance-based metrics. These results extend findings from experimental systems like *Drosophila* [74] to an avian system, and suggests that reduced genetic diversity compromises host control over microbial abundance and divergence, likely through effects on immune gene diversity [75–79] and gut microbiome regulation [28, 80, 81], rather than affecting which taxa can initially colonise the gut. Presence–absence–based models explained more variance than abundance-weighted models (Fig. 4a), yet a substantial proportion of variation remained unexplained (*>*79%), consistent with the high interindividual variability typical of avian gut microbiomes, which often harbour large numbers of transient microbial taxa [16, 82, 83]. Such unexplained variance likely reflects a combination of individual-level stochasticity, the rapid turnover of transient taxa, and any potential temporal dynamics [82–86].

Our results also reinforce the notion that co-phylogenetic signals do not necessarily imply co-diversification [22, 23, 62]. While we detected significant co-phylogenetic patterns among select bacterial lineages, event-based analyses found no support for phylogenetic congruence, and host-microbe phylogenies lacked matching cladogenesis. One plausible explanation is the temporal mismatch resulting from the relative recent divergence of Macaronesian chaffinches (0.8-3 mya) [31] together with the slow evolutionary rate of the 16S rRNA gene (*∼* 1-2% per million years) [87, 88]. The weak co-phylogenetic signals we observed and high uncertainty in phylogenetic reconciliation are consistent with either ecological filtering of environmentally-acquired microbes through conserved host traits such as gut physiology and dietary preferences [89, 90] or undetectable recent co-diversification. Future metagenomic studies with higher resolution may be able to clarify whether these patterns are indeed masked by marker limitations [15, 88, 91]. Nonetheless, given our observed weak co-phylogenetic signals and stronger influence of biogeography over host phylogeny, we expect that even with improved resolution, only a restricted number of lineages, if any, would show true co-diversification patterns. Our study addresses the longstanding assumption that birds lack strong host-microbe associations due to adaptations linked to flight [16, 57, 58, 92]. Contrary to these predictions, we detect signals of phylosymbiosis in Macaronesian finches independent of geography and diet, and that primarily emerge in abundance-based microbiome patterns. These findings align with recent evidence for pervasive co-phylogenetic signals in birds [18] and reinforces the premise that host effects operate at different taxonomic scales [14, 16, 18].

In contrast to mammals, where strong signals of phylosymbiosis are often detected across diverse clades [17, 22, 54, 93, 94] , avian studies show weaker and more inconsistent patterns [16, 18, 58, 95–97]. Several biological differences likely contribute to this contrast. Mammals possess a unique suite of traits, including live birth, lactation, and a complex digestive system requiring strict immune control, that promote vertical transmission and strong host control [28, 98]. In contrast, early contact of eggs with the environment [85, 99, 100] and rapid gut transit times can lead to bird microbiomes being more sensitive to environmental acquisition and rapid turnover. However, our work suggests that the inconsistencies in bird studies and potentially more broadly across host-microbiome systems, where some find phylosymbiosis and others do not, can be attributed, at least in part, to variability in the methods used to study this question [101]. We used synthetic long-reads [102] to characterise the chaffinch microbial communities, which allowed us to examine the different taxonomic scales at which environmental and genetic factors operate, overcoming the limitations of short-reads often lacking fine scale resolution. Moreover, environmental filtering frequently dominate host genetic signals [103, 104] making it difficult to isolate patterns of phylosymbiosis. Resolving the drivers of phylosymbiosis requires tighter control of environmental exposure, including fine-scale dietary characterisation rather than coarse ecological proxies [54, 89, 105, 106], and replicated designs that capture spatial structure [107], as in the Macaronesian finch system. Equally important is high taxonomic resolution in microbial profiling to distinguish genuine phylogenetic signals from patterns generated by low signal-to-noise ratios.

In conclusion, our framework demonstrates that resolving mechanisms of microbiome assembly requires partitioning communities by taxonomic resolution and distinguishing patterns acting on transient taxa versus the resident components of the microbiome, as spatial and evolutionary processes act simultaneously but on different ecological components. The replicated, island radiation of Macaronesian finches challenges the assumption that host genetic control plays a minimal role in shaping gut microbiome associations in birds. Specifically, we show that microbial assembly is driven by a process of convergent filtering: related birds do not share a synchronised evolutionary history with their microbes, but they appear similar because their shared traits (like gut anatomy or diet) lead them to “filter” and pick up similar bacteria from the local island environments. By moving beyond descriptions of diversity, we show that island birds maintain host-specific microbiomes within biogeographic constraints that arise through these conserved ecological filters.

## Methods

### Sample collection

We captured birds between 2018 and 2021 using 2–4 mist nets (12 m each) aided by song playbacks. Individuals were marked with unique numbered metal rings authorised by Spanish or Portuguese authorities. We determined sex and age based on plumage coloration and molt patterns [108], with a few birds sexed molecularly [109]. To collect fecal samples, we placed each bird in a clean ringing bag for up to 10 minutes, transferring fecal material with sterile swabs into screw-cap microtubes containing 1 ml of NAP buffer [110]. All ringing bags were bleach-cleaned before field use and no bag was used twice within a field day. We collected approximately 50 µl of blood from the brachial vein, preserved in absolute ethanol at room temperature until transfer to –25 °C at Biodiversity Research Institute (IMIB) facilities.

### Microbial library preparation and processing

We isolated microbial DNA in a cleanroom at IMIB’s Ecological Molecular Laboratory (University of Oviedo, Mieres). DNA was extracted using the NucleoSpin Soil extraction kit (Macherey-Nagel). Library preparation and sequencing was carried out by Loop Genomics (now Element Biosciences) using LoopSeq kits (Loop Genomics) with primers 27F (5*^′^*-AGAGTTTGATCMTGGCTCAG-3*^′^*) and 1429R (5*^′^*-TACCTTGTTACGACTT-3*^′^*), and sequenced on the NovaSeq S4 platform following the methodology described by Callahan et al. [102]. The sequencing run included seven extraction blanks, two PCR blanks, and three field blanks. Field blanks consisted of sterile swabs run through clean (bleach-sterilised) ringing bags in the absence of a bird, controlling for contamination from the bag, swab, or buffer. None of the twelve blanks yielded long-read sequences. Synthetic long reads obtained from five sequencing runs totalled 18,237,257 reads, which we processed with the DADA2 (v1.32.0) pipeline [111] for long-read data in R v4.4.2 [112].

After primer removal, we applied quality filters (maxEE = 1, truncQ = 8, maxN = 0, minQ = 3) and trimmed sequences to lengths between 1400 and 1600 bp. We performed denoising separately for each run using run-specific error models optimised for NovaSeq data (https://github.com/benjjneb/dada2/issues/1307) and the pooled option in DADA2. We merged abundance tables for chimera removal and assigned taxonomy to amplicon sequence variants (ASVs) using the RDP Naive Bayesian Classifier [113] against the GTDB v4.4 database (https://zenodo.org/records/10403693). We excluded non-bacterial sequences and filtered ASVs with fewer than 10 total reads across all samples.

To standardize sequencing depth, we rarefied samples to 2000 sequences (threshold chosen based on rarefaction curves) using the mirl function [114] over 1000 iterations, excluding samples below this threshold. Long-read 16S sequencing assigned 36.22% and 77.23% of sequences to species and genus levels respectively, with 99.96% reaching at least family-level classification. To analyze diversity below genus level, we clustered ASVs into operational taxonomic units (OTUs) at 99% similarity, yielding 14,772 OTUs (1890 with species-level and 1412 with genus-level assignments).

We constructed a phylogenetic tree of OTU sequences using the align-to-tree-mafft-iqtree pipeline in QIIME2 [115] with a maximum gap frequency of 0.9 and optimal substitution model via IQ-TREE [116].

### Diet metabarcoding library preparation and processing

We characterised finch diets using DNA metabarcoding of the same fecal samples with two marker genes: rbcL for plant material (primers rbcL1 f: 5*^′^*-TTGGCAGCATTY CGAGTAACTCC-3*^′^*, rbcLB r: 5*^′^*-AACCYTCTTCAAAAAGGTC-3*^′^*; fragment length ~184 bp [117]) and COI for arthropod material (primers Lep f: 5*^′^*-ATTCAACCAATC ATAAAGATATTGG-3*^′^* [118], ZBJ-ArtR2c-deg r: 5*^′^*-WACTAATCAATTWCCAAAH CCHCC-3*^′^*; fragment length *∼*178 bp [119]). Library preparation and sequencing were performed by AllGenetics & Biology SL using a two-step PCR protocol. First-step PCRs were carried out in triplicate in a final volume of 12.5 µL, containing 3.63 µL of template DNA, primers at 0.5 µM final concentration, 6.25 µL of Supreme NZYTaq 2× Green Master Mix (NZYTech), CES (1×), and ultrapure water to volume. Cycling conditions were: initial denaturation at 95 °C for 5 min; 35 cycles of 95 *^◦^*C for 30 s, 49 *^◦^*C for 45 s, 72 *^◦^*C for 45 s; and a final extension at 72 *^◦^*C for 7 min. The three replicate PCR products were pooled for each marker, then the rbcL and COI products were combined and oligonucleotide indices were attached in a second amplification step under identical conditions but with 5 cycles and a 60 *^◦^*C annealing temperature. A no-template negative control was included in every PCR round to check for contamination. Libraries were purified using Mag-Bind RXNPure Plus magnetic beads (Omega Biotek) following the manufacturer’s instructions, quantified using the Qubit dsDNA HS Assay (Thermo Fisher Scientific), pooled in equimolar amounts, and purified again to reduce dimer carryover. The pool was sequenced in a fraction of a NovaSeq PE250 flow cell (Illumina) aiming for a total output of 6 gigabases.

Raw data were demultiplexed by region using cutadapt [120] and separately for each marker using the DADA2 workflow [111]. We trimmed primers, filtered low-quality reads (minLen = 175, maxLen = 184, maxN = 0, maxEE = c(2,2), truncQ = 2), and performed denoising. For taxonomy assignment, we built custom reference databases for rbcL and COI markers using the DB4Q2 pipeline [121], which generates curated baselines imported into QIIME2 to create RDP-trained classifier files.

We clustered ASVs into 99% similarity OTUs and binned data to the lowest taxonomic level (73.8% and 87.4% of OTUs reached genus and family levels, respectively). After merging rbcL and COI abundance tables, we rarefied samples to 1000 sequences per sample using the mirl function [114] over 1000 iterations, excluding samples below this sequencing depth.

### Whole-genome sequencing and population genomics

We extracted DNA from blood samples using the ammonium acetate method (https://dx.doi.org/10.17504/protocols.io.knycvfw) and conducted library preparation to a mean depth of 11.7x (range: 9.4 - 16.9X) using 150-bp paired-end reads on a NovaSeq 6000 platform (Illumina).

We processed raw reads using a custom nextflow pipeline consisting of three main steps (https://github.com/EcoEvoGenomics/genotyping_pipeline/tree/main), version 2.2.4. First, we mapped adapter-trimmed reads to the *Fringilla coelebs* [122] reference genome using bwa-mem and flagged duplicates with picard MarkDuplicates (http://broadinstitute.github.io/picard/). Second, we generated 10 Mb windows across genomes for parallelised genotyping and variant calling using bcftools mpileup, call, norm, and concat tools. Finally, we filtered the resulting VCF files using vcftools (v0.1.16retaining sites with a minimum site-level quality score of 30, requiring a minimum genotype depth of 3X, and excluding sites with more than 50% missing genotypes. We additionally required a minimum minor allele count (MAC) of 3 and excluded the mitochondrial genome and Z chromosome from downstream analyses. We excluded one individual with call rates *<*90%, yielding a final dataset of 51,639,305 SNPs.

We estimated genome-wide differentiation (*F_ST_* ) between populations using a filtered biallelic dataset of 20,694,683 autosomal SNPs (minor allele count *≥* 3) with vcftools *weir-fst-pop* function. We used the –roh (–G30 and default allele frequency settings) algorithm in BCFtools to infer runs of homozygosity (ROH). To calculate *F_ROH_*(proportion of autosomal genome in ROHs), we kept ROHs *>*100-kb *<*1Mb long and with at least 100 SNPs. Principal component analysis was performed using PLINK2 (v 2.00a2.3) [123]. For phylogenetic inference, we generated mitochondrial consensus genomes with BCFtools (v1.19; call -m) under haploid ploidy settings and extracted fasta sequences with samtools (v1.19). Consensus sequences were concatenated, aligned in MAFFT (v7.49, 1000 iterations), and used to reconstruct maximum likelihood trees in IQ-TREE (v2.1.1) [116] with optimal site model selection and 1000 ultrafast bootstraps [124].

### Statistical analysis

#### Community composition and diversity patterns

Because species and island of origin were not fully nested (some species occurred on one island whilst others were on multiple islands and some islands harboured more than one species), we created a combined ’species–population’ grouping variable for all analyses.

We tested predictions of microbiome diversity from host-mediated dispersal using a general linear model (GLM) with negative binomial distribution, microbiome richness as the response variable and sex, age and population as explanatory variables. As age and sex consistently did not explain variation in microbiome diversity, we refitted the models with population as the only explanatory variable. As a sensitivity analysis that accounts for bacterial phylogeny, we repeated the above GLMs using Faith’s PD, which was estimated using the calculatePD function in biomeUtils [125].

We characterised microbiome composition using Bray-Curtis (abundance-weighted) and Jaccard (presence-absence) indices using the adespatial package [126]. To account for phylogeny, we additionally characterised microbiome composition using the weighted and unweighted UniFrac distance metrics [127]. Diet composition was characterised using the Raup-Crick dissimilarity index using the raupcrick function in the vegan package [128]. Species and population-level differences were evaluated using PERMANOVA [128] (9999 permutations, marginal tests) with age and sex as covariates. Temporal effects (sampling year and breeding season) could not be included as covariates because these factors were completely confounded with island of origin: each island was sampled during a single field season, and logistical constraints prevented repeated annual or seasonal sampling within islands. Consequently, variation associated with year/season is inseparable from island effects in our dataset.

To understand ecological processes underlying microbiome composition differences (i.e. dissimilarity), we partitioned beta diversity into turnover (species replacement along spatial or environmental gradients) and nestedness (reflecting species loss where communities are subsets of richer sites) components [129–132] using the beta.div.comp function [126]. We applied this partitioning to both abundance data (Bray-Curtis-derived percentage difference) and presence-absence data (Jaccard index) from the Baselga family of indices.

#### Island biogeography analyses

We tested predictions of island biogeography theory [73] using generalised linear mixed models (GLMMs) with microbiome richness as the response variable, island area (km2), distance from mainland, and island age (mya) as predictors, and population as a random factor. Models were fitted using glmmTMB [133] with negative binomial distribution. We used phyloseq [127] to estimate microbiome richness. As a sensitivity analysis, we repeated the GLMMs using Faith’s PD (estimated as described above).

We assessed whether individuals from the same species complex (i.e. common vs blue chaffinches) shared more microbial taxa than those occupying the same island by focusing on sympatric populations in Gran Canaria and Tenerife. We calculated shared species proportions using the binary Jaccard similarity index and evaluated differences with two-sided permutation *t*-tests (Bonferroni-corrected *p*-values). To account for bacterial phylogenetic relationships, we repeated the above analysis using unweighted UniFrac.

Geographic distance effects on microbiome dissimilarity were evaluated using Mantel tests (9999 permutations) [128] with pairwise geographic distances calculated using the geosphere package [134].

#### Geography, diet and host genetic effects

We assessed three complementary aspects of host genetic variation in relation to microbiome composition. Mitochondrial phylogenetic distance captures evolutionary divergence and shared ancestry among individuals, reflecting lineage-specific effects and phylosymbiosis patterns; nuclear *F_ST_* measures population-level genomic differentiation, testing whether genetically divergent populations harbour divergent microbiomes; and nuclear *F_ROH_* quantifies individual-level inbreeding and heterozygosity loss, testing whether genetic diversity affects microbiome stability within populations. All analyses used both Bray-Curtis (abundance-weighted) and Jaccard (presence-absence) microbiome dissimilarities, computed at every taxonomic level (phylum to OTU).

At the individual level, we assessed the relative contribution of geography, diet, and host phylogenetic distance to microbiome dissimilarity using multiple regression on distance matrices (MRM) [135] and variance partitioning [136] to estimate the variation explained uniquely and jointly by these factors. Host phylogenetic distance was estimated from a mitochondrial tree using cophenetic.phylo [137], as the non-recombining, uniparental inheritance of mitochondrial DNA provides a single genealogy appropriate for individual-level phylogenetic distances. Pairwise distances from the diet dataset were calculated using the raupcrick() function in the vegan package [128].

At the population level, we used partial Mantel tests to assess the relationship between genetic differentiation (*F_ST_* ) and microbiome dissimilarity, controlling for log-transformed geographic distance. For each population, 9 individuals were randomly subsampled and merged into a single community using the merge_samples() and subset_samples() functions [127]. Mantel r and its permutation *p*-value were estimated as the median across 1000 independent random subsamples, with uncertainty summarised as the 2.5th–97.5th percentile interval of partial Mantel *r* across draws.

Finally, to test the relationship between individual inbreeding (*F_ROH_*) and microbiome divergence, we computed each individual’s distance to its population’s median in *β*-diversity space using betadisper [128]. We modelled this divergence using GLMMs [133] with a Gaussian distribution and population as a random effect.

#### Neutral community assembly

We applied the Prokaryote Neutral Model (PNM) as done in Burns et al. [138] to distinguish taxa following neutral expectations from those potentially under host control using the fit_sncm() function in the tyRa package [139]. The PNM uses abundance-occupancy distributions under the null hypothesis of unlimited dispersal and equal fitness [65–67], identifying taxa above (host-selected) or below (dispersal-limited or selected against) neutral expectations. These two groups were collectively treated as a single ”non-neutral” variable to evaluate whether a deviation from neutrality was associated with stronger host-microbe phylogenetic associations.

#### Co-phylogeny and phylogenetic congruence analyses

We tested for host-microbiome co-phylogenetic signals using PACo (Procrustes Approach to Cophylogeny) [68, 69] applying it to paired individual host mitochondrial and microbial OTU-family phylogenies and presence-absence abundance matrices (Cailliez-corrected), with quasiswap null models and 9999 permutations. We compared global and neutrality-partitioned datasets by residual sum of squares goodness-of-fit.

For core microbial families (present on all islands and populations, prevalence *≥* 2 individuals per population), PACo was run separately on OTU and species-level clusters. We repeated analyses using random host trees to test the robustness of PACo to false positive co-phylogeny results. Significant co-phylogeny signals were Bonferroni-corrected.

To discern phylogenetic congruence from co-phylogeny [62], we performed event-based reconciliation analyses with eMPRess [72], subsampling one host population per microbial variant and testing various cost parameters. Phylogenetic congruence was supported if reconciliations showed significantly low costs and co-speciation/transfer event ratios above one [62].

## Data availability

All fastq files have been deposited in ENA sequence read archive under the accession number ERP150413. All the scripts and data for performing the analyses are available through the following link: https://github.com/bkmontero/ChaffinchMicrobiome

## Acknowledgements

We are grateful to Kasun Bodawatta and an anonymous reviewer for their thorough and constructive feedback during the review process, and Sarah Worsely for comments on an early version of this manuscript. We thank Hanna Prüter, Bart Kampenaers, Leo Chaves, and Vivi Porras for many discussions and helpful suggestions. Sara Alberdi assisted with host DNA extraction. Elena Nicoĺas, Eduardo González, Félix Medina, Ángel C. Moreno, Felipe Rodŕıguez, David P. Padilla, Domingo Trujillo, Alejandro Delgado, and Tiago Rodrigues contributed to the fieldwork. Francisco Camacho provided faecal samples from the Iberian Peninsula. Abelardo Margolles provided access to the high-speed benchtop homogenizer (FastPrep) at IPLA-CSIC. We are grateful to CESGA (Centro de Supercomputacíon de Galicia) for providing the computational resources used to perform the genomic analyses. This project was funded by two research grants from the Spanish Ministry of Science, Innovation and Universities and the European Regional Development Fund (PGC2018-097575-B-I00; PID2022-140091NB-I00) and a EECG research award granted to B.K.M. and M.A.F.G by the American Genetic Association.

## Ethics statement

All fieldwork and specimen sampling were conducted in compliance with institutional animal care and use policies. Capture and ringing of birds were authorised by the Regional Government of the Canary Islands (Exp. 2019/11937 and 2020/16669), the Regional Government of the Azores (Licença N° 5/2021/DRAAC), the Regional Government of Madeira (N° 04/IFCN/2020), and the Principality of Asturias (DECO/2021/19003). Additional permissions to conduct research within protected areas were obtained from the Cabildo of Gran Canaria (Exp. FA 16/2020), La Palma (Ref: A/EST-012/2019), Tenerife (Exp. AFF 141/19), and the National Park of Garajonay (Exp. 190.673, PTSS/2.206 and 3.129). Ethical approval for blood sampling was granted by the Regional Government of Asturias (PROAE 45/2019).

## Author contributions

B.K.M., M.G., and J.C.I. conceived and designed the study. J.C.I. coordinated fieldwork and collected samples. B.K.M. and M.R. performed whole-genome data analyses. B.K.M., M.G., and J.C.I. performed DNA extractions. B.K.M. and M.G. conducted metabarcoding data processing and statistical analyses. B.K.M. wrote the first manuscript draft with input from M.G. All authors discussed the results and contributed to the final manuscript.

