## Supplementary Material for "Environmental and genetic drivers differentially shaped resident and transient gut microbiota during the Macaronesian chaffinch radiation"

### Contents

#### Supplementary Figures


- Figure S1 – Faith's PD and UniFrac patterns
- Figure S2 – Ternary plots, abundance-based beta diversity
- Figure S3 – Ternary plots, Jaccard-based beta diversity
- Figure S4 – Shared bacterial species among sympatric finches
- Figure S5 – Prokaryotic neutral model fit


#### Supplementary Tables


- Table S1 – Sample summary by population
- Table S2 – Per-individual sample metadata
- Table S3 – PERMANOVA results for gut microbiome composition
- Table S4 – GLM results for microbiome richness across populations
- Table S5 – GLM results: microbiome richness vs. island characteristics
- Table S6 – Mantel test results (isolation-by-distance)
- Table S7 – MRM results (host phylogeny, diet, geography)
- Table S8 – PACo co-phylogenetic signal
- Table S9 – emPRess event-based reconciliation


### Supplementary Figures

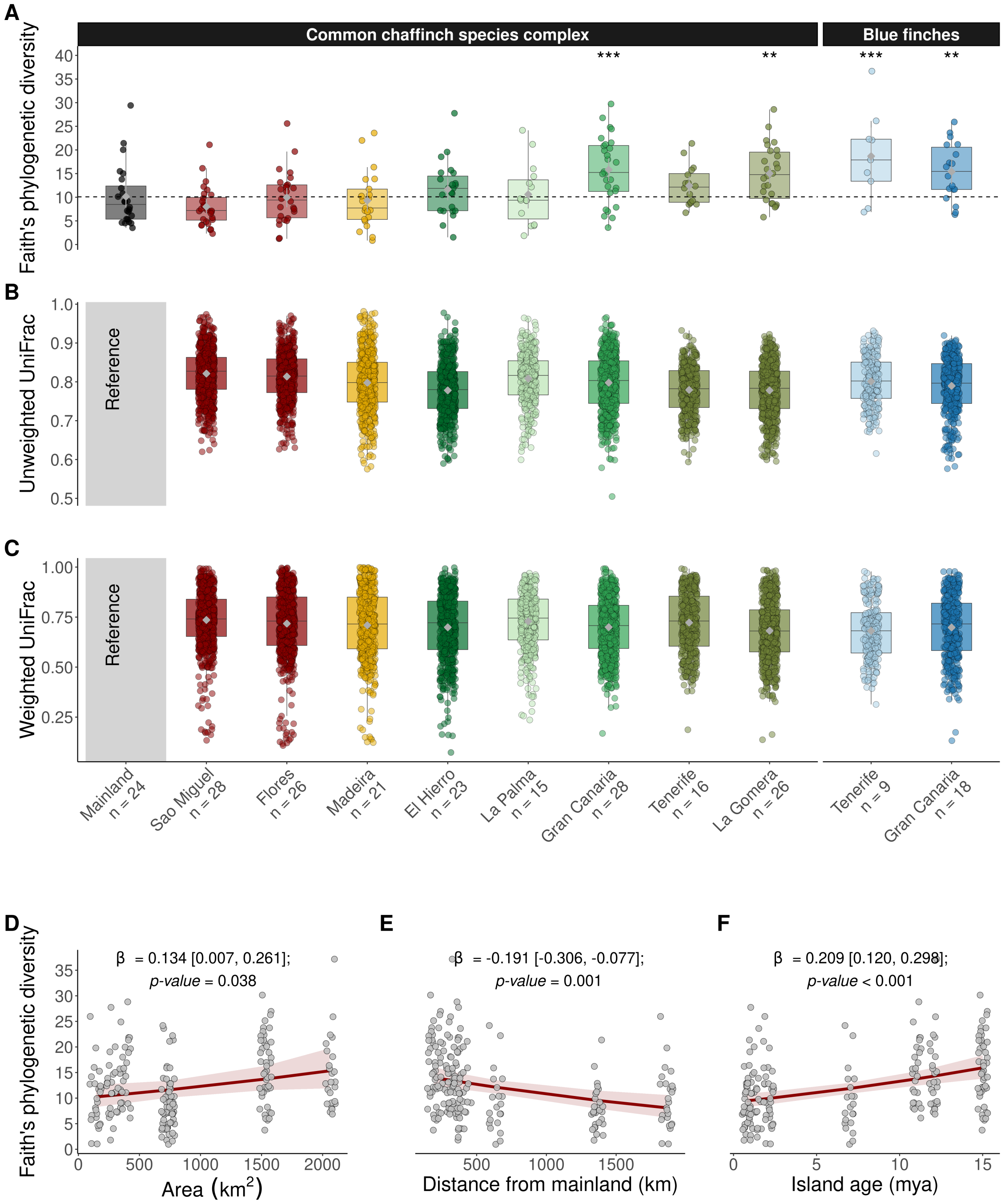


Figure S1: **Microbiome phylogenetic diversity patterns according to colonisation history and biogeography. A**. Species-level Faith's phylogenetic diversity across populations of the common chaffinch complex and blue chaffinches. For the common chaffinch, richness estimates are displayed in the order of the colonisation route: mainland, Azores, Madeira, and Canary Islands. Asterisks indicate significance levels (**P <0.01, ***P <0.001). **B–C**. Microbiome dissimilarity using weighted UniFrac (**B**) and unweighted UniFrac (**C**) between mainland and each Macaronesian population, computed from pairwise distances between every individual on each island and every mainland individual. Box plots show the distribution of pairwise dissimilarities; grey diamonds with whiskers indicate bootstrapped means and 95% confidence intervals. Mainland serves as the reference and is therefore left blank. **D–F**. Relationships between microbiome Faith's phylogenetic diversity and **(D)** island area, **(E)** distance to mainland, and **(F)** island age (negative binomial GLMMs with population as a random effect).

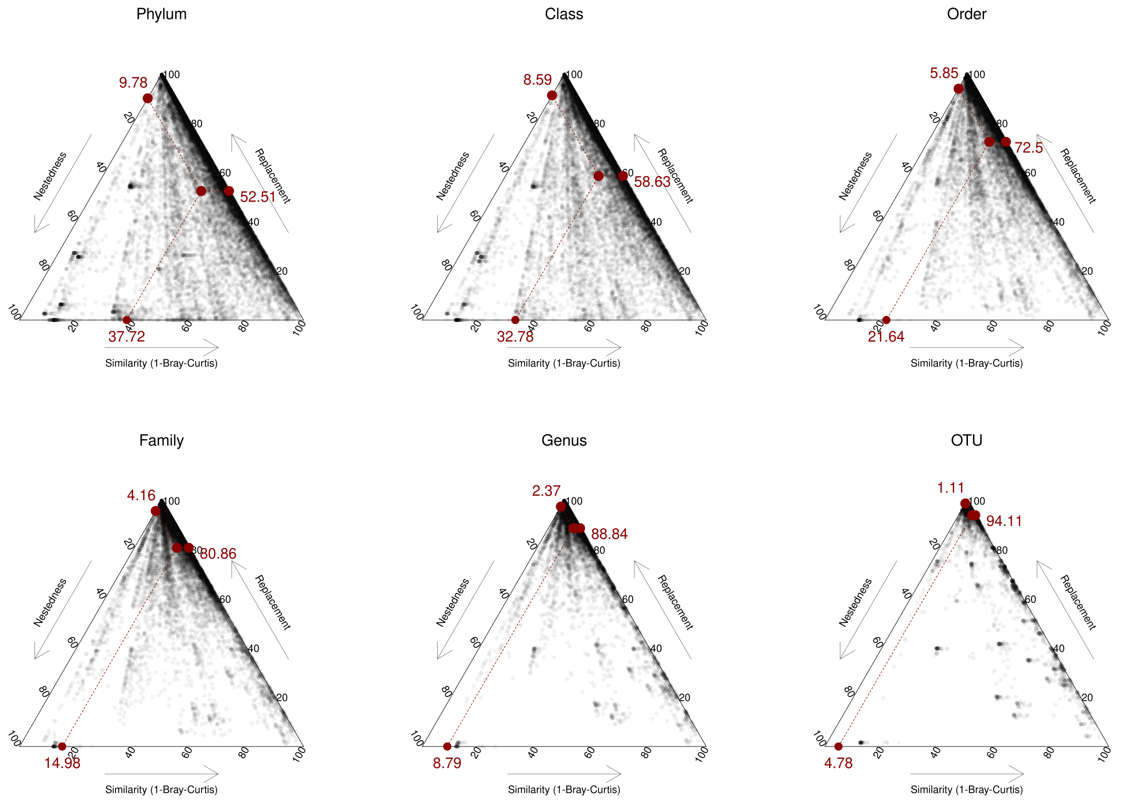


Figure S2: **Ternary plots showing the additive components of beta diversity based on abundance (Bray–Curtis) matrices at different taxonomic levels.** Each black dot corresponds to a pair of common chaffinch individuals from different populations; coordinates reflect the relative contributions of richness difference, replacement (turnover), and similarity. Red dots at the plot edges indicate the mean values of these components, and the central red dot marks the centroid.

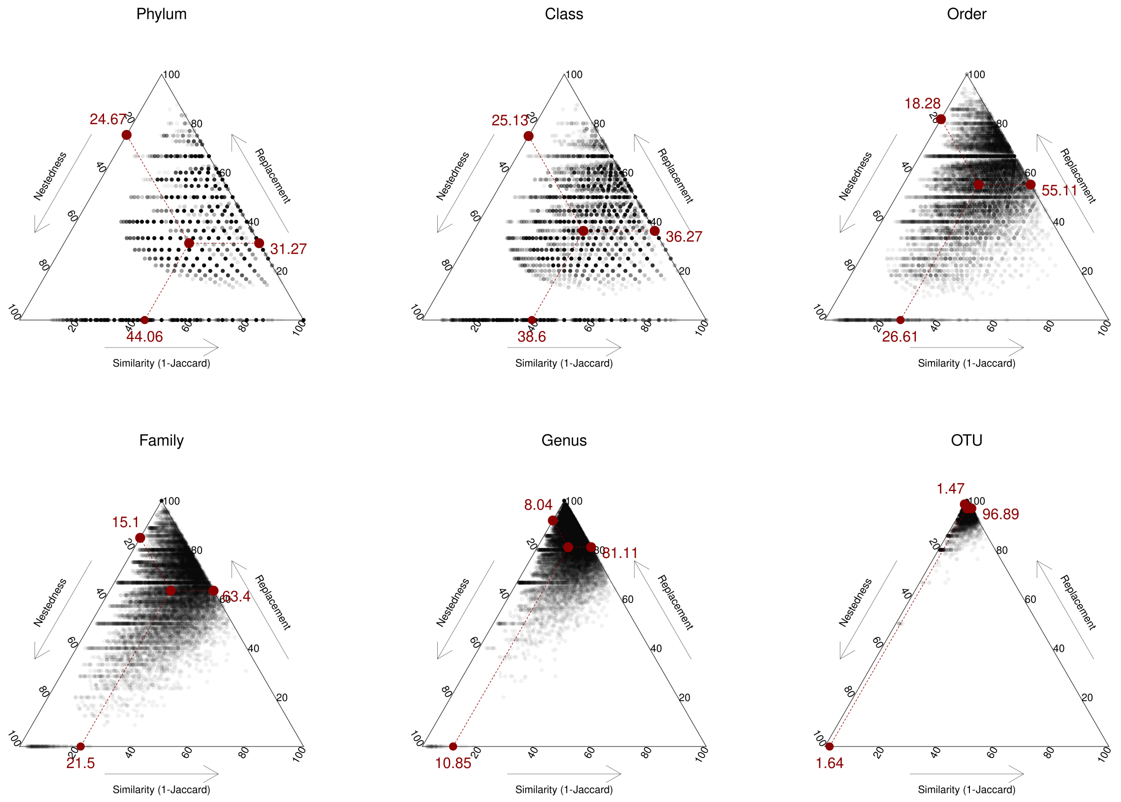


Figure S3: **Ternary plots showing the additive components of beta diversity based on presence–absence (Jaccard) matrices at different taxonomic levels.** Each black dot corresponds to a pair of common chaffinch individuals from different populations; coordinates reflect the relative contributions of richness difference, replacement (turnover), and similarity. Red dots at the plot edges indicate the mean values of these components, and the central red dot marks the centroid.

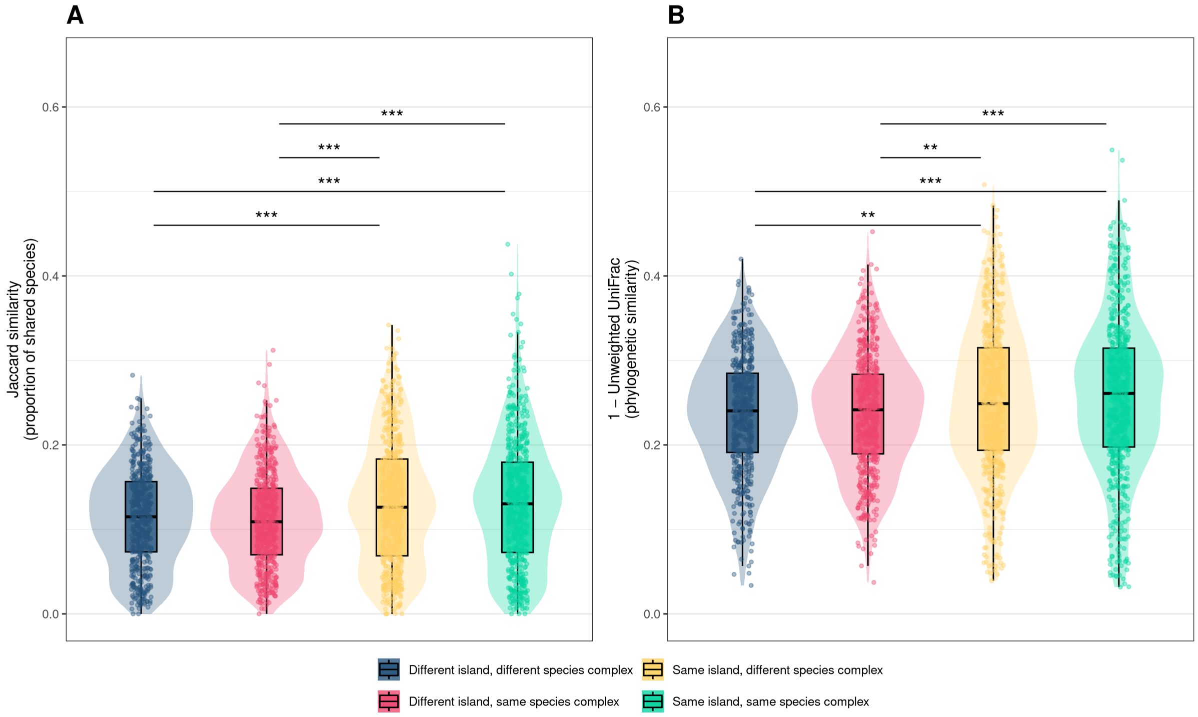


Figure S4: **Shared bacterial species among sympatric finches. A.** Proportion of shared bacterial species (Jaccard similarity) and **B.** phylogenetic similarity (unweighted UniFrac) among sympatric finch populations in Gran Canaria and Tenerife. Asterisks denote levels of statistical significance: **P<0.01; ***P<0.001. .

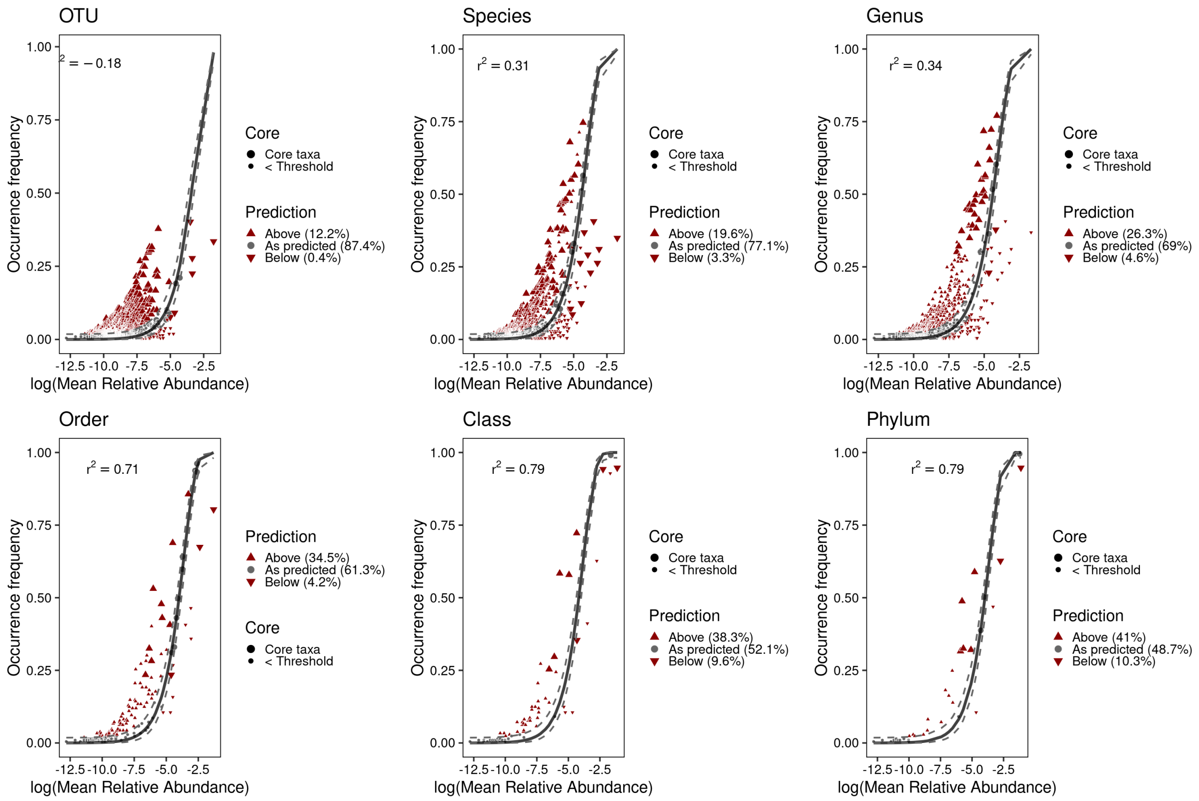


Figure S5: **Model fit to abundance-occupancy distribution expectations of the prokaryotic neutral model (PNM) across taxonomic levels.** Points represent bacterial taxa; the solid line shows the best-fit to neutral expectations, with dashed lines indicating 95% confidence intervals for the model prediction.

### Supplementary Tables

Table S1: **Sample summary by population.** Counts of individuals per population with available microbiome (16S), diet (COI and rbcL), and whole-genome sequencing (WGS) data, together with mean genome-wide $F_{ROH}$ and its 95% confidence interval. 'Diet+microb.' = samples with both microbiome and diet data; 'All three' = samples with microbiome, diet, and WGS data. Dashes indicate populations with no formal subspecies designation.

| Archipelago | Island/Mainland | Genus | Species | Subspecies | Total | $N$ microb. | $N$ diet | $N$ WGS | Diet+microb. | All three | Mean $F_{ROH}$ [95% CI] |
|---|---|---|---|---|---|---|---|---|---|---|---|
| Mainland | Mainland | Fringilla | *coelebs* | – | 28 | 24 | 25 | 2 | 21 | 2 | 0.0297 [0.018–0.0414] |
| Azores | Sao Miguel | Fringilla | *moreletti* | – | 31 | 28 | 28 | 10 | 25 | 10 | 0.2269 [0.2215–0.2323] |
| Azores | Flores | Fringilla | *moreletti* | – | 30 | 26 | 25 | 10 | 21 | 10 | 0.2546 [0.2466–0.2625] |
| Madeira | Madeira | Fringilla | *maderensis* | – | 23 | 21 | 18 | 11 | 16 | 10 | 0.614 [0.6116–0.6164] |
| Canary Islands | El Hierro | Fringilla | *canariensis* | *ombriosa* | 30 | 23 | 25 | 12 | 18 | 11 | 0.716 [0.7128–0.7193] |
| Canary Islands | La Palma | Fringilla | *canariensis* | *palmae* | 23 | 15 | 23 | 11 | 15 | 11 | 0.7756 [0.7714–0.7798] |
| Canary Islands | Gran Canaria | Fringilla | *canariensis* | *bakeri* | 36 | 28 | 34 | 10 | 26 | 10 | 0.7605 [0.7463–0.7747] |
| Canary Islands | Tenerife | Fringilla | *canariensis* | *canariensis* | 22 | 16 | 22 | 9 | 16 | 9 | 0.7872 [0.7797–0.7948] |
| Canary Islands | La Gomera | Fringilla | *canariensis* | *canariensis* | 35 | 26 | 32 | 11 | 23 | 10 | 0.7044 [0.6998–0.709] |
| Canary Islands | Tenerife | Fringilla | *teydea* | – | 12 | 9 | 12 | 9 | 9 | 9 | 0.9243 [0.9224–0.9262] |
| Canary Islands | Gran Canaria | Fringilla | *polatzeki* | – | 32 | 18 | 32 | 11 | 18 | 11 | 0.9206 [0.9183–0.9229] |
| – | – | – | – | **Total** | **302** | **234** | **276** | **106** | **208** | **103** | – |

Table S2: **Sample metadata.** Per-individual NCBI accession number, sampling location, taxonomy, collection date. Columns *Microb.*, *Diet*, and *WGS* indicate whether 16S microbiome, dietary metabarcoding, and whole-genome sequencing data are available (1) or not (0). $F_{ROH}$ is the genome fraction in runs of homozygosity (only computed for individuals with WGS data; dashes denote not applicable).

| Accession | Lab code | Ring | Genus | Species | Subsp. | Date | Lat. | Lon. | Region | Archipelago | Island/Mainland | Sex | Age | Microb. | Diet | WGS | $F_{ROH}$ |
|---|---|---|---|---|---|---|---|---|---|---|---|---|---|---|---|---|---|
| ERS16447426 | 137 | 6L69611 | *Fringilla* | *coelebs* | – | 17/05/2021 | 37.64 | -4.51 | C°rdoba | Mainland | Mainland | female | adult | 1 | 1 | 0 | – |
| ERS16447429 | 140 | E012742 | *Fringilla* | *coelebs* | – | 05/05/2021 | 40.57 | -4.22 | Madrid | Mainland | Mainland | female | adult | 1 | 1 | 0 | – |
| ERS16447431 | 141 | E012799 | *Fringilla* | *coelebs* | – | 24/06/2021 | 40.57 | -4.22 | Madrid | Mainland | Mainland | male | adult | 1 | 1 | 0 | – |
| ERS16447412 | 142 | 4L97202 | *Fringilla* | *coelebs* | – | 29/06/2021 | 40.57 | -4.22 | Madrid | Mainland | Mainland | female | adult | 1 | 1 | 0 | – |
| ERS16447420 | 23 | 6L69608 | *Fringilla* | *coelebs* | – | 13/05/2021 | 37.55 | -4.02 | Jaen | Mainland | Mainland | female | adult | 1 | 1 | 0 | – |
| ERS16447414 | 25 | 6L69612 | *Fringilla* | *coelebs* | – | 09/06/2021 | 38.20 | -3.72 | Jaen | Mainland | Mainland | female | adult | 1 | 1 | 0 | – |
| ERS16447425 | 26 | 6L69604 | *Fringilla* | *coelebs* | – | 13/05/2021 | 37.55 | -4.02 | Jaen | Mainland | Mainland | female | adult | 1 | 1 | 0 | – |
| ERS16447417 | 27 | 6L69606 | *Fringilla* | *coelebs* | – | 13/05/2021 | 37.55 | -4.02 | Jaen | Mainland | Mainland | male | adult | 1 | 1 | 0 | – |
| ERS16447416 | 279 | 4L88845 | *Fringilla* | *coelebs* | – | 20/05/2021 | 43.10 | -5.02 | Leon | Mainland | Mainland | male | adult | 1 | 0 | 0 | – |
| ERS16447423 | 28 | E012775 | *Fringilla* | *coelebs* | – | 04/06/2021 | 40.57 | -4.22 | Madrid | Mainland | Mainland | female | – | 1 | 1 | 0 | – |
| ERS16447430 | 280 | Sin anilla | *Fringilla* | *coelebs* | – | 20/05/2021 | 43.10 | -5.02 | Leon | Mainland | Mainland | male | adult | 1 | 0 | 0 | – |
| ERS16447413 | 281 | 6L45722 | *Fringilla* | *coelebs* | – | 21/04/2021 | 40.57 | -4.22 | Madrid | Mainland | Mainland | male | adult | 1 | 1 | 1 | 0.031 |
| ERS16447433 | 282 | 4L88844 | *Fringilla* | *coelebs* | – | 18/04/2021 | 40.53 | -3.94 | Madrid | Mainland | Mainland | male | adult | 1 | 1 | 1 | 0.029 |
| ERS16447415 | 29 | E012760 | *Fringilla* | *coelebs* | – | 05/06/2021 | 40.57 | -4.22 | Madrid | Mainland | Mainland | female | – | 1 | 1 | 0 | – |
| ERS16447422 | 30 | E012774 | *Fringilla* | *coelebs* | – | 04/06/2021 | 40.57 | -4.22 | Madrid | Mainland | Mainland | female | – | 1 | 1 | 0 | – |
| ERS16447421 | 31 | 5L59562 | *Fringilla* | *coelebs* | – | 06/05/2021 | 40.57 | -4.22 | Madrid | Mainland | Mainland | male | – | 1 | 1 | 0 | – |
| ERS16447419 | 32 | E012798 | *Fringilla* | *coelebs* | – | 24/06/2021 | 40.57 | -4.22 | Madrid | Mainland | Mainland | male | – | 1 | 1 | 0 | – |
| ERS16447434 | 437 | 6L61451 | *Fringilla* | *coelebs* | – | 23/06/2020 | 38.20 | -3.72 | Jaen | Mainland | Mainland | male | adult | 1 | 1 | 0 | – |
| ERS16447428 | 438 | 6L61455 | *Fringilla* | *coelebs* | – | 29/06/2020 | 38.20 | -3.72 | Jaen | Mainland | Mainland | – | fledging | 1 | 1 | 0 | – |
| ERS16447418 | 439 | 6L61456 | *Fringilla* | *coelebs* | – | 13/07/2020 | 38.20 | -3.72 | Jaen | Mainland | Mainland | – | fledging | 1 | 1 | 0 | – |
| ERS16447427 | 441 | 6L61459 | *Fringilla* | *coelebs* | – | 27/07/2020 | 38.20 | -3.72 | Jaen | Mainland | Mainland | male | adult | 1 | 1 | 0 | – |
| ERS16447424 | 443 | 6L61472 | *Fringilla* | *coelebs* | – | 31/08/2020 | 38.20 | -3.72 | Jaen | Mainland | Mainland | female | adult | 1 | 1 | 0 | – |
| ERS16447432 | 444 | 6L61477 | *Fringilla* | *coelebs* | – | 08/09/2020 | 38.20 | -3.72 | Jaen | Mainland | Mainland | female | adult | 1 | 0 | 0 | – |
| ERS16447378 | 81 | 4L95167 | *Fringilla* | *coelebs* | – | 03/05/2018 | 35.90 | -5.30 | Ceuta | Mainland | Mainland | female | adult | 1 | 1 | 0 | – |
| ERS16447438 | 24 | 6L69605 | *Fringilla* | *coelebs* | – | – | 37.55 | -4.02 | – | Mainland | Mainland | male | – | 0 | 1 | 0 | – |
| ERS16447435 | 440 | 6L61457 | *Fringilla* | *coelebs* | – | – | 38.20 | -3.72 | – | Mainland | Mainland | female | – | 0 | 1 | 0 | – |
| ERS16447436 | 442 | 6L61461 | *Fringilla* | *coelebs* | – | – | 38.20 | -3.72 | – | Mainland | Mainland | female | – | 0 | 1 | 0 | – |
| ERS16447437 | 445 | 6L61478 | *Fringilla* | *coelebs* | – | – | 38.20 | -3.72 | – | Mainland | Mainland | – | – | 0 | 1 | 0 | – |
| ERS16447329 | 3 | A443796 | *Fringilla* | *moreletti* | – | 10/06/2021 | 39.38 | -31.19 | Flores | Azores | Flores | male | adult | 1 | 1 | 1 | 0.260 |
| ERS16447330 | 301 | A443795 | *Fringilla* | *moreletti* | – | 10/06/2021 | 39.38 | -31.19 | Flores | Azores | Flores | male | adult | 1 | 1 | 0 | – |
| ERS16447309 | 313 | A443768 | *Fringilla* | *moreletti* | – | 05/06/2021 | 39.39 | -31.17 | Flores | Azores | Flores | female | adult | 1 | 1 | 0 | – |
| ERS16447338 | 350 | A443798 | *Fringilla* | *moreletti* | – | 10/06/2021 | 39.44 | -31.25 | Flores | Azores | Flores | male | adult | 1 | 1 | 1 | 0.244 |
| ERS16447351 | 353 | A443781 | *Fringilla* | *moreletti* | – | 07/06/2021 | 39.39 | -31.21 | Flores | Azores | Flores | male | adult | 1 | 1 | 1 | 0.257 |
| ERS16447315 | 355 | A443766 | *Fringilla* | *moreletti* | – | 05/06/2021 | 39.40 | -31.17 | Flores | Azores | Flores | female | adult | 1 | 1 | 0 | – |
| ERS16447321 | 356 | A443797 | *Fringilla* | *moreletti* | – | 10/06/2021 | 39.44 | -31.25 | Flores | Azores | Flores | male | adult | 1 | 0 | 0 | – |
| ERS16447311 | 357 | A443770 | *Fringilla* | *moreletti* | – | 06/06/2021 | 39.39 | -31.16 | Flores | Azores | Flores | male | adult | 1 | 1 | 0 | – |

*Continued on next page*

Table 2 continued from previous page

| Accession | Lab code | Ring | Genus | Species | Subsp. | Date | Lat. | Lon. | Region | Archipelago | Island/Mainland | Sex | Age | Microb. | Diet | WGS | $F_{ROH}$ |
|---|---|---|---|---|---|---|---|---|---|---|---|---|---|---|---|---|---|
| ERS16447300 | 358 | A443769 | *Fringilla* | *moreletti* | – | 06/06/2021 | 39.39 | -31.16 | Flores | Azores | Flores | male | adult | 1 | 0 | 0 | – |
| ERS16447326 | 371 | A443772 | *Fringilla* | *moreletti* | – | 06/06/2021 | 39.43 | -31.26 | Flores | Azores | Flores | male | adult | 1 | 1 | 0 | – |
| ERS16447323 | 372 | A443774 | *Fringilla* | *moreletti* | – | 06/06/2021 | 39.43 | -31.21 | Flores | Azores | Flores | male | adult | 1 | 1 | 0 | – |
| ERS16447301 | 377 | A443776 | *Fringilla* | *moreletti* | – | 06/06/2021 | 39.43 | -31.21 | Flores | Azores | Flores | male | adult | 1 | 1 | 0 | – |
| ERS16447332 | 378 | A443780 | *Fringilla* | *moreletti* | – | 07/06/2021 | 39.48 | -31.15 | Flores | Azores | Flores | male | adult | 1 | 1 | 1 | 0.264 |
| ERS16447298 | 380 | A443777 | *Fringilla* | *moreletti* | – | 07/06/2021 | 39.39 | -31.17 | Flores | Azores | Flores | male | adult | 1 | 0 | 0 | – |
| ERS16447341 | 393 | A492502 | *Fringilla* | *moreletti* | – | 11/06/2021 | 39.38 | -31.19 | Flores | Azores | Flores | female | adult | 1 | 1 | 1 | 0.244 |
| ERS16447347 | 395 | A492505 | *Fringilla* | *moreletti* | – | 11/06/2021 | 39.38 | -31.19 | Flores | Azores | Flores | male | adult | 1 | 1 | 0 | – |
| ERS16447335 | 396 | A492506 | *Fringilla* | *moreletti* | – | 11/06/2021 | 39.38 | -31.19 | Flores | Azores | Flores | male | adult | 1 | 1 | 0 | – |
| ERS16447342 | 397 | A492507 | *Fringilla* | *moreletti* | – | 11/06/2021 | 39.38 | -31.19 | Flores | Azores | Flores | male | adult | 1 | 1 | 0 | – |
| ERS16447346 | 398 | A492509 | *Fringilla* | *moreletti* | – | 11/06/2021 | 39.38 | -31.19 | Flores | Azores | Flores | male | adult | 1 | 0 | 0 | – |
| ERS16447304 | 419 | A443782 | *Fringilla* | *moreletti* | – | 08/06/2021 | 39.40 | -31.17 | Flores | Azores | Flores | female | adult | 1 | 1 | 0 | – |
| ERS16447306 | 420 | A443783 | *Fringilla* | *moreletti* | – | 08/06/2021 | 39.39 | -31.21 | Flores | Azores | Flores | male | adult | 1 | 0 | 0 | – |
| ERS16447334 | 424 | A443791 | *Fringilla* | *moreletti* | – | 09/06/2021 | 39.44 | -31.16 | Flores | Azores | Flores | male | adult | 1 | 1 | 1 | 0.258 |
| ERS16447345 | 425 | A443799 | *Fringilla* | *moreletti* | – | 11/06/2021 | 39.38 | -31.19 | Flores | Azores | Flores | male | adult | 1 | 1 | 1 | 0.238 |
| ERS16447336 | 426 | A443784 | *Fringilla* | *moreletti* | – | 09/06/2021 | 39.38 | -31.20 | Flores | Azores | Flores | male | adult | 1 | 1 | 1 | 0.244 |
| ERS16447344 | 6 | A492503 | *Fringilla* | *moreletti* | – | 11/06/2021 | 39.38 | -31.19 | Flores | Azores | Flores | male | adult | 1 | 1 | 1 | 0.268 |
| ERS16447333 | 9 | A492504 | *Fringilla* | *moreletti* | – | 11/06/2021 | 39.38 | -31.19 | Flores | Azores | Flores | female | adult | 1 | 1 | 1 | 0.268 |
| ERS16447294 | 107 | A492501 | *Fringilla* | *moreletti* | – | – | 39.38 | -31.19 | – | Azores | Flores | male | – | 0 | 1 | 0 | – |
| ERS16447352 | 295 | A443793 | *Fringilla* | *moreletti* | – | – | 39.38 | -31.19 | – | Azores | Flores | male | – | 0 | 1 | 0 | – |
| ERS16447353 | 302 | A443794 | *Fringilla* | *moreletti* | – | – | 39.38 | -31.19 | – | Azores | Flores | female | – | 0 | 1 | 0 | – |
| ERS16447297 | 370 | A443771 | *Fringilla* | *moreletti* | – | – | 39.43 | -31.26 | – | Azores | Flores | female | – | 0 | 1 | 0 | – |
| ERS16447328 | 100 | A443713 | *Fringilla* | *moreletti* | – | 23/08/2019 | 37.75 | -25.28 | Sao Miguel | Azores | Sao Miguel | male | adult | 1 | 1 | 0 | – |
| ERS16447312 | 101 | A443714 | *Fringilla* | *moreletti* | – | 23/08/2019 | 37.75 | -25.28 | Sao Miguel | Azores | Sao Miguel | male | adult | 1 | 1 | 1 | 0.221 |
| ERS16447307 | 102 | A443716 | *Fringilla* | *moreletti* | – | 23/08/2019 | 37.83 | -25.26 | Sao Miguel | Azores | Sao Miguel | male | adult | 1 | 1 | 0 | – |
| ERS16447317 | 103 | A443728 | *Fringilla* | *moreletti* | – | 26/08/2019 | 37.82 | -25.23 | Sao Miguel | Azores | Sao Miguel | female | fledging | 1 | 0 | 0 | – |
| ERS16447340 | 104 | A443720 | *Fringilla* | *moreletti* | – | 26/08/2019 | 37.77 | -25.34 | Sao Miguel | Azores | Sao Miguel | male | fledging | 1 | 0 | 0 | – |
| ERS16447337 | 105 | A443724 | *Fringilla* | *moreletti* | – | 26/08/2019 | 37.77 | -25.33 | Sao Miguel | Azores | Sao Miguel | male | fledging | 1 | 1 | 1 | 0.227 |
| ERS16447308 | 106 | A443718 | *Fringilla* | *moreletti* | – | 26/08/2019 | 37.77 | -25.34 | Sao Miguel | Azores | Sao Miguel | female | fledging | 1 | 1 | 1 | 0.228 |
| ERS16447331 | 12 | A492513 | *Fringilla* | *moreletti* | – | 13/06/2021 | 37.86 | -25.78 | Sao Miguel | Azores | Sao Miguel | male | adult | 1 | 1 | 0 | – |
| ERS16447343 | 14 | A492515 | *Fringilla* | *moreletti* | – | 13/06/2021 | 37.87 | -25.80 | Sao Miguel | Azores | Sao Miguel | male | adult | 1 | 1 | 1 | 0.226 |
| ERS16447303 | 15 | A492516 | *Fringilla* | *moreletti* | – | 13/06/2021 | 37.87 | -25.80 | Sao Miguel | Azores | Sao Miguel | male | adult | 1 | 0 | 0 | – |
| ERS16447319 | 16 | A492517 | *Fringilla* | *moreletti* | – | 13/06/2021 | 37.84 | -25.78 | Sao Miguel | Azores | Sao Miguel | male | adult | 1 | 1 | 1 | 0.215 |
| ERS16447316 | 17 | A492520 | *Fringilla* | *moreletti* | – | 14/06/2021 | 37.75 | -25.41 | Sao Miguel | Azores | Sao Miguel | male | adult | 1 | 1 | 0 | – |
| ERS16447327 | 18 | A492523 | *Fringilla* | *moreletti* | – | 15/06/2021 | 37.79 | -25.64 | Sao Miguel | Azores | Sao Miguel | female | adult | 1 | 1 | 1 | 0.219 |
| ERS16447325 | 189 | A492546 | *Fringilla* | *moreletti* | – | 19/06/2021 | 37.75 | -25.28 | Sao Miguel | Azores | Sao Miguel | male | adult | 1 | 1 | 1 | 0.239 |
| ERS16447322 | 19 | A492521 | *Fringilla* | *moreletti* | – | 15/06/2021 | 37.79 | -25.64 | Sao Miguel | Azores | Sao Miguel | male | adult | 1 | 1 | 0 | – |
| ERS16447318 | 190 | A492537 | *Fringilla* | *moreletti* | – | 18/06/2021 | 37.78 | -25.39 | Sao Miguel | Azores | Sao Miguel | male | adult | 1 | 1 | 0 | – |
| ERS16447349 | 191 | A492532 | *Fringilla* | *moreletti* | – | 17/06/2021 | 37.86 | -25.78 | Sao Miguel | Azores | Sao Miguel | male | adult | 1 | 1 | 0 | – |
| ERS16447350 | 193 | A492539 | *Fringilla* | *moreletti* | – | 18/06/2021 | 37.82 | -25.23 | Sao Miguel | Azores | Sao Miguel | female | adult | 1 | 1 | 0 | – |

Continued on next page

*Table 2 continued from previous page*

| Accession | Lab code | Ring | Genus | Species | Subsp. | Date | Lat. | Lon. | Region | Archipelago | Island/Mainland | Sex | Age | Microb. | Diet | WGS | $F_{ROH}$ |
|---|---|---|---|---|---|---|---|---|---|---|---|---|---|---|---|---|---|
| ERS16447348 | 195 | A492530 | *Fringilla* | *moreletti* | – | 16/06/2021 | 37.86 | -25.77 | Sao Miguel | Azores | Sao Miguel | female | adult | 1 | 1 | 0 | – |
| ERS16447339 | 196 | A492541 | *Fringilla* | *moreletti* | – | 18/06/2021 | 37.77 | -25.18 | Sao Miguel | Azores | Sao Miguel | male | adult | 1 | 1 | 1 | 0.235 |
| ERS16447320 | 197 | A492529 | *Fringilla* | *moreletti* | – | 16/06/2021 | 37.86 | -25.77 | Sao Miguel | Azores | Sao Miguel | male | adult | 1 | 1 | 0 | – |
| ERS16447302 | 201 | A492527 | *Fringilla* | *moreletti* | – | 16/06/2021 | 37.86 | -25.77 | Sao Miguel | Azores | Sao Miguel | male | adult | 1 | 1 | 0 | – |
| ERS16447324 | 202 | A492544 | *Fringilla* | *moreletti* | – | 19/06/2021 | 37.83 | -25.26 | Sao Miguel | Azores | Sao Miguel | male | adult | 1 | 1 | 1 | 0.234 |
| ERS16447314 | 204 | A492535 | *Fringilla* | *moreletti* | – | 17/06/2021 | 37.87 | -25.77 | Sao Miguel | Azores | Sao Miguel | female | adult | 1 | 1 | 0 | – |
| ERS16447305 | 205 | A492545 | *Fringilla* | *moreletti* | – | 19/06/2021 | 37.80 | -25.16 | Sao Miguel | Azores | Sao Miguel | male | adult | 1 | 1 | 0 | – |
| ERS16447313 | 21 | A492522 | *Fringilla* | *moreletti* | – | 15/06/2021 | 37.79 | -25.64 | Sao Miguel | Azores | Sao Miguel | male | adult | 1 | 1 | 1 | 0.225 |
| ERS16447310 | 231 | A443729 | *Fringilla* | *moreletti* | – | 26/08/2019 | 37.82 | -25.23 | Sao Miguel | Azores | Sao Miguel | male | adult | 1 | 1 | 0 | – |
| ERS16447299 | 99 | A443712 | *Fringilla* | *moreletti* | – | 23/08/2019 | 37.83 | -25.16 | Sao Miguel | Azores | Sao Miguel | male | adult | 1 | 1 | 0 | – |
| ERS16447354 | 13 | A492514 | *Fringilla* | *moreletti* | – | – | 37.87 | -25.80 | – | Azores | Sao Miguel | female | – | 0 | 1 | 0 | – |
| ERS16447295 | 20 | A492524 | *Fringilla* | *moreletti* | – | – | 37.86 | -25.79 | – | Azores | Sao Miguel | male | – | 0 | 1 | 0 | – |
| ERS16447296 | 200 | A492543 | *Fringilla* | *moreletti* | – | – | 37.83 | -25.26 | – | Azores | Sao Miguel | male | – | 0 | 1 | 0 | – |
| ERS16447146 | 287 | A443763 | *Fringilla* | *maderensis* | – | 06/05/2021 | 32.71 | -16.88 | Madeira | Madeira | Madeira | male | adult | 1 | 0 | 0 | – |
| ERS16447162 | 293 | A443752 | *Fringilla* | *maderensis* | – | 03/05/2021 | 32.82 | -17.19 | Madeira | Madeira | Madeira | male | adult | 1 | 1 | 1 | 0.609 |
| ERS16447151 | 312 | A443740 | *Fringilla* | *maderensis* | – | 30/04/2021 | 32.72 | -16.89 | Madeira | Madeira | Madeira | male | adult | 1 | 1 | 0 | – |
| ERS16447155 | 314 | A443759 | *Fringilla* | *maderensis* | – | 05/05/2021 | 32.82 | -17.19 | Madeira | Madeira | Madeira | male | adult | 1 | 1 | 0 | – |
| ERS16447156 | 344 | A443734 | *Fringilla* | *maderensis* | – | 27/04/2021 | 32.72 | -16.86 | Madeira | Madeira | Madeira | male | adult | 1 | 1 | 0 | – |
| ERS16447145 | 345 | A443735 | *Fringilla* | *maderensis* | – | 27/04/2021 | 32.72 | -16.86 | Madeira | Madeira | Madeira | male | adult | 1 | 0 | 0 | – |
| ERS16447149 | 346 | A443738 | *Fringilla* | *maderensis* | – | 29/04/2021 | 32.75 | -16.87 | Madeira | Madeira | Madeira | male | adult | 1 | 1 | 0 | – |
| ERS16447166 | 348 | A443760 | *Fringilla* | *maderensis* | – | 05/05/2021 | 32.82 | -17.19 | Madeira | Madeira | Madeira | male | adult | 1 | 1 | 1 | 0.618 |
| ERS16447147 | 376 | A443744 | *Fringilla* | *maderensis* | – | 30/04/2021 | 32.72 | -16.89 | Madeira | Madeira | Madeira | male | adult | 1 | 0 | 0 | – |
| ERS16447154 | 385 | A443747 | *Fringilla* | *maderensis* | – | 01/05/2021 | 32.82 | -17.15 | Madeira | Madeira | Madeira | male | adult | 1 | 1 | 1 | 0.616 |
| ERS16447163 | 399 | A443751 | *Fringilla* | *maderensis* | – | 03/05/2021 | 32.85 | -17.19 | Madeira | Madeira | Madeira | male | adult | 1 | 1 | 1 | 0.612 |
| ERS16447164 | 401 | A443756 | *Fringilla* | *maderensis* | – | 05/05/2021 | 32.75 | -17.02 | Madeira | Madeira | Madeira | male | adult | 1 | 1 | 1 | 0.618 |
| ERS16447161 | 402 | A443757 | *Fringilla* | *maderensis* | – | 05/05/2021 | 32.74 | -17.06 | Madeira | Madeira | Madeira | male | adult | 1 | 1 | 0 | – |
| ERS16447159 | 403 | A443753 | *Fringilla* | *maderensis* | – | 03/05/2021 | 32.77 | -17.12 | Madeira | Madeira | Madeira | male | adult | 1 | 1 | 1 | 0.611 |
| ERS16447160 | 405 | A443745 | *Fringilla* | *maderensis* | – | 01/05/2021 | 32.84 | -17.15 | Madeira | Madeira | Madeira | male | adult | 1 | 1 | 1 | 0.612 |
| ERS16447165 | 406 | A443742 | *Fringilla* | *maderensis* | – | 30/04/2021 | 32.72 | -16.89 | Madeira | Madeira | Madeira | female | adult | 1 | 0 | 1 | 0.616 |
| ERS16447152 | 407 | A443746 | *Fringilla* | *maderensis* | – | 01/05/2021 | 32.82 | -17.15 | Madeira | Madeira | Madeira | male | adult | 1 | 1 | 0 | – |
| ERS16447148 | 408 | A443743 | *Fringilla* | *maderensis* | – | 30/04/2021 | 32.71 | -16.88 | Madeira | Madeira | Madeira | male | adult | 1 | 0 | 0 | – |
| ERS16447157 | 88 | A443731 | *Fringilla* | *maderensis* | – | 26/04/2021 | 32.71 | -16.88 | Madeira | Madeira | Madeira | male | adult | 1 | 1 | 1 | 0.609 |
| ERS16447158 | 90 | A443733 | *Fringilla* | *maderensis* | – | 27/04/2021 | 32.71 | -16.88 | Madeira | Madeira | Madeira | male | adult | 1 | 1 | 1 | 0.619 |
| ERS16447153 | 91 | A443761 | *Fringilla* | *maderensis* | – | 05/05/2021 | 32.85 | -17.19 | Madeira | Madeira | Madeira | male | adult | 1 | 1 | 1 | 0.613 |
| ERS16447168 | 288 | A443732 | *Fringilla* | *maderensis* | – | – | 32.71 | -16.88 | – | Madeira | Madeira | male | – | 0 | 1 | 0 | – |
| ERS16447167 | 92 | A407539 | *Fringilla* | *maderensis* | – | – | 32.85 | -17.19 | – | Madeira | Madeira | male | – | 0 | 1 | 0 | – |
| ERS16447183 | 255 | 4L23024 | *Fringilla* | *canariensis* | *ombriosa* | 09/03/2020 | 27.74 | -18.00 | El Hierro | Canary Islands | El Hierro | male | adult | 1 | 1 | 1 | 0.718 |
| ERS16447178 | 256 | 4L23022 | *Fringilla* | *canariensis* | *ombriosa* | 09/03/2020 | 27.74 | -18.00 | El Hierro | Canary Islands | El Hierro | female | adult | 1 | 1 | 0 | – |
| ERS16447181 | 257 | 4L23023 | *Fringilla* | *canariensis* | *ombriosa* | 09/03/2020 | 27.74 | -18.00 | El Hierro | Canary Islands | El Hierro | male | adult | 1 | 1 | 1 | 0.713 |
| ERS16447182 | 258 | 4L23025 | *Fringilla* | *canariensis* | *ombriosa* | 09/03/2020 | 27.74 | -18.00 | El Hierro | Canary Islands | El Hierro | male | adult | 1 | 1 | 1 | 0.712 |

*Continued on next page*

*Table 2 continued from previous page*

| Accession | Lab code | Ring | Genus | Species | Subsp. | Date | Lat. | Lon. | Region | Archipelago | Island/Mainland | Sex | Age | Microb. | Diet | WGS | $F_{ROH}$ |
|---|---|---|---|---|---|---|---|---|---|---|---|---|---|---|---|---|---|
| ERS16447177 | 259 | 4L23026 | *Fringilla* | *canariensis* | *ombriosa* | 10/03/2020 | 27.74 | -18.00 | El Hierro | Canary Islands | El Hierro | female | adult | 1 | 1 | 0 | – |
| ERS16447179 | 266 | 4L23027 | *Fringilla* | *canariensis* | *ombriosa* | 10/03/2020 | 27.74 | -18.00 | El Hierro | Canary Islands | El Hierro | male | adult | 1 | 1 | 0 | – |
| ERS16447180 | 317 | 4L23033 | *Fringilla* | *canariensis* | *ombriosa* | 10/03/2020 | 27.74 | -18.00 | El Hierro | Canary Islands | El Hierro | male | adult | 1 | 0 | 0 | – |
| ERS16447195 | 319 | 4L23035 | *Fringilla* | *canariensis* | *ombriosa* | 11/03/2020 | 27.74 | -17.99 | El Hierro | Canary Islands | El Hierro | female | adult | 1 | 0 | 0 | – |
| ERS16447173 | 320 | 4L23034 | *Fringilla* | *canariensis* | *ombriosa* | 11/03/2020 | 27.74 | -17.99 | El Hierro | Canary Islands | El Hierro | male | adult | 1 | 1 | 1 | 0.717 |
| ERS16447188 | 321 | 4L23036 | *Fringilla* | *canariensis* | *ombriosa* | 11/03/2020 | 27.73 | -18.01 | El Hierro | Canary Islands | El Hierro | female | adult | 1 | 1 | 1 | 0.717 |
| ERS16447187 | 324 | 4L23037 | *Fringilla* | *canariensis* | *ombriosa* | 11/03/2020 | 27.73 | -18.01 | El Hierro | Canary Islands | El Hierro | male | adult | 1 | 1 | 0 | – |
| ERS16447176 | 325 | 4L23043 | *Fringilla* | *canariensis* | *ombriosa* | 12/03/2020 | 27.74 | -17.99 | El Hierro | Canary Islands | El Hierro | male | adult | 1 | 1 | 0 | – |
| ERS16447175 | 326 | 4L23038 | *Fringilla* | *canariensis* | *ombriosa* | 12/03/2020 | 27.74 | -17.99 | El Hierro | Canary Islands | El Hierro | female | adult | 1 | 1 | 0 | – |
| ERS16447189 | 327 | 4L23044 | *Fringilla* | *canariensis* | *ombriosa* | 12/03/2020 | 27.74 | -17.99 | El Hierro | Canary Islands | El Hierro | male | adult | 1 | 0 | 0 | – |
| ERS16447190 | 330 | 4L23039 | *Fringilla* | *canariensis* | *ombriosa* | 12/03/2020 | 27.74 | -17.99 | El Hierro | Canary Islands | El Hierro | male | adult | 1 | 1 | 1 | 0.714 |
| ERS16447191 | 332 | 4L23041 | *Fringilla* | *canariensis* | *ombriosa* | 12/03/2020 | 27.74 | -17.99 | El Hierro | Canary Islands | El Hierro | male | adult | 1 | 1 | 0 | – |
| ERS16447174 | 333 | 4L23045 | *Fringilla* | *canariensis* | *ombriosa* | 12/03/2020 | 27.73 | -18.01 | El Hierro | Canary Islands | El Hierro | male | adult | 1 | 0 | 1 | 0.717 |
| ERS16447186 | 334 | 4L23042 | *Fringilla* | *canariensis* | *ombriosa* | 12/03/2020 | 27.74 | -17.99 | El Hierro | Canary Islands | El Hierro | female | adult | 1 | 0 | 0 | – |
| ERS16447194 | 34 | 4L23014 | *Fringilla* | *canariensis* | *ombriosa* | 07/03/2020 | 27.74 | -18.00 | El Hierro | Canary Islands | El Hierro | male | adult | 1 | 1 | 1 | 0.726 |
| ERS16447184 | 38 | 4L23016 | *Fringilla* | *canariensis* | *ombriosa* | 08/03/2020 | 27.74 | -18.00 | El Hierro | Canary Islands | El Hierro | male | adult | 1 | 1 | 1 | 0.708 |
| ERS16447185 | 41 | 4L23021 | *Fringilla* | *canariensis* | *ombriosa* | 09/03/2020 | 27.74 | -18.00 | El Hierro | Canary Islands | El Hierro | male | adult | 1 | 1 | 1 | 0.715 |
| ERS16447193 | 42 | 4L23019 | *Fringilla* | *canariensis* | *ombriosa* | 09/03/2020 | 27.74 | -18.00 | El Hierro | Canary Islands | El Hierro | male | adult | 1 | 1 | 1 | 0.711 |
| ERS16447192 | 43 | 4L23030 | *Fringilla* | *canariensis* | *ombriosa* | 10/03/2020 | 27.74 | -18.00 | El Hierro | Canary Islands | El Hierro | male | adult | 1 | 1 | 1 | 0.725 |
| ERS16447171 | 261 | 4L23028 | *Fringilla* | *canariensis* | *ombriosa* | – | 27.74 | -18.00 | – | Canary Islands | El Hierro | female | – | 0 | 1 | 0 | – |
| ERS16447198 | 303 | 4L23029 | *Fringilla* | *canariensis* | *ombriosa* | – | 27.74 | -18.00 | – | Canary Islands | El Hierro | female | – | 0 | 1 | 0 | – |
| ERS16447199 | 304 | 4L23031 | *Fringilla* | *canariensis* | *ombriosa* | – | 27.74 | -18.00 | – | Canary Islands | El Hierro | male | – | 0 | 1 | 0 | – |
| ERS16447200 | 305 | 4L23032 | *Fringilla* | *canariensis* | *ombriosa* | – | 27.74 | -18.00 | – | Canary Islands | El Hierro | male | – | 0 | 1 | 0 | – |
| ERS16447172 | 331 | 4L23040 | *Fringilla* | *canariensis* | *ombriosa* | – | 27.74 | -17.99 | – | Canary Islands | El Hierro | male | – | 0 | 1 | 0 | – |
| ERS16447196 | 37 | 4L23015 | *Fringilla* | *canariensis* | *ombriosa* | – | 27.74 | -18.00 | – | Canary Islands | El Hierro | male | – | 0 | 1 | 0 | – |
| ERS16447197 | 40 | 4L23017 | *Fringilla* | *canariensis* | *ombriosa* | – | 27.74 | -18.00 | – | Canary Islands | El Hierro | female | – | 0 | 1 | 0 | – |
| ERS16447270 | 206 | 4L95846 | *Fringilla* | *canariensis* | *bakeri* | 01/09/2020 | 27.96 | -15.59 | Gran Canaria | Canary Islands | Gran Canaria | male | fledging | 1 | 1 | 1 | 0.761 |
| ERS16447261 | 207 | 4L95850 | *Fringilla* | *canariensis* | *bakeri* | 01/09/2020 | 27.96 | -15.59 | Gran Canaria | Canary Islands | Gran Canaria | female | fledging | 1 | 1 | 0 | – |
| ERS16447269 | 209 | 4L95847 | *Fringilla* | *canariensis* | *bakeri* | 01/09/2020 | 27.96 | -15.59 | Gran Canaria | Canary Islands | Gran Canaria | male | fledging | 1 | 1 | 0 | – |
| ERS16447278 | 210 | 4L95849 | *Fringilla* | *canariensis* | *bakeri* | 01/09/2020 | 27.96 | -15.59 | Gran Canaria | Canary Islands | Gran Canaria | male | adult | 1 | 1 | 0 | – |
| ERS16447259 | 211 | 4L95821 | *Fringilla* | *canariensis* | *bakeri* | 28/08/2020 | 27.96 | -15.59 | Gran Canaria | Canary Islands | Gran Canaria | male | fledging | 1 | 1 | 0 | – |
| ERS16447280 | 213 | 4L95824 | *Fringilla* | *canariensis* | *bakeri* | 29/08/2020 | 28.09 | -15.59 | Gran Canaria | Canary Islands | Gran Canaria | male | fledging | 1 | 1 | 0 | – |
| ERS16447284 | 214 | 4L95827 | *Fringilla* | *canariensis* | *bakeri* | 29/08/2020 | 28.09 | -15.59 | Gran Canaria | Canary Islands | Gran Canaria | female | adult | 1 | 1 | 1 | 0.769 |
| ERS16447264 | 215 | 4L95830 | *Fringilla* | *canariensis* | *bakeri* | 29/08/2020 | 28.09 | -15.59 | Gran Canaria | Canary Islands | Gran Canaria | female | adult | 1 | 1 | 1 | 0.768 |
| ERS16447287 | 216 | 4L95834 | *Fringilla* | *canariensis* | *bakeri* | 30/08/2020 | 28.08 | -15.58 | Gran Canaria | Canary Islands | Gran Canaria | female | fledging | 1 | 1 | 0 | – |
| ERS16447263 | 217 | 4L95838 | *Fringilla* | *canariensis* | *bakeri* | 30/08/2020 | 28.08 | -15.58 | Gran Canaria | Canary Islands | Gran Canaria | male | fledging | 1 | 1 | 1 | 0.793 |
| ERS16447285 | 222 | 4L95832 | *Fringilla* | *canariensis* | *bakeri* | 29/08/2020 | 28.08 | -15.58 | Gran Canaria | Canary Islands | Gran Canaria | female | fledging | 1 | 1 | 0 | – |
| ERS16447267 | 224 | 4L95840 | *Fringilla* | *canariensis* | *bakeri* | 30/08/2020 | 28.08 | -15.58 | Gran Canaria | Canary Islands | Gran Canaria | male | fledging | 1 | 1 | 0 | – |
| ERS16447268 | 225 | 4L95823 | *Fringilla* | *canariensis* | *bakeri* | 28/08/2020 | 27.96 | -15.59 | Gran Canaria | Canary Islands | Gran Canaria | female | fledging | 1 | 1 | 0 | – |
| ERS16447286 | 226 | 4L95825 | *Fringilla* | *canariensis* | *bakeri* | 29/08/2020 | 28.09 | -15.59 | Gran Canaria | Canary Islands | Gran Canaria | male | adult | 1 | 1 | 1 | 0.773 |

*Continued on next page*

Table 2 continued from previous page

| Accession | Lab code | Ring | Genus | Species | Subsp. | Date | Lat. | Lon. | Region | Archipelago | Island/Mainland | Sex | Age | Microb. | Diet | WGS | $F_{ROH}$ |
|---|---|---|---|---|---|---|---|---|---|---|---|---|---|---|---|---|---|
| ERS16447265 | 227 | 4L95828 | *Fringilla* | *canariensis* | *bakeri* | 29/08/2020 | 28.09 | -15.59 | Gran Canaria | Canary Islands | Gran Canaria | male | fledging | 1 | 1 | 0 | – |
| ERS16447275 | 244 | 4L95837 | *Fringilla* | *canariensis* | *bakeri* | 30/08/2020 | 28.08 | -15.58 | Gran Canaria | Canary Islands | Gran Canaria | male | fledging | 1 | 1 | 0 | – |
| ERS16447262 | 246 | 4L95826 | *Fringilla* | *canariensis* | *bakeri* | 29/08/2020 | 28.09 | -15.59 | Gran Canaria | Canary Islands | Gran Canaria | male | fledging | 1 | 1 | 1 | 0.775 |
| ERS16447276 | 247 | 4L95822 | *Fringilla* | *canariensis* | *bakeri* | 28/08/2020 | 27.96 | -15.59 | Gran Canaria | Canary Islands | Gran Canaria | female | fledging | 1 | 1 | 1 | 0.735 |
| ERS16447266 | 248 | 4L95836 | *Fringilla* | *canariensis* | *bakeri* | 30/08/2020 | 28.08 | -15.58 | Gran Canaria | Canary Islands | Gran Canaria | male | fledging | 1 | 1 | 0 | – |
| ERS16447283 | 252 | 4L95841 | *Fringilla* | *canariensis* | *bakeri* | 31/08/2020 | 27.96 | -15.59 | Gran Canaria | Canary Islands | Gran Canaria | male | fledging | 1 | 1 | 0 | – |
| ERS16447273 | 263 | 4L95842 | *Fringilla* | *canariensis* | *bakeri* | 31/08/2020 | 27.96 | -15.59 | Gran Canaria | Canary Islands | Gran Canaria | male | adult | 1 | 1 | 0 | – |
| ERS16447277 | 268 | 4L95844 | *Fringilla* | *canariensis* | *bakeri* | 31/08/2020 | 27.96 | -15.59 | Gran Canaria | Canary Islands | Gran Canaria | male | fledging | 1 | 1 | 1 | 0.730 |
| ERS16447272 | 273 | 4L95831 | *Fringilla* | *canariensis* | *bakeri* | 29/08/2020 | 28.08 | -15.58 | Gran Canaria | Canary Islands | Gran Canaria | female | fledging | 1 | 1 | 1 | 0.761 |
| ERS16447281 | 69 | 4L95848 | *Fringilla* | *canariensis* | *bakeri* | 01/09/2020 | 27.96 | -15.59 | Gran Canaria | Canary Islands | Gran Canaria | female | adult | 1 | 1 | 0 | – |
| ERS16447282 | 70 | L541866 | *Fringilla* | *canariensis* | *bakeri* | 01/09/2020 | 27.96 | -15.59 | Gran Canaria | Canary Islands | Gran Canaria | female | adult | 1 | 0 | 0 | – |
| ERS16447274 | 73 | L541867 | *Fringilla* | *canariensis* | *bakeri* | 01/09/2020 | 27.96 | -15.59 | Gran Canaria | Canary Islands | Gran Canaria | male | adult | 1 | 1 | 0 | – |
| ERS16447279 | 75 | L541861 | *Fringilla* | *canariensis* | *bakeri* | 01/09/2020 | 27.96 | -15.59 | Gran Canaria | Canary Islands | Gran Canaria | female | fledging | 1 | 1 | 1 | 0.740 |
| ERS16447271 | 78 | L541863 | *Fringilla* | *canariensis* | *bakeri* | 01/09/2020 | 27.96 | -15.59 | Gran Canaria | Canary Islands | Gran Canaria | male | adult | 1 | 0 | 0 | – |
| ERS16447288 | 221 | 4L95829 | *Fringilla* | *canariensis* | *bakeri* | – | 28.09 | -15.59 | – | Canary Islands | Gran Canaria | female | – | 0 | 1 | 0 | – |
| ERS16447290 | 228 | 4L95835 | *Fringilla* | *canariensis* | *bakeri* | – | 28.08 | -15.58 | – | Canary Islands | Gran Canaria | male | – | 0 | 1 | 0 | – |
| ERS16447260 | 241 | 4L95843 | *Fringilla* | *canariensis* | *bakeri* | – | 27.96 | -15.59 | – | Canary Islands | Gran Canaria | female | – | 0 | 1 | 0 | – |
| ERS16447292 | 242 | 4L95845 | *Fringilla* | *canariensis* | *bakeri* | – | 27.96 | -15.59 | – | Canary Islands | Gran Canaria | female | – | 0 | 1 | 0 | – |
| ERS16447289 | 243 | 4L95833 | *Fringilla* | *canariensis* | *bakeri* | – | 28.08 | -15.58 | – | Canary Islands | Gran Canaria | male | – | 0 | 1 | 0 | – |
| ERS16447291 | 272 | 4L95839 | *Fringilla* | *canariensis* | *bakeri* | – | 28.08 | -15.58 | – | Canary Islands | Gran Canaria | male | – | 0 | 1 | 0 | – |
| ERS16447293 | 76 | L541862 | *Fringilla* | *canariensis* | *bakeri* | – | 27.96 | -15.59 | – | Canary Islands | Gran Canaria | male | – | 0 | 1 | 0 | – |
| ERS16447258 | 79 | L541864 | *Fringilla* | *canariensis* | *bakeri* | – | 27.96 | -15.59 | – | Canary Islands | Gran Canaria | male | – | 0 | 1 | 0 | – |
| ERS16447383 | 208 | 2A506481 | *Fringilla* | *polatzeki* | – | 01/09/2020 | 27.96 | -15.59 | Gran Canaria | Canary Islands | Gran Canaria | male | fledging | 1 | 1 | 0 | – |
| ERS16447381 | 212 | 2A493932 | *Fringilla* | *polatzeki* | – | 28/08/2020 | 27.96 | -15.59 | Gran Canaria | Canary Islands | Gran Canaria | male | fledging | 1 | 1 | 0 | – |
| ERS16447392 | 220 | 2A506484 | *Fringilla* | *polatzeki* | – | 01/09/2020 | 27.96 | -15.59 | Gran Canaria | Canary Islands | Gran Canaria | female | adult | 1 | 1 | 1 | 0.920 |
| ERS16447384 | 223 | 2A493931 | *Fringilla* | *polatzeki* | – | 28/08/2020 | 27.96 | -15.59 | Gran Canaria | Canary Islands | Gran Canaria | female | fledging | 1 | 1 | 0 | – |
| ERS16447397 | 240 | 2A493958 | *Fringilla* | *polatzeki* | – | 31/08/2020 | 27.96 | -15.59 | Gran Canaria | Canary Islands | Gran Canaria | female | fledging | 1 | 1 | 1 | 0.920 |
| ERS16447386 | 253 | 2A493955 | *Fringilla* | *polatzeki* | – | 31/08/2020 | 27.96 | -15.59 | Gran Canaria | Canary Islands | Gran Canaria | male | fledging | 1 | 1 | 1 | 0.924 |
| ERS16447395 | 262 | 2A493954 | *Fringilla* | *polatzeki* | – | 31/08/2020 | 27.96 | -15.59 | Gran Canaria | Canary Islands | Gran Canaria | male | fledging | 1 | 1 | 1 | 0.923 |
| ERS16447388 | 264 | 2A493956 | *Fringilla* | *polatzeki* | – | 31/08/2020 | 27.96 | -15.59 | Gran Canaria | Canary Islands | Gran Canaria | male | fledging | 1 | 1 | 1 | 0.921 |
| ERS16447389 | 267 | 2A493957 | *Fringilla* | *polatzeki* | – | 31/08/2020 | 27.96 | -15.59 | Gran Canaria | Canary Islands | Gran Canaria | male | fledging | 1 | 1 | 1 | 0.924 |
| ERS16447390 | 278 | 2A444432 | *Fringilla* | *polatzeki* | – | 26/08/2020 | 27.93 | -15.69 | Gran Canaria | Canary Islands | Gran Canaria | male | adult | 1 | 1 | 1 | 0.923 |
| ERS16447385 | 289 | 2A444435 | *Fringilla* | *polatzeki* | – | 26/08/2020 | 27.93 | -15.69 | Gran Canaria | Canary Islands | Gran Canaria | male | fledging | 1 | 1 | 1 | 0.920 |
| ERS16447394 | 306 | 2A444426 | *Fringilla* | *polatzeki* | – | 26/08/2020 | 27.93 | -15.69 | Gran Canaria | Canary Islands | Gran Canaria | female | fledging | 1 | 1 | 0 | – |
| ERS16447380 | 315 | 2A444427 | *Fringilla* | *polatzeki* | – | 26/08/2020 | 27.93 | -15.69 | Gran Canaria | Canary Islands | Gran Canaria | female | fledging | 1 | 1 | 0 | – |
| ERS16447387 | 340 | 2A444429 | *Fringilla* | *polatzeki* | – | 26/08/2020 | 27.93 | -15.69 | Gran Canaria | Canary Islands | Gran Canaria | male | adult | 1 | 1 | 1 | 0.912 |
| ERS16447382 | 342 | 2A444424 | *Fringilla* | *polatzeki* | – | 26/08/2020 | 27.93 | -15.69 | Gran Canaria | Canary Islands | Gran Canaria | male | adult | 1 | 1 | 0 | – |
| ERS16447393 | 71 | 2A506482 | *Fringilla* | *polatzeki* | – | 01/09/2020 | 27.96 | -15.59 | Gran Canaria | Canary Islands | Gran Canaria | female | fledging | 1 | 1 | 1 | 0.922 |
| ERS16447391 | 74 | 2A506487 | *Fringilla* | *polatzeki* | – | 01/09/2020 | 27.96 | -15.59 | Gran Canaria | Canary Islands | Gran Canaria | male | fledging | 1 | 1 | 0 | – |
| ERS16447396 | 77 | 2A506483 | *Fringilla* | *polatzeki* | – | 01/09/2020 | 27.96 | -15.59 | Gran Canaria | Canary Islands | Gran Canaria | female | fledging | 1 | 1 | 1 | 0.919 |

Continued on next page

*Table 2 continued from previous page*

| Accession | Lab code | Ring | Genus | Species | Subsp. | Date | Lat. | Lon. | Region | Archipelago | Island/Mainland | Sex | Age | Microb. | Diet | WGS | $F_{ROH}$ |
|---|---|---|---|---|---|---|---|---|---|---|---|---|---|---|---|---|---|
| ERS16447408 | 245 | 2A493933 | *Fringilla* | *polatzeki* | – | – | 27.96 | -15.59 | – | Canary Islands | Gran Canaria | male | – | 0 | 1 | 0 | – |
| ERS16447405 | 249 | 2A444438 | *Fringilla* | *polatzeki* | – | – | 27.93 | -15.69 | – | Canary Islands | Gran Canaria | male | – | 0 | 1 | 0 | – |
| ERS16447406 | 250 | 2A444439 | *Fringilla* | *polatzeki* | – | – | 27.93 | -15.69 | – | Canary Islands | Gran Canaria | male | – | 0 | 1 | 0 | – |
| ERS16447407 | 251 | 2A444440 | *Fringilla* | *polatzeki* | – | – | 27.93 | -15.69 | – | Canary Islands | Gran Canaria | female | – | 0 | 1 | 0 | – |
| ERS16447400 | 270 | 2A444430 | *Fringilla* | *polatzeki* | – | – | 27.93 | -15.69 | – | Canary Islands | Gran Canaria | female | – | 0 | 1 | 0 | – |
| ERS16447409 | 274 | 2A493959 | *Fringilla* | *polatzeki* | – | – | 27.96 | -15.59 | – | Canary Islands | Gran Canaria | male | – | 0 | 1 | 0 | – |
| ERS16447379 | 277 | 2A444431 | *Fringilla* | *polatzeki* | – | – | 27.93 | -15.69 | – | Canary Islands | Gran Canaria | male | – | 0 | 1 | 0 | – |
| ERS16447401 | 283 | 2A444433 | *Fringilla* | *polatzeki* | – | – | 27.93 | -15.69 | – | Canary Islands | Gran Canaria | female | – | 0 | 1 | 0 | – |
| ERS16447398 | 285 | 2493932 | *Fringilla* | *polatzeki* | – | – | 27.93 | -15.69 | – | Canary Islands | Gran Canaria | male | – | 0 | 1 | 0 | – |
| ERS16447404 | 290 | 2A444437 | *Fringilla* | *polatzeki* | – | – | 27.93 | -15.69 | – | Canary Islands | Gran Canaria | female | – | 0 | 1 | 0 | – |
| ERS16447403 | 291 | 2A444436 | *Fringilla* | *polatzeki* | – | – | 27.93 | -15.69 | – | Canary Islands | Gran Canaria | female | – | 0 | 1 | 0 | – |
| ERS16447399 | 300 | 2595243 | *Fringilla* | *polatzeki* | – | – | 27.93 | -15.69 | – | Canary Islands | Gran Canaria | male | – | 0 | 1 | 0 | – |
| ERS16447410 | 68 | 2A506485 | *Fringilla* | *polatzeki* | – | – | 27.96 | -15.59 | – | Canary Islands | Gran Canaria | female | – | 0 | 1 | 0 | – |
| ERS16447411 | 72 | 2A506486 | *Fringilla* | *polatzeki* | – | – | 27.96 | -15.59 | – | Canary Islands | Gran Canaria | male | – | 0 | 1 | 0 | – |
| ERS16447213 | 116 | 4L23096 | *Fringilla* | *canariensis* | *canariensis* | 14/06/2020 | 28.14 | -17.28 | La Gomera | Canary Islands | La Gomera | female | adult | 1 | 1 | 0 | – |
| ERS16447225 | 117 | 4L23097 | *Fringilla* | *canariensis* | *canariensis* | 14/06/2020 | 28.14 | -17.28 | La Gomera | Canary Islands | La Gomera | male | adult | 1 | 1 | 1 | 0.711 |
| ERS16447219 | 118 | 4L23098 | *Fringilla* | *canariensis* | *canariensis* | 14/06/2020 | 28.14 | -17.28 | La Gomera | Canary Islands | La Gomera | male | fledging | 1 | 1 | 1 | 0.696 |
| ERS16447230 | 119 | 4L23099 | *Fringilla* | *canariensis* | *canariensis* | 14/06/2020 | 28.14 | -17.28 | La Gomera | Canary Islands | La Gomera | male | adult | 1 | 1 | 0 | – |
| ERS16447214 | 120 | 4L23100 | *Fringilla* | *canariensis* | *canariensis* | 14/06/2020 | 28.14 | -17.28 | La Gomera | Canary Islands | La Gomera | male | adult | 1 | 1 | 1 | 0.714 |
| ERS16447211 | 121 | 4L88842 | *Fringilla* | *canariensis* | *canariensis* | 14/06/2020 | 28.14 | -17.28 | La Gomera | Canary Islands | La Gomera | female | fledging | 1 | 1 | 0 | – |
| ERS16447218 | 124 | 4L88843 | *Fringilla* | *canariensis* | *canariensis* | 14/06/2020 | 28.14 | -17.28 | La Gomera | Canary Islands | La Gomera | female | fledging | 1 | 1 | 1 | 0.706 |
| ERS16447235 | 169 | 4L23078 | *Fringilla* | *canariensis* | *canariensis* | 12/06/2020 | 28.13 | -17.26 | La Gomera | Canary Islands | La Gomera | female | adult | 1 | 0 | 0 | – |
| ERS16447227 | 172 | 4L23073 | *Fringilla* | *canariensis* | *canariensis* | 11/06/2020 | 28.13 | -17.26 | La Gomera | Canary Islands | La Gomera | male | adult | 1 | 1 | 1 | 0.699 |
| ERS16447216 | 174 | 4L23082 | *Fringilla* | *canariensis* | *canariensis* | 12/06/2020 | 28.14 | -17.25 | La Gomera | Canary Islands | La Gomera | male | adult | 1 | 1 | 0 | – |
| ERS16447245 | 179 | 4L23075 | *Fringilla* | *canariensis* | *canariensis* | 11/06/2020 | 28.13 | -17.26 | La Gomera | Canary Islands | La Gomera | female | adult | 1 | 1 | 1 | 0.697 |
| ERS16447204 | 181 | 4L23079 | *Fringilla* | *canariensis* | *canariensis* | 12/06/2020 | 28.13 | -17.26 | La Gomera | Canary Islands | La Gomera | male | adult | 1 | 1 | 0 | – |
| ERS16447241 | 182 | 4L23080 | *Fringilla* | *canariensis* | *canariensis* | 12/06/2020 | 28.13 | -17.26 | La Gomera | Canary Islands | La Gomera | male | adult | 1 | 1 | 1 | 0.695 |
| ERS16447228 | 186 | 4L23081 | *Fringilla* | *canariensis* | *canariensis* | 12/06/2020 | 28.14 | -17.25 | La Gomera | Canary Islands | La Gomera | male | adult | 1 | 1 | 1 | 0.706 |
| ERS16447232 | 55 | 4L23083 | *Fringilla* | *canariensis* | *canariensis* | 13/06/2020 | 28.13 | -17.22 | La Gomera | Canary Islands | La Gomera | male | adult | 1 | 1 | 0 | – |
| ERS16447208 | 56 | 4L23084 | *Fringilla* | *canariensis* | *canariensis* | 13/06/2020 | 28.13 | -17.22 | La Gomera | Canary Islands | La Gomera | male | adult | 1 | 1 | 0 | – |
| ERS16447223 | 58 | 4L23085 | *Fringilla* | *canariensis* | *canariensis* | 13/06/2020 | 28.13 | -17.22 | La Gomera | Canary Islands | La Gomera | male | adult | 1 | 1 | 1 | 0.713 |
| ERS16447217 | 59 | 4L23087 | *Fringilla* | *canariensis* | *canariensis* | 13/06/2020 | 28.13 | -17.22 | La Gomera | Canary Islands | La Gomera | male | fledging | 1 | 1 | 0 | – |
| ERS16447210 | 60 | 4L23088 | *Fringilla* | *canariensis* | *canariensis* | 13/06/2020 | 28.13 | -17.22 | La Gomera | Canary Islands | La Gomera | male | adult | 1 | 1 | 0 | – |
| ERS16447220 | 62 | 4L23090 | *Fringilla* | *canariensis* | *canariensis* | 13/06/2020 | 28.13 | -17.22 | La Gomera | Canary Islands | La Gomera | male | adult | 1 | 0 | 0 | – |
| ERS16447236 | 63 | 4L23091 | *Fringilla* | *canariensis* | *canariensis* | 13/06/2020 | 28.13 | -17.22 | La Gomera | Canary Islands | La Gomera | female | fledging | 1 | 1 | 1 | 0.706 |
| ERS16447212 | 64 | 4L23092 | *Fringilla* | *canariensis* | *canariensis* | 13/06/2020 | 28.14 | -17.22 | La Gomera | Canary Islands | La Gomera | male | adult | 1 | 1 | 0 | – |
| ERS16447233 | 67 | 4L23093 | *Fringilla* | *canariensis* | *canariensis* | 14/06/2020 | 28.15 | -17.24 | La Gomera | Canary Islands | La Gomera | male | adult | 1 | 1 | 0 | – |
| ERS16447207 | 85 | 4L88826 | *Fringilla* | *canariensis* | *canariensis* | 03/02/2018 | 28.13 | -17.22 | La Gomera | Canary Islands | La Gomera | male | adult | 1 | 1 | 0 | – |
| ERS16447215 | 86 | 4L88825 | *Fringilla* | *canariensis* | *canariensis* | 03/02/2018 | 28.13 | -17.22 | La Gomera | Canary Islands | La Gomera | male | adult | 1 | 1 | 0 | – |
| ERS16447209 | 87 | 4L88827 | *Fringilla* | *canariensis* | *canariensis* | 04/02/2018 | 28.13 | -17.22 | La Gomera | Canary Islands | La Gomera | male | adult | 1 | 0 | 1 | 0.704 |

*Continued on next page*

*Table 2 continued from previous page*

| Accession | Lab code | Ring | Genus | Species | Subsp. | Date | Lat. | Lon. | Region | Archipelago | Island/Mainland | Sex | Age | Microb. | Diet | WGS | $F_{ROH}$ |
|---|---|---|---|---|---|---|---|---|---|---|---|---|---|---|---|---|---|
| ERS16447203 | 109 | 4L23094 | *Fringilla* | *canariensis* | *canariensis* | – | 28.15 | -17.24 | – | Canary Islands | La Gomera | male | – | 0 | 1 | 0 | – |
| ERS16447252 | 110 | 4L23095 | *Fringilla* | *canariensis* | *canariensis* | – | 28.15 | -17.24 | – | Canary Islands | La Gomera | male | – | 0 | 1 | 0 | – |
| ERS16447246 | 176 | 4L23071 | *Fringilla* | *canariensis* | *canariensis* | – | 28.14 | -17.25 | – | Canary Islands | La Gomera | female | – | 0 | 1 | 0 | – |
| ERS16447248 | 178 | 4L23076 | *Fringilla* | *canariensis* | *canariensis* | – | 28.13 | -17.26 | – | Canary Islands | La Gomera | male | – | 0 | 1 | 0 | – |
| ERS16447249 | 180 | 4L23077 | *Fringilla* | *canariensis* | *canariensis* | – | 28.13 | -17.26 | – | Canary Islands | La Gomera | female | – | 0 | 1 | 0 | – |
| ERS16447247 | 183 | 4L23074 | *Fringilla* | *canariensis* | *canariensis* | – | 28.13 | -17.26 | – | Canary Islands | La Gomera | male | – | 0 | 1 | 0 | – |
| ERS16447250 | 57 | 4L23086 | *Fringilla* | *canariensis* | *canariensis* | – | 28.13 | -17.22 | – | Canary Islands | La Gomera | male | – | 0 | 1 | 0 | – |
| ERS16447251 | 61 | 4L23089 | *Fringilla* | *canariensis* | *canariensis* | – | 28.13 | -17.22 | – | Canary Islands | La Gomera | male | – | 0 | 1 | 0 | – |
| ERS16447253 | 84 | 4L88824 | *Fringilla* | *canariensis* | *canariensis* | – | 28.13 | -17.22 | – | Canary Islands | La Gomera | male | – | 0 | 1 | 0 | – |
| ERS16447360 | 296 | 4L88864 | *Fringilla* | *canariensis* | *palmae* | 29/05/2021 | 28.52 | -17.83 | La Palma | Canary Islands | La Palma | female | adult | 1 | 1 | 1 | 0.773 |
| ERS16447365 | 308 | 4L88848 | *Fringilla* | *canariensis* | *palmae* | 25/05/2021 | 28.72 | -17.77 | La Palma | Canary Islands | La Palma | male | adult | 1 | 1 | 1 | 0.768 |
| ERS16447366 | 310 | 4L88849 | *Fringilla* | *canariensis* | *palmae* | 25/05/2021 | 28.77 | -17.91 | La Palma | Canary Islands | La Palma | male | adult | 1 | 1 | 1 | 0.783 |
| ERS16447359 | 311 | 4L88850 | *Fringilla* | *canariensis* | *palmae* | 25/05/2021 | 28.81 | -17.91 | La Palma | Canary Islands | La Palma | male | adult | 1 | 1 | 1 | 0.765 |
| ERS16447368 | 362 | 4L88851 | *Fringilla* | *canariensis* | *palmae* | 25/05/2021 | 28.80 | -17.91 | La Palma | Canary Islands | La Palma | male | adult | 1 | 1 | 1 | 0.776 |
| ERS16447357 | 367 | 4L88853 | *Fringilla* | *canariensis* | *palmae* | 26/05/2021 | 28.61 | -17.83 | La Palma | Canary Islands | La Palma | male | adult | 1 | 1 | 1 | 0.772 |
| ERS16447358 | 387 | 4L88854 | *Fringilla* | *canariensis* | *palmae* | 26/05/2021 | 28.64 | -17.83 | La Palma | Canary Islands | La Palma | male | adult | 1 | 1 | 0 | – |
| ERS16447361 | 388 | 4L88855 | *Fringilla* | *canariensis* | *palmae* | 27/05/2021 | 28.81 | -17.81 | La Palma | Canary Islands | La Palma | male | adult | 1 | 1 | 0 | – |
| ERS16447367 | 392 | 4L88857 | *Fringilla* | *canariensis* | *palmae* | 27/05/2021 | 28.78 | -17.92 | La Palma | Canary Islands | La Palma | male | adult | 1 | 1 | 1 | 0.773 |
| ERS16447371 | 409 | 4L88859 | *Fringilla* | *canariensis* | *palmae* | 27/05/2021 | 28.77 | -17.90 | La Palma | Canary Islands | La Palma | male | adult | 1 | 1 | 0 | – |
| ERS16447369 | 410 | 4L88860 | *Fringilla* | *canariensis* | *palmae* | 29/05/2021 | 28.52 | -17.83 | La Palma | Canary Islands | La Palma | male | adult | 1 | 1 | 1 | 0.784 |
| ERS16447364 | 412 | 4L88869 | *Fringilla* | *canariensis* | *palmae* | 29/05/2021 | 28.52 | -17.83 | La Palma | Canary Islands | La Palma | male | adult | 1 | 1 | 1 | 0.782 |
| ERS16447370 | 413 | 4L88865 | *Fringilla* | *canariensis* | *palmae* | 29/05/2021 | 28.52 | -17.83 | La Palma | Canary Islands | La Palma | female | adult | 1 | 1 | 0 | – |
| ERS16447362 | 414 | 4L88863 | *Fringilla* | *canariensis* | *palmae* | 29/05/2021 | 28.52 | -17.83 | La Palma | Canary Islands | La Palma | male | adult | 1 | 1 | 1 | 0.775 |
| ERS16447363 | 97 | 4L88867 | *Fringilla* | *canariensis* | *palmae* | 29/05/2021 | 28.52 | -17.83 | La Palma | Canary Islands | La Palma | male | adult | 1 | 1 | 1 | 0.781 |
| ERS16447355 | 297 | 4L88847 | *Fringilla* | *canariensis* | *palmae* | – | 28.72 | -17.78 | – | Canary Islands | La Palma | male | – | 0 | 1 | 0 | – |
| ERS16447374 | 298 | 4L88861 | *Fringilla* | *canariensis* | *palmae* | – | 28.52 | -17.83 | – | Canary Islands | La Palma | female | – | 0 | 1 | 0 | – |
| ERS16447372 | 318 | 4L88856 | *Fringilla* | *canariensis* | *palmae* | – | 28.79 | -17.92 | – | Canary Islands | La Palma | male | – | 0 | 1 | 0 | – |
| ERS16447356 | 363 | 4L88852 | *Fringilla* | *canariensis* | *palmae* | – | 28.62 | -17.82 | – | Canary Islands | La Palma | male | – | 0 | 1 | 0 | – |
| ERS16447373 | 391 | 4L88858 | *Fringilla* | *canariensis* | *palmae* | – | 28.78 | -17.92 | – | Canary Islands | La Palma | male | – | 0 | 1 | 0 | – |
| ERS16447375 | 416 | 4L88862 | *Fringilla* | *canariensis* | *palmae* | – | 28.52 | -17.83 | – | Canary Islands | La Palma | female | – | 0 | 1 | 0 | – |
| ERS16447377 | 417 | 4L88871 | *Fringilla* | *canariensis* | *palmae* | – | 28.52 | -17.83 | – | Canary Islands | La Palma | male | – | 0 | 1 | 0 | – |
| ERS16447376 | 96 | 4L88868 | *Fringilla* | *canariensis* | *palmae* | – | 28.52 | -17.83 | – | Canary Islands | La Palma | female | – | 0 | 1 | 0 | – |
| ERS16447238 | 132 | 4L88968 | *Fringilla* | *canariensis* | *canariensis* | 14/05/2020 | 28.32 | -16.84 | Tenerife | Canary Islands | Tenerife | male | adult | 1 | 1 | 0 | 0.811 |
| ERS16447244 | 135 | 4L88978 | *Fringilla* | *canariensis* | *canariensis* | 14/05/2020 | 28.32 | -16.84 | Tenerife | Canary Islands | Tenerife | female | fledging | 1 | 1 | 0 | – |
| ERS16447231 | 145 | 2L55185 | *Fringilla* | *canariensis* | *canariensis* | 19/02/2020 | 28.41 | -16.43 | Tenerife | Canary Islands | Tenerife | male | adult | 1 | 1 | 1 | 0.784 |
| ERS16447229 | 153 | 2L55194 | *Fringilla* | *canariensis* | *canariensis* | 20/02/2020 | 28.46 | -16.40 | Tenerife | Canary Islands | Tenerife | male | adult | 1 | 1 | 1 | 0.781 |
| ERS16447224 | 156 | 2L55191 | *Fringilla* | *canariensis* | *canariensis* | 19/02/2020 | 28.41 | -16.43 | Tenerife | Canary Islands | Tenerife | male | adult | 1 | 1 | 1 | 0.779 |
| ERS16447242 | 157 | 4L88958 | *Fringilla* | *canariensis* | *canariensis* | 26/02/2020 | 28.53 | -16.24 | Tenerife | Canary Islands | Tenerife | female | adult | 1 | 1 | 1 | 0.786 |
| ERS16447234 | 159 | 4L88960 | *Fringilla* | *canariensis* | *canariensis* | 26/02/2020 | 28.54 | -16.31 | Tenerife | Canary Islands | Tenerife | male | adult | 1 | 1 | 1 | 0.776 |
| ERS16447205 | 160 | 4L88961 | *Fringilla* | *canariensis* | *canariensis* | 27/02/2020 | 28.53 | -16.31 | Tenerife | Canary Islands | Tenerife | male | adult | 1 | 1 | 0 | – |

*Continued on next page*

*Table 2 continued from previous page*

| Accession | Lab code | Ring | Genus | Species | Subsp. | Date | Lat. | Lon. | Region | Archipelago | Island/Mainland | Sex | Age | Microb. | Diet | WGS | $F_{ROH}$ |
|---|---|---|---|---|---|---|---|---|---|---|---|---|---|---|---|---|---|
| ERS16447240 | 162 | 4L88963 | *Fringilla* | *canariensis* | *canariensis* | 27/02/2020 | 28.54 | -16.31 | Tenerife | Canary Islands | Tenerife | male | adult | 1 | 1 | 0 | – |
| ERS16447226 | 44 | 4L88964 | *Fringilla* | *canariensis* | *canariensis* | 11/05/2020 | 28.54 | -16.31 | Tenerife | Canary Islands | Tenerife | male | adult | 1 | 1 | 1 | 0.786 |
| ERS16447206 | 45 | 4L88965 | *Fringilla* | *canariensis* | *canariensis* | 14/05/2020 | 28.32 | -16.84 | Tenerife | Canary Islands | Tenerife | male | adult | 1 | 1 | 0 | – |
| ERS16447222 | 49 | 4L88972 | *Fringilla* | *canariensis* | *canariensis* | 14/05/2020 | 28.32 | -16.84 | Tenerife | Canary Islands | Tenerife | male | adult | 1 | 1 | 0 | – |
| ERS16447221 | 50 | 4L88966 | *Fringilla* | *canariensis* | *canariensis* | 14/05/2020 | 28.32 | -16.84 | Tenerife | Canary Islands | Tenerife | female | adult | 1 | 1 | 1 | 0.791 |
| ERS16447239 | 51 | 4L88971 | *Fringilla* | *canariensis* | *canariensis* | 14/05/2020 | 28.32 | -16.84 | Tenerife | Canary Islands | Tenerife | male | adult | 1 | 1 | 1 | 0.800 |
| ERS16447237 | 53 | 4L88979 | *Fringilla* | *canariensis* | *canariensis* | 14/05/2020 | 28.32 | -16.84 | Tenerife | Canary Islands | Tenerife | female | fledging | 1 | 1 | 0 | – |
| ERS16447243 | 54 | 4L88974 | *Fringilla* | *canariensis* | *canariensis* | 14/05/2020 | 28.32 | -16.84 | Tenerife | Canary Islands | Tenerife | male | adult | 1 | 1 | 1 | 0.780 |
| ERS16447255 | 131 | 4L88975 | *Fringilla* | *canariensis* | *canariensis* | – | 28.32 | -16.84 | – | Canary Islands | Tenerife | female | – | 0 | 1 | 0 | – |
| ERS16447256 | 133 | 4L88976 | *Fringilla* | *canariensis* | *canariensis* | – | 28.32 | -16.84 | – | Canary Islands | Tenerife | male | – | 0 | 1 | 0 | – |
| ERS16447257 | 134 | 4L88977 | *Fringilla* | *canariensis* | *canariensis* | – | 28.32 | -16.84 | – | Canary Islands | Tenerife | male | – | 0 | 1 | 0 | – |
| ERS16447201 | 158 | 4L88959 | *Fringilla* | *canariensis* | *canariensis* | – | 28.53 | -16.24 | – | Canary Islands | Tenerife | male | – | 0 | 1 | 0 | – |
| ERS16447202 | 48 | 4L88970 | *Fringilla* | *canariensis* | *canariensis* | – | 28.32 | -16.84 | – | Canary Islands | Tenerife | male | – | 0 | 1 | 0 | – |
| ERS16447254 | 52 | 4L88967 | *Fringilla* | *canariensis* | *canariensis* | – | 28.32 | -16.84 | – | Canary Islands | Tenerife | male | – | 0 | 1 | 0 | – |
| ERS16447458 | 129 | 2A444422 | *Fringilla* | *teydea* | – | 03/06/2020 | 28.39 | -16.46 | Tenerife | Canary Islands | Tenerife | male | adult | 1 | 1 | 1 | 0.926 |
| ERS16447461 | 144 | 2L55184 | *Fringilla* | *teydea* | – | 19/02/2020 | 28.41 | -16.43 | Tenerife | Canary Islands | Tenerife | male | adult | 1 | 1 | 1 | 0.927 |
| ERS16447463 | 146 | 2L55186 | *Fringilla* | *teydea* | – | 19/02/2020 | 28.41 | -16.43 | Tenerife | Canary Islands | Tenerife | female | adult | 1 | 1 | 1 | 0.922 |
| ERS16447460 | 147 | 2L55187 | *Fringilla* | *teydea* | – | 19/02/2020 | 28.41 | -16.43 | Tenerife | Canary Islands | Tenerife | male | adult | 1 | 1 | 1 | 0.926 |
| ERS16447462 | 149 | 2A357348 | *Fringilla* | *teydea* | – | 19/02/2020 | 28.41 | -16.43 | Tenerife | Canary Islands | Tenerife | female | adult | 1 | 1 | 1 | 0.921 |
| ERS16447457 | 150 | 2L55189 | *Fringilla* | *teydea* | – | 19/02/2020 | 28.41 | -16.43 | Tenerife | Canary Islands | Tenerife | male | adult | 1 | 1 | 1 | 0.925 |
| ERS16447459 | 151 | 2L55190 | *Fringilla* | *teydea* | – | 19/02/2020 | 28.41 | -16.43 | Tenerife | Canary Islands | Tenerife | male | adult | 1 | 1 | 1 | 0.924 |
| ERS16447464 | 154 | 2L55192 | *Fringilla* | *teydea* | – | 19/02/2020 | 28.41 | -16.43 | Tenerife | Canary Islands | Tenerife | female | adult | 1 | 1 | 1 | 0.921 |
| ERS16447465 | 163 | 2L55195 | *Fringilla* | *teydea* | – | 29/02/2020 | 28.31 | -16.60 | Tenerife | Canary Islands | Tenerife | male | adult | 1 | 1 | 1 | 0.927 |
| ERS16447467 | 130 | 2A444423 | *Fringilla* | *teydea* | – | – | 28.39 | -16.46 | – | Canary Islands | Tenerife | female | – | 0 | 1 | 0 | – |
| ERS16447468 | 148 | 2L55188 | *Fringilla* | *teydea* | – | – | 28.41 | -16.43 | – | Canary Islands | Tenerife | female | – | 0 | 1 | 0 | – |
| ERS16447466 | 152 | 2A357396 | *Fringilla* | *teydea* | – | – | 28.41 | -16.43 | – | Canary Islands | Tenerife | female | – | 0 | 1 | 0 | – |

Table S3: **PERMANOVA results for gut microbiome composition.** Tests were performed using Bray-Curtis, Jaccard, weighted UniFrac, and unweighted UniFrac distance matrices as response variables, and host population (Species_Label), age, and sex as predictors. Significant terms ($p < 0.05$) shown in bold.

| Distance metric | Taxonomic level | Predictor | $R^2$ | $F$ | $p$-value |
|---|---|---|---|---|---|
| **Jaccard** | **Phylum** | **Species_Label** | **0.132** | **3.362** | **<0.001** |
| **Jaccard** | **Phylum** | **Age_cat** | **0.008** | **2.068** | **0.049** |
| Jaccard | Phylum | Sex | 0.006 | 1.616 | 0.122 |
| **Bray-Curtis** | **Phylum** | **Species_Label** | **0.199** | **5.471** | **<0.001** |
| Bray-Curtis | Phylum | Age_cat | 0.002 | 0.494 | 0.774 |
| Bray-Curtis | Phylum | Sex | 0.002 | 0.553 | 0.724 |
| **Unweighted UniFrac** | **Phylum** | **Species_Label** | **0.128** | **3.231** | **<0.001** |
| Unweighted UniFrac | Phylum | Age_cat | 0.008 | 1.955 | 0.065 |
| Unweighted UniFrac | Phylum | Sex | 0.006 | 1.543 | 0.15 |
| **Weighted UniFrac** | **Phylum** | **Species_Label** | **0.224** | **6.336** | **<0.001** |
| Weighted UniFrac | Phylum | Age_cat | 0.002 | 0.546 | 0.691 |
| Weighted UniFrac | Phylum | Sex | 0.001 | 0.387 | 0.822 |
| **Jaccard** | **Class** | **Species_Label** | **0.129** | **3.252** | **<0.001** |
| **Jaccard** | **Class** | **Age_cat** | **0.007** | **1.83** | **0.043** |
| Jaccard | Class | Sex | 0.006 | 1.462 | 0.122 |
| **Bray-Curtis** | **Class** | **Species_Label** | **0.195** | **5.375** | **<0.001** |
| Bray-Curtis | Class | Age_cat | 0.001 | 0.366 | 0.899 |
| Bray-Curtis | Class | Sex | 0.002 | 0.646 | 0.675 |
| **Unweighted UniFrac** | **Class** | **Species_Label** | **0.125** | **3.14** | **<0.001** |
| Unweighted UniFrac | Class | Age_cat | 0.007 | 1.852 | 0.051 |
| Unweighted UniFrac | Class | Sex | 0.006 | 1.52 | 0.121 |
| **Weighted UniFrac** | **Class** | **Species_Label** | **0.217** | **6.125** | **<0.001** |
| Weighted UniFrac | Class | Age_cat | 0.002 | 0.482 | 0.77 |
| Weighted UniFrac | Class | Sex | 0.002 | 0.442 | 0.804 |
| **Jaccard** | **Order** | **Species_Label** | **0.126** | **3.165** | **<0.001** |

*Continued on next page*

*Table 3 continued from previous page*

| Distance metric | Taxonomic level | Predictor | $R^2$ | $F$ | $p$-value |
|---|---|---|---|---|---|
| Jaccard | Order | Age_cat | 0.005 | 1.135 | 0.26 |
| Jaccard | Order | Sex | 0.004 | 1.127 | 0.269 |
| **Bray-Curtis** | **Order** | **Species_Label** | **0.159** | **4.198** | **<0.001** |
| Bray-Curtis | Order | Age_cat | 0.003 | 0.851 | 0.548 |
| Bray-Curtis | Order | Sex | 0.003 | 0.874 | 0.525 |
| **Unweighted UniFrac** | **Order** | **Species_Label** | **0.129** | **3.241** | **<0.001** |
| Unweighted UniFrac | Order | Age_cat | 0.005 | 1.282 | 0.161 |
| Unweighted UniFrac | Order | Sex | 0.005 | 1.17 | 0.242 |
| **Weighted UniFrac** | **Order** | **Species_Label** | **0.187** | **5.085** | **<0.001** |
| Weighted UniFrac | Order | Age_cat | 0.003 | 0.87 | 0.506 |
| Weighted UniFrac | Order | Sex | 0.004 | 0.965 | 0.427 |
| **Jaccard** | **Family** | **Species_Label** | **0.122** | **3.029** | **<0.001** |
| Jaccard | Family | Age_cat | 0.004 | 1.066 | 0.318 |
| Jaccard | Family | Sex | 0.004 | 0.962 | 0.513 |
| **Bray-Curtis** | **Family** | **Species_Label** | **0.171** | **4.576** | **<0.001** |
| Bray-Curtis | Family | Age_cat | 0.003 | 0.739 | 0.735 |
| Bray-Curtis | Family | Sex | 0.004 | 1.031 | 0.377 |
| **Unweighted UniFrac** | **Family** | **Species_Label** | **0.127** | **3.173** | **<0.001** |
| Unweighted UniFrac | Family | Age_cat | 0.005 | 1.218 | 0.169 |
| Unweighted UniFrac | Family | Sex | 0.004 | 1.031 | 0.383 |
| **Weighted UniFrac** | **Family** | **Species_Label** | **0.19** | **5.192** | **<0.001** |
| Weighted UniFrac | Family | Age_cat | 0.003 | 0.795 | 0.593 |
| Weighted UniFrac | Family | Sex | 0.003 | 0.92 | 0.473 |
| **Jaccard** | **Genus** | **Species_Label** | **0.11** | **2.685** | **<0.001** |
| Jaccard | Genus | Age_cat | 0.004 | 0.998 | 0.446 |
| Jaccard | Genus | Sex | 0.004 | 0.991 | 0.458 |
| **Bray-Curtis** | **Genus** | **Species_Label** | **0.157** | **4.128** | **<0.001** |
| Bray-Curtis | Genus | Age_cat | 0.004 | 0.952 | 0.468 |

*Continued on next page*

*Table 3 continued from previous page*

| Distance metric | Taxonomic level | Predictor | $R^2$ | $F$ | $p$-value |
|---|---|---|---|---|---|
| Bray-Curtis | Genus | Sex | 0.005 | 1.311 | 0.123 |
| **Unweighted UniFrac** | **Genus** | **Species_Label** | **0.12** | **2.989** | **<0.001** |
| Unweighted UniFrac | Genus | Age_cat | 0.004 | 1.082 | 0.282 |
| Unweighted UniFrac | Genus | Sex | 0.004 | 0.992 | 0.437 |
| **Weighted UniFrac** | **Genus** | **Species_Label** | **0.187** | **5.068** | **<0.001** |
| Weighted UniFrac | Genus | Age_cat | 0.003 | 0.816 | 0.59 |
| Weighted UniFrac | Genus | Sex | 0.004 | 1.013 | 0.384 |
| **Jaccard** | **Species** | **Species_Label** | **0.104** | **2.543** | **<0.001** |
| Jaccard | Species | Age_cat | 0.004 | 0.976 | 0.527 |
| Jaccard | Species | Sex | 0.004 | 0.982 | 0.495 |
| **Bray-Curtis** | **Species** | **Species_Label** | **0.157** | **4.115** | **<0.001** |
| Bray-Curtis | Species | Age_cat | 0.004 | 0.93 | 0.512 |
| Bray-Curtis | Species | Sex | 0.005 | 1.21 | 0.168 |
| **Unweighted UniFrac** | **Species** | **Species_Label** | **0.117** | **2.909** | **<0.001** |
| Unweighted UniFrac | Species | Age_cat | 0.004 | 1.08 | 0.277 |
| Unweighted UniFrac | Species | Sex | 0.004 | 0.996 | 0.434 |
| **Weighted UniFrac** | **Species** | **Species_Label** | **0.186** | **5.061** | **<0.001** |
| Weighted UniFrac | Species | Age_cat | 0.003 | 0.824 | 0.583 |
| Weighted UniFrac | Species | Sex | 0.004 | 1.013 | 0.384 |
| **Jaccard** | **OTU99** | **Species_Label** | **0.081** | **1.91** | **<0.001** |
| Jaccard | OTU99 | Age_cat | 0.004 | 1.047 | 0.172 |
| Jaccard | OTU99 | Sex | 0.004 | 1.031 | 0.257 |
| **Bray-Curtis** | **OTU99** | **Species_Label** | **0.134** | **3.393** | **<0.001** |
| Bray-Curtis | OTU99 | Age_cat | 0.005 | 1.236 | 0.121 |
| Bray-Curtis | OTU99 | Sex | 0.004 | 1.047 | 0.3 |
| **Unweighted UniFrac** | **OTU99** | **Species_Label** | **0.11** | **2.704** | **<0.001** |
| Unweighted UniFrac | OTU99 | Age_cat | 0.004 | 1.061 | 0.285 |
| Unweighted UniFrac | OTU99 | Sex | 0.004 | 1.062 | 0.285 |

*Continued on next page*

*Table 3 continued from previous page*

| Distance metric | Taxonomic level | Predictor | $R^2$ | $F$ | $p$-value |
|---|---|---|---|---|---|
| **Weighted UniFrac** | **OTU99** | **Species_Label** | **0.184** | **4.988** | **<0.001** |
| Weighted UniFrac | OTU99 | Age_cat | 0.003 | 0.866 | 0.536 |
| Weighted UniFrac | OTU99 | Sex | 0.004 | 1.019 | 0.376 |

Table S4: **Generalised linear model (GLM) results for microbiome richness across populations.** The mainland population (*Fringilla coelebs*) is used as the reference group. Both observed richness and Faith's phylogenetic diversity are reported at each taxonomic level. Significant terms ($p < 0.05$) shown in bold.

| Alpha diversity metric | Taxonomic level | Predictor | Estimate | Std. error | $z$ | Pr($>|z|$) | CI 95% lo | CI 95% hi |
|---|---|---|---|---|---|---|---|---|
| **Richness** | **Phylum** | **(Intercept)** | **7.167** | **0.082** | **24.164** | **<0.0001** | **6.088** | **8.382** |
| Richness | Phylum | São Miguel *F. moreletti* | 6.25 | 0.114 | -1.197 | 0.2312 | 4.995 | 7.821 |
| Richness | Phylum | Flores *F. moreletti* | 7.962 | 0.111 | 0.951 | 0.3417 | 6.412 | 9.896 |
| Richness | Phylum | Madeira *F. maderensis* | 6.762 | 0.121 | -0.481 | 0.6308 | 5.33 | 8.567 |
| Richness | Phylum | El Hierro *F. c. ombriosa* | 7.478 | 0.115 | 0.369 | 0.7123 | 5.963 | 9.378 |
| Richness | Phylum | La Palma *F. c. palmae* | 7.4 | 0.13 | 0.246 | 0.8058 | 5.721 | 9.539 |
| **Richness** | **Phylum** | **Gran Canaria *F. c. bakeri*** | **9.607** | **0.105** | **2.785** | **0.0053** | **7.825** | **11.822** |
| Richness | Phylum | Tenerife *F. c. canariensis* | 8.875 | 0.122 | 1.75 | 0.0801 | 6.979 | 11.271 |
| **Richness** | **Phylum** | **La Gomera *F. c. canariensis*** | **9.615** | **0.107** | **2.752** | **0.0059** | **7.806** | **11.868** |
| **Richness** | **Phylum** | **Tenerife *F. teydea*** | **12** | **0.135** | **3.83** | **0.0001** | **9.2** | **15.599** |
| **Richness** | **Phylum** | **Gran Canaria *F. polatzeki*** | **9.667** | **0.116** | **2.576** | **0.01** | **7.698** | **12.141** |
| **Faith's PD** | **Phylum** | **(Intercept)** | **3.333** | **0.112** | **10.768** | **<0.0001** | **2.655** | **4.118** |
| Faith's PD | Phylum | São Miguel *F. moreletti* | 2.929 | 0.157 | -0.824 | 0.4101 | 2.151 | 3.988 |
| Faith's PD | Phylum | Flores *F. moreletti* | 3.615 | 0.152 | 0.534 | 0.5934 | 2.685 | 4.88 |
| Faith's PD | Phylum | Madeira *F. maderensis* | 3.048 | 0.168 | -0.534 | 0.5931 | 2.188 | 4.228 |
| Faith's PD | Phylum | El Hierro *F. c. ombriosa* | 3.435 | 0.159 | 0.189 | 0.8501 | 2.515 | 4.69 |
| Faith's PD | Phylum | La Palma *F. c. palmae* | 3.333 | 0.18 | 0 | 1 | 2.328 | 4.728 |
| Faith's PD | Phylum | Gran Canaria *F. c. bakeri* | 4.357 | 0.144 | 1.862 | 0.0626 | 3.294 | 5.795 |
| Faith's PD | Phylum | Tenerife *F. c. canariensis* | 4.063 | 0.167 | 1.185 | 0.2362 | 2.921 | 5.629 |
| Faith's PD | Phylum | La Gomera *F. c. canariensis* | 4.308 | 0.146 | 1.752 | 0.0798 | 3.239 | 5.755 |
| **Faith's PD** | **Phylum** | **Tenerife *F. teydea*** | **5.556** | **0.18** | **2.833** | **0.0046** | **3.879** | **7.879** |
| Faith's PD | Phylum | Gran Canaria *F. polatzeki* | 4.278 | 0.16 | 1.563 | 0.1182 | 3.125 | 5.851 |
| **Richness** | **Class** | **(Intercept)** | **9.958** | **0.091** | **25.289** | **<0.0001** | **8.332** | **11.901** |
| Richness | Class | São Miguel *F. moreletti* | 8.107 | 0.127 | -1.618 | 0.1057 | 6.317 | 10.399 |
| Richness | Class | Flores *F. moreletti* | 11.192 | 0.124 | 0.94 | 0.3475 | 8.772 | 14.284 |
| Richness | Class | Madeira *F. maderensis* | 9.333 | 0.134 | -0.483 | 0.6292 | 7.173 | 12.142 |

*Continued on next page*

*Table 4 continued from previous page*

| Alpha diversity metric | Taxonomic level | Predictor | Estimate | Std. error | $z$ | Pr(> \|$z$\|) | CI 95% lo | CI 95% hi |
|---|---|---|---|---|---|---|---|---|
| Richness | Class | El Hierro *F. c. ombriosa* | 11.043 | 0.128 | 0.806 | 0.42 | 8.589 | 14.203 |
| Richness | Class | La Palma *F. c. palmae* | 10.867 | 0.145 | 0.604 | 0.5461 | 8.185 | 14.432 |
| **Richness** | **Class** | **Gran Canaria *F. c. bakeri*** | **14.5** | **0.119** | **3.151** | **0.0016** | **11.481** | **18.324** |
| Richness | Class | Tenerife *F. c. canariensis* | 12.188 | 0.14 | 1.446 | 0.148 | 9.272 | 16.033 |
| **Richness** | **Class** | **La Gomera *F. c. canariensis*** | **14.731** | **0.121** | **3.237** | **0.0012** | **11.625** | **18.68** |
| **Richness** | **Class** | **Tenerife *F. teydea*** | **17.778** | **0.159** | **3.638** | **0.0003** | **13.033** | **24.348** |
| **Richness** | **Class** | **Gran Canaria *F. polatzeki*** | **14.778** | **0.132** | **2.988** | **0.0028** | **11.412** | **19.159** |
| **Faith's PD** | **Class** | **(Intercept)** | **4.458** | **0.097** | **15.461** | **<0.0001** | **3.666** | **5.357** |
| Faith's PD | Class | São Miguel *F. moreletti* | 3.679 | 0.138 | -1.393 | 0.1637 | 2.805 | 4.822 |
| Faith's PD | Class | Flores *F. moreletti* | 4.846 | 0.131 | 0.634 | 0.5258 | 3.747 | 6.278 |
| Faith's PD | Class | Madeira *F. maderensis* | 3.952 | 0.146 | -0.824 | 0.4102 | 2.961 | 5.258 |
| Faith's PD | Class | El Hierro *F. c. ombriosa* | 4.739 | 0.136 | 0.449 | 0.6536 | 3.629 | 6.191 |
| Faith's PD | Class | La Palma *F. c. palmae* | 4.733 | 0.153 | 0.391 | 0.6958 | 3.494 | 6.374 |
| **Faith's PD** | **Class** | **Gran Canaria *F. c. bakeri*** | **6.321** | **0.122** | **2.851** | **0.0044** | **4.983** | **8.058** |
| Faith's PD | Class | Tenerife *F. c. canariensis* | 5.25 | 0.146 | 1.121 | 0.2622 | 3.937 | 6.978 |
| **Faith's PD** | **Class** | **La Gomera *F. c. canariensis*** | **6.154** | **0.125** | **2.581** | **0.0099** | **4.826** | **7.879** |
| **Faith's PD** | **Class** | **Tenerife *F. teydea*** | **7.222** | **0.157** | **3.067** | **0.0022** | **5.283** | **9.798** |
| **Faith's PD** | **Class** | **Gran Canaria *F. polatzeki*** | **6.444** | **0.134** | **2.749** | **0.006** | **4.956** | **8.388** |
| **Richness** | **Order** | **(Intercept)** | **19.333** | **0.1** | **29.623** | **<0.0001** | **15.948** | **23.609** |
| Richness | Order | São Miguel *F. moreletti* | 15.393 | 0.138 | -1.652 | 0.0986 | 11.736 | 20.165 |
| Richness | Order | Flores *F. moreletti* | 20.346 | 0.138 | 0.369 | 0.712 | 15.508 | 26.679 |
| Richness | Order | Madeira *F. maderensis* | 17.571 | 0.147 | -0.649 | 0.5162 | 13.171 | 23.465 |
| Richness | Order | El Hierro *F. c. ombriosa* | 21.739 | 0.142 | 0.826 | 0.409 | 16.455 | 28.731 |
| Richness | Order | La Palma *F. c. palmae* | 21.733 | 0.16 | 0.731 | 0.4647 | 15.913 | 29.819 |
| **Richness** | **Order** | **Gran Canaria *F. c. bakeri*** | **29.25** | **0.134** | **3.091** | **0.002** | **22.481** | **38.021** |
| Richness | Order | Tenerife *F. c. canariensis* | 23.688 | 0.156 | 1.3 | 0.1935 | 17.469 | 32.246 |
| **Richness** | **Order** | **La Gomera *F. c. canariensis*** | **27.577** | **0.136** | **2.602** | **0.0093** | **21.095** | **36.035** |
| **Richness** | **Order** | **Tenerife *F. teydea*** | **35.333** | **0.185** | **3.268** | **0.0011** | **24.765** | **51.122** |
| **Richness** | **Order** | **Gran Canaria *F. polatzeki*** | **28.278** | **0.15** | **2.54** | **0.0111** | **21.109** | **37.986** |

*Continued on next page*

*Table 4 continued from previous page*

| Alpha diversity metric | Taxonomic level | Predictor | Estimate | Std. error | $z$ | Pr(> $\|z\|$) | CI 95% lo | CI 95% hi |
|---|---|---|---|---|---|---|---|---|
| **Faith's PD** | **Order** | **(Intercept)** | **6.583** | **0.098** | **19.247** | **<0.0001** | **5.422** | **7.962** |
| Faith's PD | Order | São Miguel *F. moreletti* | 5.714 | 0.136 | -1.037 | 0.2996 | 4.372 | 7.467 |
| Faith's PD | Order | Flores *F. moreletti* | 7 | 0.134 | 0.456 | 0.6482 | 5.379 | 9.115 |
| Faith's PD | Order | Madeira *F. maderensis* | 6 | 0.146 | -0.637 | 0.5244 | 4.506 | 7.981 |
| Faith's PD | Order | El Hierro *F. c. ombriosa* | 7.261 | 0.138 | 0.711 | 0.477 | 5.543 | 9.514 |
| Faith's PD | Order | La Palma *F. c. palmae* | 7.267 | 0.155 | 0.638 | 0.5236 | 5.36 | 9.838 |
| **Faith's PD** | **Order** | **Gran Canaria *F. c. bakeri*** | **9.857** | **0.127** | **3.191** | **0.0014** | **7.699** | **12.644** |
| Faith's PD | Order | Tenerife *F. c. canariensis* | 8 | 0.149 | 1.306 | 0.1917 | 5.968 | 10.719 |
| **Faith's PD** | **Order** | **La Gomera *F. c. canariensis*** | **9.5** | **0.129** | **2.843** | **0.0045** | **7.383** | **12.245** |
| **Faith's PD** | **Order** | **Tenerife *F. teydea*** | **11.444** | **0.167** | **3.306** | **0.0009** | **8.245** | **15.893** |
| **Faith's PD** | **Order** | **Gran Canaria *F. polatzeki*** | **9.722** | **0.14** | **2.782** | **0.0054** | **7.39** | **12.804** |
| **Richness** | **Family** | **(Intercept)** | **31.792** | **0.106** | **32.679** | **<0.0001** | **25.972** | **39.347** |
| **Richness** | **Family** | **São Miguel *F. moreletti*** | **22.214** | **0.146** | **-2.457** | **0.014** | **16.67** | **29.555** |
| Richness | Family | Flores *F. moreletti* | 31.731 | 0.147 | -0.013 | 0.9896 | 23.779 | 42.306 |
| Richness | Family | Madeira *F. maderensis* | 28 | 0.156 | -0.816 | 0.4144 | 20.647 | 38.029 |
| Richness | Family | El Hierro *F. c. ombriosa* | 35.87 | 0.151 | 0.8 | 0.4236 | 26.687 | 48.236 |
| Richness | Family | La Palma *F. c. palmae* | 33.8 | 0.17 | 0.36 | 0.7191 | 24.274 | 47.379 |
| **Richness** | **Family** | **Gran Canaria *F. c. bakeri*** | **47.393** | **0.143** | **2.793** | **0.0052** | **35.771** | **62.693** |
| Richness | Family | Tenerife *F. c. canariensis* | 38.937 | 0.166 | 1.219 | 0.2227 | 28.172 | 54.122 |
| **Richness** | **Family** | **La Gomera *F. c. canariensis*** | **44.423** | **0.146** | **2.297** | **0.0216** | **33.368** | **59.093** |
| **Richness** | **Family** | **Tenerife *F. teydea*** | **57.111** | **0.199** | **2.946** | **0.0032** | **39.014** | **85.242** |
| **Richness** | **Family** | **Gran Canaria *F. polatzeki*** | **45.944** | **0.16** | **2.301** | **0.0214** | **33.624** | **63.021** |
| **Faith's PD** | **Family** | **(Intercept)** | **8.042** | **0.099** | **21.046** | **<0.0001** | **6.619** | **9.763** |
| Faith's PD | Family | São Miguel *F. moreletti* | 6.607 | 0.139 | -1.419 | 0.156 | 5.034 | 8.667 |
| Faith's PD | Family | Flores *F. moreletti* | 8.269 | 0.137 | 0.204 | 0.8385 | 6.323 | 10.816 |
| Faith's PD | Family | Madeira *F. maderensis* | 7.19 | 0.147 | -0.759 | 0.4478 | 5.384 | 9.597 |
| Faith's PD | Family | El Hierro *F. c. ombriosa* | 8.957 | 0.14 | 0.772 | 0.4403 | 6.813 | 11.779 |
| Faith's PD | Family | La Palma *F. c. palmae* | 8.8 | 0.157 | 0.572 | 0.5671 | 6.463 | 11.985 |
| **Faith's PD** | **Family** | **Gran Canaria *F. c. bakeri*** | **12.071** | **0.129** | **3.14** | **0.0017** | **9.371** | **15.564** |

*Continued on next page*

*Table 4 continued from previous page*

| Alpha diversity metric | Taxonomic level | Predictor | Estimate | Std. error | $z$ | Pr($>\|z\|$) | CI 95% lo | CI 95% hi |
|---|---|---|---|---|---|---|---|---|
| Faith's PD | Family | Tenerife *F. c. canariensis* | 10.063 | 0.152 | 1.479 | 0.1391 | 7.479 | 13.55 |
| **Faith's PD** | **Family** | **La Gomera *F. c. canariensis*** | **11.5** | **0.132** | **2.709** | **0.0067** | **8.881** | **14.904** |
| **Faith's PD** | **Family** | **Tenerife *F. teydea*** | **14.222** | **0.173** | **3.293** | **0.001** | **10.148** | **20.015** |
| **Faith's PD** | **Family** | **Gran Canaria *F. polatzeki*** | **12.111** | **0.143** | **2.855** | **0.0043** | **9.148** | **16.056** |
| **Richness** | **Genus** | **(Intercept)** | **56.25** | **0.122** | **33.09** | **<0.0001** | **44.671** | **72.055** |
| **Richness** | **Genus** | **São Miguel *F. moreletti*** | **38.214** | **0.167** | **-2.317** | **0.0205** | **27.504** | **52.965** |
| Richness | Genus | Flores *F. moreletti* | 51.385 | 0.169 | -0.535 | 0.5926 | 36.842 | 71.572 |
| Richness | Genus | Madeira *F. maderensis* | 48.905 | 0.179 | -0.783 | 0.4334 | 34.475 | 69.544 |
| Richness | Genus | El Hierro *F. c. ombriosa* | 63.783 | 0.174 | 0.723 | 0.4697 | 45.352 | 89.772 |
| Richness | Genus | La Palma *F. c. palmae* | 53.133 | 0.197 | -0.29 | 0.7718 | 36.295 | 78.588 |
| **Richness** | **Genus** | **Gran Canaria *F. c. bakeri*** | **82.25** | **0.165** | **2.298** | **0.0216** | **59.371** | **113.665** |
| Richness | Genus | Tenerife *F. c. canariensis* | 67.75 | 0.192 | 0.969 | 0.3328 | 46.652 | 99.244 |
| **Richness** | **Genus** | **La Gomera *F. c. canariensis*** | **80.808** | **0.168** | **2.153** | **0.0313** | **58.03** | **112.378** |
| **Richness** | **Genus** | **Tenerife *F. teydea*** | **106.444** | **0.231** | **2.759** | **0.0058** | **68.525** | **170.253** |
| **Richness** | **Genus** | **Gran Canaria *F. polatzeki*** | **82.333** | **0.185** | **2.057** | **0.0397** | **57.388** | **118.809** |
| **Faith's PD** | **Genus** | **(Intercept)** | **9.958** | **0.101** | **22.71** | **<0.0001** | **8.175** | **12.16** |
| Faith's PD | Genus | São Miguel *F. moreletti* | 7.714 | 0.142 | -1.803 | 0.0715 | 5.841 | 10.18 |
| Faith's PD | Genus | Flores *F. moreletti* | 9.692 | 0.141 | -0.192 | 0.8474 | 7.354 | 12.771 |
| Faith's PD | Genus | Madeira *F. maderensis* | 9.048 | 0.15 | -0.64 | 0.5219 | 6.745 | 12.136 |
| Faith's PD | Genus | El Hierro *F. c. ombriosa* | 11.348 | 0.143 | 0.915 | 0.3604 | 8.578 | 15.018 |
| Faith's PD | Genus | La Palma *F. c. palmae* | 10.4 | 0.162 | 0.267 | 0.7892 | 7.571 | 14.31 |
| **Faith's PD** | **Genus** | **Gran Canaria *F. c. bakeri*** | **15.286** | **0.133** | **3.214** | **0.0013** | **11.77** | **19.854** |
| Faith's PD | Genus | Tenerife *F. c. canariensis* | 12.25 | 0.156 | 1.325 | 0.1852 | 9.025 | 16.661 |
| **Faith's PD** | **Genus** | **La Gomera *F. c. canariensis*** | **14.538** | **0.136** | **2.783** | **0.0054** | **11.138** | **18.983** |
| **Faith's PD** | **Genus** | **Tenerife *F. teydea*** | **18.222** | **0.18** | **3.352** | **0.0008** | **12.845** | **26.061** |
| **Faith's PD** | **Genus** | **Gran Canaria *F. polatzeki*** | **14.944** | **0.148** | **2.734** | **0.0062** | **11.18** | **20.013** |
| **Richness** | **Species** | **(Intercept)** | **64.667** | **0.124** | **33.73** | **<0.0001** | **51.192** | **83.168** |
| **Richness** | **Species** | **São Miguel *F. moreletti*** | **43.964** | **0.169** | **-2.28** | **0.0226** | **31.494** | **61.216** |
| Richness | Species | Flores *F. moreletti* | 57.923 | 0.172 | -0.642 | 0.521 | 41.321 | 81.083 |

*Continued on next page*

*Table 4 continued from previous page*

| Alpha diversity metric | Taxonomic level | Predictor | Estimate | Std. error | $z$ | $Pr(>\|z\|)$ | CI 95% lo | CI 95% hi |
|---|---|---|---|---|---|---|---|---|
| Richness | Species | Madeira *F. maderensis* | 56.524 | 0.181 | -0.743 | 0.4577 | 39.643 | 80.8 |
| Richness | Species | El Hierro *F. c. ombriosa* | 76 | 0.176 | 0.915 | 0.36 | 53.764 | 107.518 |
| Richness | Species | La Palma *F. c. palmae* | 58.067 | 0.2 | -0.539 | 0.5897 | 39.435 | 86.428 |
| **Richness** | **Species** | **Gran Canaria *F. c. bakeri*** | **96.286** | **0.168** | **2.371** | **0.0177** | **69.149** | **133.726** |
| Richness | Species | Tenerife *F. c. canariensis* | 78.375 | 0.195 | 0.986 | 0.3242 | 53.665 | 115.503 |
| **Richness** | **Species** | **La Gomera *F. c. canariensis*** | **95.577** | **0.171** | **2.287** | **0.0222** | **68.284** | **133.594** |
| **Richness** | **Species** | **Tenerife *F. teydea*** | **124.889** | **0.235** | **2.802** | **0.0051** | **79.855** | **201.38** |
| **Richness** | **Species** | **Gran Canaria *F. polatzeki*** | **97.167** | **0.188** | **2.165** | **0.0304** | **67.352** | **141.031** |
| **Faith's PD** | **Species** | **(Intercept)** | **10.083** | **0.102** | **22.571** | **<0.0001** | **8.26** | **12.344** |
| Faith's PD | Species | São Miguel *F. moreletti* | 7.964 | 0.143 | -1.651 | 0.0987 | 6.016 | 10.535 |
| Faith's PD | Species | Flores *F. moreletti* | 9.923 | 0.142 | -0.113 | 0.9103 | 7.508 | 13.113 |
| Faith's PD | Species | Madeira *F. maderensis* | 9.333 | 0.151 | -0.511 | 0.6091 | 6.94 | 12.554 |
| Faith's PD | Species | El Hierro *F. c. ombriosa* | 11.783 | 0.144 | 1.08 | 0.2802 | 8.882 | 15.637 |
| Faith's PD | Species | La Palma *F. c. palmae* | 10.533 | 0.164 | 0.266 | 0.7904 | 7.64 | 14.55 |
| **Faith's PD** | **Species** | **Gran Canaria *F. c. bakeri*** | **15.821** | **0.135** | **3.341** | **0.0008** | **12.146** | **20.611** |
| Faith's PD | Species | Tenerife *F. c. canariensis* | 12.437 | 0.158 | 1.326 | 0.1848 | 9.129 | 16.981 |
| **Faith's PD** | **Species** | **La Gomera *F. c. canariensis*** | **15.115** | **0.137** | **2.946** | **0.0032** | **11.546** | **19.794** |
| **Faith's PD** | **Species** | **Tenerife *F. teydea*** | **18.667** | **0.183** | **3.371** | **0.0007** | **13.101** | **26.833** |
| **Faith's PD** | **Species** | **Gran Canaria *F. polatzeki*** | **15.389** | **0.15** | **2.815** | **0.0049** | **11.474** | **20.681** |
| **Richness** | **OTU99** | **(Intercept)** | **109.625** | **0.139** | **33.833** | **<0.0001** | **84.461** | **145.707** |
| Richness | OTU99 | São Miguel *F. moreletti* | 82.429 | 0.189 | -1.505 | 0.1324 | 56.705 | 119.415 |
| Richness | OTU99 | Flores *F. moreletti* | 112.962 | 0.192 | 0.156 | 0.8762 | 77.307 | 164.758 |
| Richness | OTU99 | Madeira *F. maderensis* | 121.476 | 0.203 | 0.505 | 0.6133 | 81.619 | 181.415 |
| Richness | OTU99 | El Hierro *F. c. ombriosa* | 157.565 | 0.198 | 1.831 | 0.0671 | 106.785 | 232.735 |
| Richness | OTU99 | La Palma *F. c. palmae* | 86.933 | 0.224 | -1.034 | 0.3009 | 56.325 | 136.103 |
| **Richness** | **OTU99** | **Gran Canaria *F. c. bakeri*** | **168.714** | **0.189** | **2.282** | **0.0225** | **116.195** | **244.138** |
| Richness | OTU99 | Tenerife *F. c. canariensis* | 125.312 | 0.219 | 0.61 | 0.542 | 81.882 | 194.085 |
| **Richness** | **OTU99** | **La Gomera *F. c. canariensis*** | **184.269** | **0.192** | **2.703** | **0.0069** | **126.191** | **268.582** |
| **Richness** | **OTU99** | **Tenerife *F. teydea*** | **256.889** | **0.265** | **3.217** | **0.0013** | **155.523** | **441.704** |

*Continued on next page*

*Table 4 continued from previous page*

| Alpha diversity metric | Taxonomic level | Predictor | Estimate | Std. error | $z$ | Pr(> $\|z\|$) | CI 95% lo | CI 95% hi |
|---|---|---|---|---|---|---|---|---|
| **Richness** | **OTU99** | **Gran Canaria *F. polatzeki*** | **192.389** | **0.212** | **2.659** | **0.0078** | **127.414** | **292.828** |
| **Faith's PD** | **OTU99** | **(Intercept)** | **11.375** | **0.102** | **23.831** | **<0.0001** | **9.33** | **13.923** |
| Faith's PD | OTU99 | São Miguel *F. moreletti* | 9.321 | 0.142 | -1.407 | 0.1594 | 7.06 | 12.297 |
| Faith's PD | OTU99 | Flores *F. moreletti* | 11.923 | 0.141 | 0.334 | 0.7385 | 9.043 | 15.717 |
| Faith's PD | OTU99 | Madeira *F. maderensis* | 11.429 | 0.149 | 0.031 | 0.9749 | 8.53 | 15.32 |
| Faith's PD | OTU99 | El Hierro *F. c. ombriosa* | 13.826 | 0.144 | 1.36 | 0.1739 | 10.437 | 18.324 |
| Faith's PD | OTU99 | La Palma *F. c. palmae* | 11.533 | 0.164 | 0.084 | 0.9329 | 8.367 | 15.937 |
| **Faith's PD** | **OTU99** | **Gran Canaria *F. c. bakeri*** | **17.929** | **0.135** | **3.374** | **0.0007** | **13.761** | **23.354** |
| Faith's PD | OTU99 | Tenerife *F. c. canariensis* | 13.625 | 0.158 | 1.139 | 0.2548 | 9.997 | 18.616 |
| **Faith's PD** | **OTU99** | **La Gomera *F. c. canariensis*** | **17.077** | **0.137** | **2.956** | **0.0031** | **13.043** | **22.361** |
| **Faith's PD** | **OTU99** | **Tenerife *F. teydea*** | **21.778** | **0.183** | **3.548** | **0.0004** | **15.283** | **31.351** |
| **Faith's PD** | **OTU99** | **Gran Canaria *F. polatzeki*** | **17.778** | **0.15** | **2.975** | **0.0029** | **13.259** | **23.89** |

Table S5: **GLMM results testing predictions of Island Biogeography Theory.** GLMMs with a negative binomial distribution were fitted with microbiome richness as response variable and island area, distance to the mainland, and island age as predictors at each taxonomic level. Significant terms ($p < 0.05$) shown in bold.

| Taxonomic level | Predictor | Estimate | Std. error | $z$ | Pr(> \|z\|) |
|---|---|---|---|---|---|
| **Phylum** | **Intercept** | **2.1106** | **0.0476** | **44.3** | **<0.0001** |
| **Phylum** | **Island area** | **0.1017** | **0.0447** | **2.28** | **0.0228** |
| **Phylum** | **Intercept** | **2.1145** | **0.048** | **44.03** | **<0.0001** |
| **Phylum** | **Distance to mainland** | **-0.104** | **0.0508** | **-2.05** | **0.0406** |
| **Phylum** | **Intercept** | **2.1118** | **0.0321** | **65.86** | **<0.0001** |
| **Phylum** | **Island age** | **0.144** | **0.0323** | **4.46** | **<0.0001** |
| **Class** | **Intercept** | **2.4776** | **0.0621** | **39.87** | **<0.0001** |
| Class | Island area | 0.1071 | 0.0583 | 1.84 | 0.0659 |
| **Class** | **Intercept** | **2.4786** | **0.0567** | **43.73** | **<0.0001** |
| **Class** | **Distance to mainland** | **-0.1391** | **0.0598** | **-2.33** | **0.02** |
| **Class** | **Intercept** | **2.4794** | **0.0425** | **58.27** | **<0.0001** |
| **Class** | **Island age** | **0.1748** | **0.0429** | **4.07** | **<0.0001** |
| **Order** | **Intercept** | **3.132** | **0.0636** | **49.26** | **<0.0001** |
| **Order** | **Island area** | **0.1226** | **0.0601** | **2.04** | **0.0413** |
| **Order** | **Intercept** | **3.1323** | **0.0553** | **56.61** | **<0.0001** |
| **Order** | **Distance to mainland** | **-0.1616** | **0.0578** | **-2.8** | **0.0052** |

*Continued on next page*

*Table 5 continued from previous page*

| Taxonomic level | Predictor | Estimate | Std. error | $z$ | Pr($>\|z\|$) |
|---|---|---|---|---|---|
| **Order** | **Intercept** | **3.1339** | **0.0442** | **70.82** | **<0.0001** |
| **Order** | **Island age** | **0.1839** | **0.0445** | **4.13** | **<0.0001** |
| **Family** | **Intercept** | **3.5948** | **0.0718** | **50.03** | **<0.0001** |
| **Family** | **Island area** | **0.1345** | **0.0678** | **1.98** | **0.0472** |
| **Family** | **Intercept** | **3.5939** | **0.0601** | **59.83** | **<0.0001** |
| **Family** | **Distance to mainland** | **-0.1892** | **0.0627** | **-3.02** | **0.0025** |
| **Family** | **Intercept** | **3.5974** | **0.0512** | **70.21** | **<0.0001** |
| **Family** | **Island age** | **0.2059** | **0.0517** | **3.98** | **<0.0001** |
| **Genus** | **Intercept** | **4.1455** | **0.0786** | **52.77** | **<0.0001** |
| **Genus** | **Island area** | **0.1506** | **0.0744** | **2.03** | **0.0428** |
| **Genus** | **Intercept** | **4.1444** | **0.0624** | **66.43** | **<0.0001** |
| **Genus** | **Distance to mainland** | **-0.2142** | **0.0644** | **-3.33** | **0.0009** |
| **Genus** | **Intercept** | **4.1473** | **0.0539** | **76.96** | **<0.0001** |
| **Genus** | **Island age** | **0.2312** | **0.0543** | **4.26** | **<0.0001** |
| **Species** | **Intercept** | **4.2922** | **0.0828** | **51.86** | **<0.0001** |
| **Species** | **Island area** | **0.1557** | **0.0782** | **1.99** | **0.0465** |
| **Species** | **Intercept** | **4.2913** | **0.0646** | **66.44** | **<0.0001** |
| **Species** | **Distance to mainland** | **-0.2264** | **0.0667** | **-3.39** | **0.0007** |

*Continued on next page*

*Table 5 continued from previous page*

| Taxonomic level | Predictor | Estimate | Std. error | $z$ | $\Pr(>\|z\|)$ |
|---|---|---|---|---|---|
| **Species** | **Intercept** | **4.295** | **0.0569** | **75.51** | **<0.0001** |
| **Species** | **Island age** | **0.2429** | **0.0574** | **4.23** | **<0.0001** |
| **OTU99** | **Intercept** | **4.9287** | **0.0968** | **50.9** | **<0.0001** |
| OTU99 | Island area | 0.1282 | 0.0913 | 1.4 | 0.1603 |
| **OTU99** | **Intercept** | **4.9272** | **0.0788** | **62.51** | **<0.0001** |
| **OTU99** | **Distance to mainland** | **-0.2098** | **0.0817** | **-2.57** | **0.0103** |
| **OTU99** | **Intercept** | **4.9307** | **0.0712** | **69.3** | **<0.0001** |
| **OTU99** | **Island age** | **0.2383** | **0.0722** | **3.3** | **0.001** |

Table S6: **Mantel test results examining isolation-by-distance effects on microbiome dissimilarity among populations.** All tests are based on 9999 permutations; all $p$-values are $< 0.05$.

| Taxonomic level | Distance metric | Mantel $r$ | $p$-value |
|---|---|---|---|
| Phylum | Bray-Curtis | 0.1656 | 0.0001 |
| Phylum | Jaccard | 0.1029 | 0.0004 |
| Phylum | Unweighted UniFrac | 0.1408 | 0.0001 |
| Phylum | Weighted UniFrac | 0.1462 | 0.0002 |
| Class | Bray-Curtis | 0.1415 | 0.0001 |
| Class | Jaccard | 0.0777 | 0.0184 |
| Class | Unweighted UniFrac | 0.1121 | 0.0012 |
| Class | Weighted UniFrac | 0.126 | 0.0006 |
| Order | Bray-Curtis | 0.1208 | 0.0004 |
| Order | Jaccard | 0.1777 | 0.0002 |
| Order | Unweighted UniFrac | 0.1873 | 0.0001 |
| Order | Weighted UniFrac | 0.1009 | 0.0048 |
| Family | Bray-Curtis | 0.181 | 0.0001 |
| Family | Jaccard | 0.2502 | 0.0001 |
| Family | Unweighted UniFrac | 0.2399 | 0.0001 |
| Family | Weighted UniFrac | 0.1292 | 0.0005 |
| Genus | Bray-Curtis | 0.1647 | 0.0001 |
| Genus | Jaccard | 0.3507 | 0.0001 |
| Genus | Unweighted UniFrac | 0.2876 | 0.0001 |
| Genus | Weighted UniFrac | 0.1309 | 0.0008 |
| Species | Bray-Curtis | 0.1582 | 0.0001 |
| Species | Jaccard | 0.3662 | 0.0001 |
| Species | Unweighted UniFrac | 0.2896 | 0.0001 |
| Species | Weighted UniFrac | 0.1304 | 0.0011 |
| OTU99 | Bray-Curtis | 0.0626 | 0.0422 |
| OTU99 | Jaccard | 0.4289 | 0.0001 |
| OTU99 | Unweighted UniFrac | 0.3206 | 0.0001 |
| OTU99 | Weighted UniFrac | 0.1256 | 0.0008 |

Table S7: **Multiple regression on distance matrices (MRM) results.** Models test the relative contribution of host phylogeny, diet, and geography to microbiome dissimilarity (Bray-Curtis and Jaccard metrics) at each taxonomic level.

| Taxonomic level | Distance | Predictor | Estimate | $p$-value | Indiv. $R^2$ | Model $R^2$ | $F$ | Model $p$ |
|---|---|---|---|---|---|---|---|---|
| Phylum | Bray-Curtis | Int | 0.4819 | 1 | | 0.0325 | 58.8 | 0.0003 |
| Phylum | Bray-Curtis | host_dist | 0.442 | 0.3409 | 0.0069 | 0.0325 | 58.8 | 0.0003 |
| Phylum | Bray-Curtis | diet_dist | 0.0356 | 0.1066 | 0.0149 | 0.0325 | 58.8 | 0.0003 |
| **Phylum** | **Bray-Curtis** | **lgeo_dist** | **0.0384** | **0.0002** | **0.0274** | **0.0325** | **58.8** | **0.0003** |
| Phylum | Jaccard | Int | 0.5056 | 1 | | 0.0227 | 40.57 | 0.0003 |
| **Phylum** | **Jaccard** | **host_dist** | **0.7112** | **0.0055** | **0.0161** | **0.0227** | **40.57** | **0.0003** |
| Phylum | Jaccard | diet_dist | 0.008 | 0.5207 | 0.0063 | 0.0227 | 40.57 | 0.0003 |
| **Phylum** | **Jaccard** | **lgeo_dist** | **0.0128** | **0.0167** | **0.0106** | **0.0227** | **40.57** | **0.0003** |
| Class | Bray-Curtis | Int | 0.544 | 1 | | 0.0293 | 52.75 | 0.0002 |
| **Class** | **Bray-Curtis** | **host_dist** | **0.9905** | **0.022** | **0.0147** | **0.0293** | **52.75** | **0.0002** |
| Class | Bray-Curtis | diet_dist | 0.013 | 0.5342 | 0.0086 | 0.0293 | 52.75 | 0.0002 |
| **Class** | **Bray-Curtis** | **lgeo_dist** | **0.0322** | **0.0006** | **0.02** | **0.0293** | **52.75** | **0.0002** |
| Class | Jaccard | Int | 0.5765 | 0.9977 | | 0.0453 | 83.04 | 0.0001 |
| **Class** | **Jaccard** | **host_dist** | **1.111** | **0.0002** | **0.0422** | **0.0453** | **83.04** | **0.0001** |
| Class | Jaccard | diet_dist | 0.0163 | 0.2376 | 0.0115 | 0.0453 | 83.04 | 0.0001 |
| Class | Jaccard | lgeo_dist | 0.0031 | 0.6065 | 0.006 | 0.0453 | 83.04 | 0.0001 |
| Order | Bray-Curtis | Int | 0.6744 | 1 | | 0.0198 | 35.33 | 0.0018 |
| Order | Bray-Curtis | host_dist | 0.6625 | 0.0776 | 0.0088 | 0.0198 | 35.33 | 0.0018 |
| Order | Bray-Curtis | diet_dist | 0.0076 | 0.6775 | 0.0055 | 0.0198 | 35.33 | 0.0018 |
| **Order** | **Bray-Curtis** | **lgeo_dist** | **0.0251** | **0.0028** | **0.0146** | **0.0198** | **35.33** | **0.0018** |
| Order | Jaccard | Int | 0.6611 | 1 | | 0.063 | 117.69 | 0.0001 |
| **Order** | **Jaccard** | **host_dist** | **0.6365** | **0.0094** | **0.0344** | **0.063** | **117.69** | **0.0001** |
| **Order** | **Jaccard** | **diet_dist** | **0.0277** | **0.0143** | **0.0334** | **0.063** | **117.69** | **0.0001** |
| **Order** | **Jaccard** | **lgeo_dist** | **0.0134** | **0.0079** | **0.0316** | **0.063** | **117.69** | **0.0001** |
| Family | Bray-Curtis | Int | 0.7215 | 1 | | 0.0649 | 121.47 | 0.0001 |
| **Family** | **Bray-Curtis** | **host_dist** | **1.4512** | **0.0001** | **0.0447** | **0.0649** | **121.47** | **0.0001** |
| Family | Bray-Curtis | diet_dist | 0.0094 | 0.488 | 0.0155 | 0.0649 | 121.47 | 0.0001 |

*Continued on next page*

*Table 7 continued from previous page*

| Taxonomic level | Distance | Predictor | Estimate | $p$-value | Indiv. $R^2$ | Model $R^2$ | $F$ | Model $p$ |
|---|---|---|---|---|---|---|---|---|
| **Family** | **Bray-Curtis** | **lgeo_dist** | **0.0291** | **0.0001** | **0.0328** | **0.0649** | **121.47** | **0.0001** |
| Family | Jaccard | Int | 0.6937 | 1 | | 0.078 | 148 | 0.0002 |
| Family | Jaccard | host_dist | 0.2119 | 0.4466 | 0.0138 | 0.078 | 148 | 0.0002 |
| **Family** | **Jaccard** | **diet_dist** | **0.0309** | **0.0176** | **0.0431** | **0.078** | **148** | **0.0002** |
| **Family** | **Jaccard** | **lgeo_dist** | **0.0231** | **0.0001** | **0.0626** | **0.078** | **148** | **0.0002** |
| Genus | Bray-Curtis | Int | 0.8104 | 1 | | 0.0528 | 97.53 | 0.0001 |
| **Genus** | **Bray-Curtis** | **host_dist** | **1.0782** | **0.0001** | **0.0353** | **0.0528** | **97.53** | **0.0001** |
| Genus | Bray-Curtis | diet_dist | 0.0131 | 0.3044 | 0.0153 | 0.0528 | 97.53 | 0.0001 |
| **Genus** | **Bray-Curtis** | **lgeo_dist** | **0.022** | **0.0009** | **0.0271** | **0.0528** | **97.53** | **0.0001** |
| Genus | Jaccard | Int | 0.7993 | 0.9962 | | 0.1371 | 277.91 | 0.0001 |
| Genus | Jaccard | host_dist | -0.0812 | 0.6763 | 0.0055 | 0.1371 | 277.91 | 0.0001 |
| **Genus** | **Jaccard** | **diet_dist** | **0.0266** | **0.0031** | **0.062** | **0.1371** | **277.91** | **0.0001** |
| **Genus** | **Jaccard** | **lgeo_dist** | **0.0271** | **0.0001** | **0.123** | **0.1371** | **277.91** | **0.0001** |
| Species | Bray-Curtis | Int | 0.8241 | 1 | | 0.0517 | 95.32 | 0.0001 |
| **Species** | **Bray-Curtis** | **host_dist** | **1.0894** | **0.0002** | **0.0362** | **0.0517** | **95.32** | **0.0001** |
| Species | Bray-Curtis | diet_dist | 0.0108 | 0.389 | 0.0139 | 0.0517 | 95.32 | 0.0001 |
| **Species** | **Bray-Curtis** | **lgeo_dist** | **0.0205** | **0.0009** | **0.025** | **0.0517** | **95.32** | **0.0001** |
| Species | Jaccard | Int | 0.8315 | 0.6375 | | 0.1504 | 309.73 | 0.0001 |
| Species | Jaccard | host_dist | -0.0557 | 0.7359 | 0.007 | 0.1504 | 309.73 | 0.0001 |
| **Species** | **Jaccard** | **diet_dist** | **0.0243** | **0.0013** | **0.0699** | **0.1504** | **309.73** | **0.0001** |
| **Species** | **Jaccard** | **lgeo_dist** | **0.0239** | **0.0001** | **0.1341** | **0.1504** | **309.73** | **0.0001** |
| OTU99 | Bray-Curtis | Int | 0.9017 | 0.9804 | | 0.0431 | 78.85 | 0.0001 |
| **OTU99** | **Bray-Curtis** | **host_dist** | **1.3133** | **0.0001** | **0.0426** | **0.0431** | **78.85** | **0.0001** |
| OTU99 | Bray-Curtis | diet_dist | -0.0046 | 0.7383 | 0.003 | 0.0431 | 78.85 | 0.0001 |
| OTU99 | Bray-Curtis | lgeo_dist | 0.0044 | 0.4651 | 0.0039 | 0.0431 | 78.85 | 0.0001 |
| **OTU99** | **Jaccard** | **Int** | **0.9492** | **0.0001** | | **0.2079** | **459.35** | **0.0001** |
| **OTU99** | **Jaccard** | **host_dist** | **0.0912** | **0.0112** | **0.0384** | **0.2079** | **459.35** | **0.0001** |
| **OTU99** | **Jaccard** | **diet_dist** | **0.0072** | **0.0001** | **0.0856** | **0.2079** | **459.35** | **0.0001** |
| **OTU99** | **Jaccard** | **lgeo_dist** | **0.0097** | **0.0001** | **0.1839** | **0.2079** | **459.35** | **0.0001** |

**Estimate**: regression coefficient ($\beta$) for that predictor. ***p*-value**: *p*-value for that specific predictor. **Indiv.** $R^2$: individual-predictor $R^2$ from single-predictor models. **Model** $R^2$: overall full-model $R^2$. $F$: overall model $F$-statistic. **Model** $p$: *p*-value for the overall $F$-test. Predictors: `host_dist` = mitogenome cophenetic distance; `diet_dist` = Raup-Crick dissimilarity on diet ASVs; `lgeo_dist` = log-transformed pairwise geographic distance. Significant predictors ($p < 0.05$) shown in bold.

Table S8: **PACo (Procrustean Analysis of Cophylogeny) results for core microbial families.** Co-phylogenetic signal estimates at species- and OTU-level taxonomic clustering of microbial OTUs with Bonferroni corrected *p-values* of tests using host and random trees. Significant PACo tests ( Bonferroni-adjusted $p < 0.05$ against the host tree *and* non-significant ($p > 0.05$) against random trees) shown in bold .

| Taxonomic clustering | Test | $R^2$ | SS | $p$-value | Host tree adj. $p$ | Random tree adj. $p$ |
|---|---|---|---|---|---|---|
| Species | 70_9 | 0.1374 | 0.9811 | 0.643 | 1 | 1 |
| Species | Acetobacteraceae | 0.1696 | 0.9712 | 0.149 | 1 | 1 |
| Species | Beijerinckiaceae | 0.1134 | 0.9871 | 0.266 | 1 | 1 |
| **Species** | **Burkholderiaceae** | **0.4523** | **0.7954** | **0.001** | **0.027** | **1** |
| Species | Burkholderiaceae_B | 0.249 | 0.938 | 0.038 | 1 | 1 |
| Species | Cellulomonadaceae | 0.1983 | 0.9607 | 0.336 | 1 | 1 |
| Species | Dermatophilaceae | 0.3045 | 0.9073 | 0.021 | 0.567 | 1 |
| Species | Enterobacteriaceae | 0.2346 | 0.945 | 0.019 | 0.513 | 1 |
| Species | Ilumatobacteraceae | 0.2298 | 0.9472 | 0.121 | 1 | 1 |
| Species | Isosphaeraceae | 0.1175 | 0.9862 | 0.981 | 1 | 1 |
| **Species** | **Lactobacillaceae** | **0.2989** | **0.9107** | **0.001** | **0.027** | **1** |
| **Species** | **Microbacteriaceae** | **0.1533** | **0.9765** | **0.001** | **0.027** | **1** |
| Species | Micrococcaceae | 0.2321 | 0.9462 | 0.041 | 1 | 1 |
| Species | Micromonosporaceae | 0.2057 | 0.9577 | 0.501 | 1 | 1 |
| Species | Mycobacteriaceae | 0.2056 | 0.9577 | 0.003 | 0.081 | 1 |
| Species | Nakamurellaceae | 0.1322 | 0.9825 | 0.915 | 1 | 1 |
| Species | Nocardioidaceae | 0.1586 | 0.9748 | 0.16 | 1 | 1 |
| Species | Polyangiaceae | 0.2373 | 0.9437 | 0.285 | 1 | 1 |
| Species | Propionibacteriaceae | 0.1224 | 0.985 | 0.62 | 1 | 1 |
| Species | Pseudomonadaceae | 0.2245 | 0.9496 | 0.387 | 1 | 1 |
| Species | Pseudonocardiaceae | 0.1936 | 0.9625 | 0.043 | 1 | 1 |

*Continued on next page*

*Table 8 continued from previous page*

| Taxonomic clustering | Test | $R^2$ | SS | $p$-value | Host tree adj. $p$ | Random tree adj. $p$ |
|---|---|---|---|---|---|---|
| Species | Rhizobiaceae | 0.0966 | 0.9907 | 0.407 | 1 | 1 |
| Species | Solirubrobacteraceae | 0.1426 | 0.9797 | 0.186 | 1 | 1 |
| Species | Sphingomonadaceae | 0.1735 | 0.9699 | 0.471 | 1 | 1 |
| Species | Staphylococcaceae | 0.2681 | 0.9281 | 0.253 | 1 | 1 |
| Species | Streptomycetaceae | 0.212 | 0.9551 | 0.811 | 1 | 1 |
| Species | Xanthobacteraceae | 0.2181 | 0.9524 | 0.005 | 0.135 | 1 |
| OTU99 | 70_9 | 0.2093 | 0.9562 | 0.21 | 1 | 1 |
| OTU99 | Acetobacteraceae | 0.188 | 0.9647 | 0.001 | 0.03 | 0.03 |
| OTU99 | Beijerinckiaceae | 0.1293 | 0.9833 | 0.001 | 0.03 | 0.03 |
| OTU99 | Burkholderiaceae | 0.5026 | 0.7474 | 0.001 | 0.03 | 1 |
| OTU99 | Burkholderiaceae_B | 0.2431 | 0.9409 | 0.021 | 0.63 | 1 |
| OTU99 | Cellulomonadaceae | 0.2882 | 0.9169 | 0.004 | 0.12 | 0.9 |
| OTU99 | Dermatophilaceae | 0.3204 | 0.8973 | 0.001 | 0.03 | 0.45 |
| OTU99 | Devosiaceae | 0.4093 | 0.8325 | 0.022 | 0.66 | 1 |
| OTU99 | Enterobacteriaceae | 0.2444 | 0.9403 | 0.001 | 0.03 | 0.03 |
| OTU99 | Ilumatobacteraceae | 0.2873 | 0.9175 | 0.036 | 1 | 1 |
| OTU99 | Isosphaeraceae | 0.2374 | 0.9436 | 0.001 | 0.03 | 0.03 |
| OTU99 | Jatrophihabitantaceae | 0.2519 | 0.9365 | 0.001 | 0.03 | 1 |
| OTU99 | Kineococcaceae | 0.1897 | 0.964 | 0.596 | 1 | 1 |
| OTU99 | Lactobacillaceae | 0.2002 | 0.9599 | 0.001 | 0.03 | 0.21 |
| OTU99 | Microbacteriaceae | 0.1876 | 0.9648 | 0.001 | 0.03 | 0.03 |
| OTU99 | Micrococcaceae | 0.2594 | 0.9327 | 0.001 | 0.03 | 0.03 |
| OTU99 | Micromonosporaceae | 0.2612 | 0.9318 | 0.132 | 1 | 1 |
| OTU99 | Mycobacteriaceae | 0.2024 | 0.959 | 0.001 | 0.03 | 0.09 |
| OTU99 | Nakamurellaceae | 0.2016 | 0.9593 | 0.592 | 1 | 1 |

*Continued on next page*

*Table 8 continued from previous page*

| Taxonomic clustering | Test | $R^2$ | SS | $p$-value | Host tree adj. $p$ | Random tree adj. $p$ |
|---|---|---|---|---|---|---|
| OTU99 | Nocardioidaceae | 0.1978 | 0.9609 | 0.001 | 0.03 | 1 |
| OTU99 | Polyangiaceae | 0.3949 | 0.8441 | 0.001 | 0.03 | 1 |
| OTU99 | Propionibacteriaceae | 0.1848 | 0.9658 | 0.002 | 0.06 | 1 |
| OTU99 | Pseudomonadaceae | 0.2592 | 0.9328 | 0.56 | 1 | 1 |
| OTU99 | Pseudonocardiaceae | 0.1933 | 0.9626 | 0.001 | 0.03 | 1 |
| OTU99 | Rhizobiaceae | 0.2004 | 0.9598 | 0.001 | 0.03 | 1 |
| OTU99 | Solirubrobacteraceae | 0.1931 | 0.9627 | 0.001 | 0.03 | 0.03 |
| OTU99 | Sphingomonadaceae | 0.1755 | 0.9692 | 0.003 | 0.09 | 1 |
| OTU99 | Staphylococcaceae | 0.3607 | 0.8699 | 0.013 | 0.39 | 1 |
| OTU99 | Streptomycetaceae | 0.2346 | 0.945 | 0.338 | 1 | 1 |
| OTU99 | Xanthobacteraceae | 0.3123 | 0.9024 | 0.001 | 0.03 | 0.78 |

Table S9: **emPRess event-based reconciliation analysis output.** Summary of phylogenetic congruence results between host and core microbial lineages under five cost parameterisations (duplication / host transfer / loss). MPRs: number of maximum parsimony reconciliations. $p$-values are computed against 100 random host trees.

| | | **Cost value parameters** | | | | **Events** | | | | |
|---|---|---|---|---|---|---|---|---|---|---|
| Core symbiont family | Variant | Dup. | Host transfer | Loss | MPRs | Co-spec. | Duplications | Nr. Transfers | Nr. Losses | $p$ (100 trials) |
| Burkholderiaceae | OTU | 1 | 1 | 1 | > 10000 | 9 | 89 | 106 | 1 | 0.95 |
| Burkholderiaceae | OTU | 4 | 1 | 1 | > 10000 | 9 | 82 | 113 | 1 | 1 |
| Burkholderiaceae | OTU | 2 | 1 | 2 | > 10000 | 9 | 82 | 113 | 1 | 1 |
| Burkholderiaceae | OTU | 4 | 2 | 1 | > 10000 | 21 | 82 | 101 | 17 | 1 |
| Burkholderiaceae | OTU | 2 | 4 | 1 | 1920 | 17 | 134 | 53 | 24 | 0.009 |
| Burkholderiaceae | Species | 1 | 1 | 1 | > 10000 | 5 | 6 | 28 | 2 | 0.86 |
| Burkholderiaceae | Species | 4 | 1 | 1 | > 10000 | 5 | 5 | 29 | 2 | 0.98 |
| Burkholderiaceae | Species | 2 | 1 | 2 | 3072 | 3 | 5 | 31 | 0 | 0.97 |
| Burkholderiaceae | Species | 4 | 2 | 1 | 360 | 8 | 5 | 26 | 6 | 0.92 |
| Burkholderiaceae | Species | 2 | 4 | 1 | 20 | 9 | 13 | 17 | 11 | 0.13 |
| Lactobacillaceae | OTU | 1 | 1 | 1 | > 10000 | 100 | 308 | 460 | 3 | 1 |
| Lactobacillaceae | OTU | 4 | 1 | 1 | > 10000 | 90 | 247 | 531 | 3 | 1 |
| Lactobacillaceae | OTU | 2 | 1 | 2 | > 10000 | 89 | 250 | 529 | 0 | 1 |
| Lactobacillaceae | OTU | 4 | 2 | 1 | > 10000 | 103 | 249 | 516 | 18 | 1 |
| Lactobacillaceae | OTU | 2 | 4 | 1 | > 10000 | 151 | 569 | 148 | 286 | 0.07 |
| Lactobacillaceae | Species | 1 | 1 | 1 | > 10000 | 8 | 5 | 44 | 0 | 0.81 |
| Lactobacillaceae | Species | 4 | 1 | 1 | 1920 | 8 | 4 | 45 | 0 | 0.71 |
| Lactobacillaceae | Species | 2 | 1 | 2 | 1536 | 8 | 4 | 45 | 0 | 0.78 |
| Lactobacillaceae | Species | 4 | 2 | 1 | 846 | 13 | 4 | 40 | 7 | 0.72 |
| Lactobacillaceae | Species | 2 | 4 | 1 | 1470 | 17 | 21 | 19 | 31 | 0.48 |
| Microbacteriaceae | OTU | 1 | 1 | 1 | > 10000 | 65 | 54 | 374 | 20 | 0.99 |
| Microbacteriaceae | OTU | 4 | 1 | 1 | > 10000 | 67 | 44 | 382 | 12 | 1 |
| Microbacteriaceae | OTU | 2 | 1 | 2 | > 10000 | 52 | 44 | 397 | 0 | 1 |
| Microbacteriaceae | OTU | 4 | 2 | 1 | > 10000 | 104 | 44 | 345 | 57 | 1 |
| Microbacteriaceae | OTU | 2 | 4 | 1 | > 10000 | 137 | 133 | 223 | 213 | 0.05 |
| Microbacteriaceae | Species | 1 | 1 | 1 | > 10000 | 13 | 8 | 79 | 0 | 0.31 |

*Table 9 continued from previous page*

| | | **Cost value parameters** | | | | **Events** | | | | |
|---|---|---|---|---|---|---|---|---|---|---|
| Core symbiont family | Variant | Dup. | Host transfer | Loss | MPRs | Co-spec. | Duplications | Nr. Transfers | Nr. Losses | $p$ (100 trials) |
| Microbacteriaceae | Species | 4 | 1 | 1 | $> 10000$ | 13 | 7 | 80 | 0 | 0.84 |
| Microbacteriaceae | Species | 2 | 1 | 2 | $> 10000$ | 13 | 7 | 80 | 0 | 0.61 |
| Microbacteriaceae | Species | 4 | 2 | 1 | $> 10000$ | 20 | 7 | 73 | 9 | 0.83 |
| Microbacteriaceae | Species | 2 | 4 | 1 | $> 10000$ | 26 | 30 | 44 | 46 | 0.68 |
